# Di-Gluebodies as covalently-rigidified, modular protein assemblies enable simultaneous determination of high-resolution, low-size, cryo-EM structures

**DOI:** 10.1101/2024.06.13.598841

**Authors:** Gangshun Yi, Dimitrios Mamalis, Mingda Ye, Loic Carrique, Michael Fairhead, Huanyu Li, Katharina L. Duerr, Peijun Zhang, David B. Sauer, Frank von Delft, Benjamin G. Davis, Robert J. C. Gilbert

**Affiliations:** Division of Structural Biology, Centre for Human Genetics, University of Oxford; Roosevelt Drive, Oxford OX3 7BN, UK; Calleva Centre for Evolution and Human Science, Magdalen College; High Street, Oxford OX1 4AU, UK; Department of Chemistry, University of Oxford, OX1 3TA, Oxford, UK; The Rosalind Franklin Institute; Oxfordshire, OX11 0QX, UK; Centre for Medicines Discovery, University of Oxford; NDM Research Building, Roosevelt Drive, Oxford, OX3 7FZ, UK; Diamond Light Source, Harwell Science and Innovation Campus; Fermi Avenue, Didcot OX11 0DE, UK; Chinese Academy of Medical Sciences Oxford Institute, University of Oxford; Roosevelt Drive, Oxford OX3 7BN, UK; Research Complex at Harwell, Harwell Science and Innovation Campus; Fermi Avenue, Didcot, OX11 0DE, UK; Department of Biochemistry, University of Johannesburg, Auckland Park, Johannesburg 2006, South Africa; Department of Pharmacology, University of Oxford; Oxford, OX1 3QT, UK

## Abstract

Cryo-EM has become a routine structural biology method, yet elucidation of small proteins (<100 kDa) and increasing throughput remain challenging. Here, we describe covalently-dimerized engineered nanobodies, Di-Gluebodies, as novel and modular imaging scaffolds to address the challenges. They can simultaneously display two copies of the same protein or two distinct proteins, providing sufficient rigidity required by cryo-EM. We validate this method with multiple soluble and membrane targets, demonstrating that this provides a flexible and scalable platform for expanded protein structure determination.

## Introduction

Single-particle cryo-EM is a routine method for protein structure determination, and has developed rapidly since it was first shown useful for elucidating structures of large protein complexes^1^. Nonetheless, structural determination of smaller proteins (<100 kDa) by cryo-EM remains challenging^2^. Consequently, less than 3.5% of deposited structures in the Electron Microscopy Data Bank (EMDB) have a molecular weight below 100 kDa ^3^, despite the fact that small proteins are abundant in nature, with 92.3% protein-coding genes in humans generating products below 100 kDa and 74.5% below 50 kDa ^4^. Therefore, to determine protein structures not accessible by other methods or in conditions often closer to physiological^5^, new tools and methods are needed to facilitate cryo-EM of small targets.

A major challenge for small-protein cryo-EM is the low signal-to-noise ratio of the particles^2^. This leads to difficulty in particle picking and alignment during data processing, with these inaccuracies ultimately resulting in high B-factors and lower resolutions for 3D reconstructions^2^. Therefore, cryo-EM reconstruction generally remains easier for larger targets and complexes, as they provide sufficient signal for particle alignment. Taking advantage of this, methods for increasing particle size and symmetry with fiducial markers have been developed^4^. Three categories of toolsets are currently available: high-symmetry scaffolding particles^6^, and fusions with^4, 7^ or binding by additional protein structures ^8-10^. Such binding modules include antibody fragments (Fab), nanobodies (Nb), as well as synthetic backbones with evolved or selected binding sequences. These modules can further be fused or bound to other proteins for still greater fiducial mass, including the NabFab^11^, megabody^12^, Legobody^13^ and BRIL^14^ technologies (**Extended Data Figure 1**). BRIL, in particular, coupled with the engineering of rigidly bound epitopes into the target sequence, enables more generic Fab and nanobody targeting to multiple proteins^12, 14^, thereby partially avoiding the time-consuming processes of binder generation and selection.

Despite this demonstrated proof-of-principle, challenges with binder generation and binder:target complex assembly still present problems for such fiducial-assisted single-particle cryo-EM. Fusing tags or inserting epitopes requires modifying the target itself and thereby risks altering the protein’s native structure, potentially also reducing its expression, or requiring extensive screening of constructs^15^. In addition, most tools do not have symmetry and also require chimeric constructs and thus need subcloning and expression optimization^6, 11, 13^. Above all, these existing scaffolds are mono-specific, and consequently do not address generality or modularity and therefore the urgent need for greater sample throughput (Extended Data Figure 1). Therefore, there remains a need to develop a general strategy for fiducial optimization, ideally with the potential to multiplex structure determination. Here, we show that a generic and kinetically-biased ‘gluebody’ interface, previously discovered through trapping *in crystallo*,^16^ can also be trapped in solution by kinetically-controlled sidechain-to-sidechain bond-formation in a manner that leads to ready formation of complexes; these display ideally balanced interface flexibility and yet complex rigidity that in turn allows rapid cryo-EM structure determination, even of small targets.

## Results

### Homo Di-Gluebodies enable cryo-EM structural determination for small proteins

One ideal ‘plug-and-play’ design for this needed tool would be through the rigid complexation of two binding modules via a structurally robust interface away from their binding loops, in which case, modularity, symmetry, and bi-specificity are all achieved. Analysis of our previous crystallographic observations of protein modules, based on the widely-used nanobodies^16^, that self-assemble in the solid-phase (‘Gluebodies’) revealed a surprising and seemingly generic non-covalent *in crystallo* interface that we reasoned could be rendered stable through designed covalency. In this way, kinetically-controlled trapping *in crystallo* might parallel and so guide efficient kinetically-controlled trapping in solution.

In principle, many covalent bond-forming (‘conjugation’) methods are available to link protein modules. However, few allow the generation of a minimally-sized link. Further, this method must balance the needed proximity for our designed interfacial rigidity and yet a sufficiently efficient reaction under the inherently challenging second-order kinetics of protein dimerization under conditions of low concentration. The observed critical role of pivotal residues in driving ‘glue sites’ at protein-protein interfaces in the solid phase,^16^ led us to test these as promising sites for such chemistry. Ultimately, after a wide survey of diverse in-solution covalent bond-forming methods exploring differing natural and unnatural amino acid residues, we found that kinetically-controlled Cu(II)-catalyzed oxidation at a pivotal^16^ Cys12 residue site (together with the presence of needed Gb ‘glue’ mutations previously discovered through iterative X-ray-guided engineering^16^) offered a route to direct and rapid, high-yielding formation of disulfide-linked homodimeric Gluebodies (homoDiGbs) without side reactions (e.g. off-site oxidation, **Figure 1a-c** and **Extended Data Figure 2** and 3) to form a robust covalent interface.

**Figure 1.**
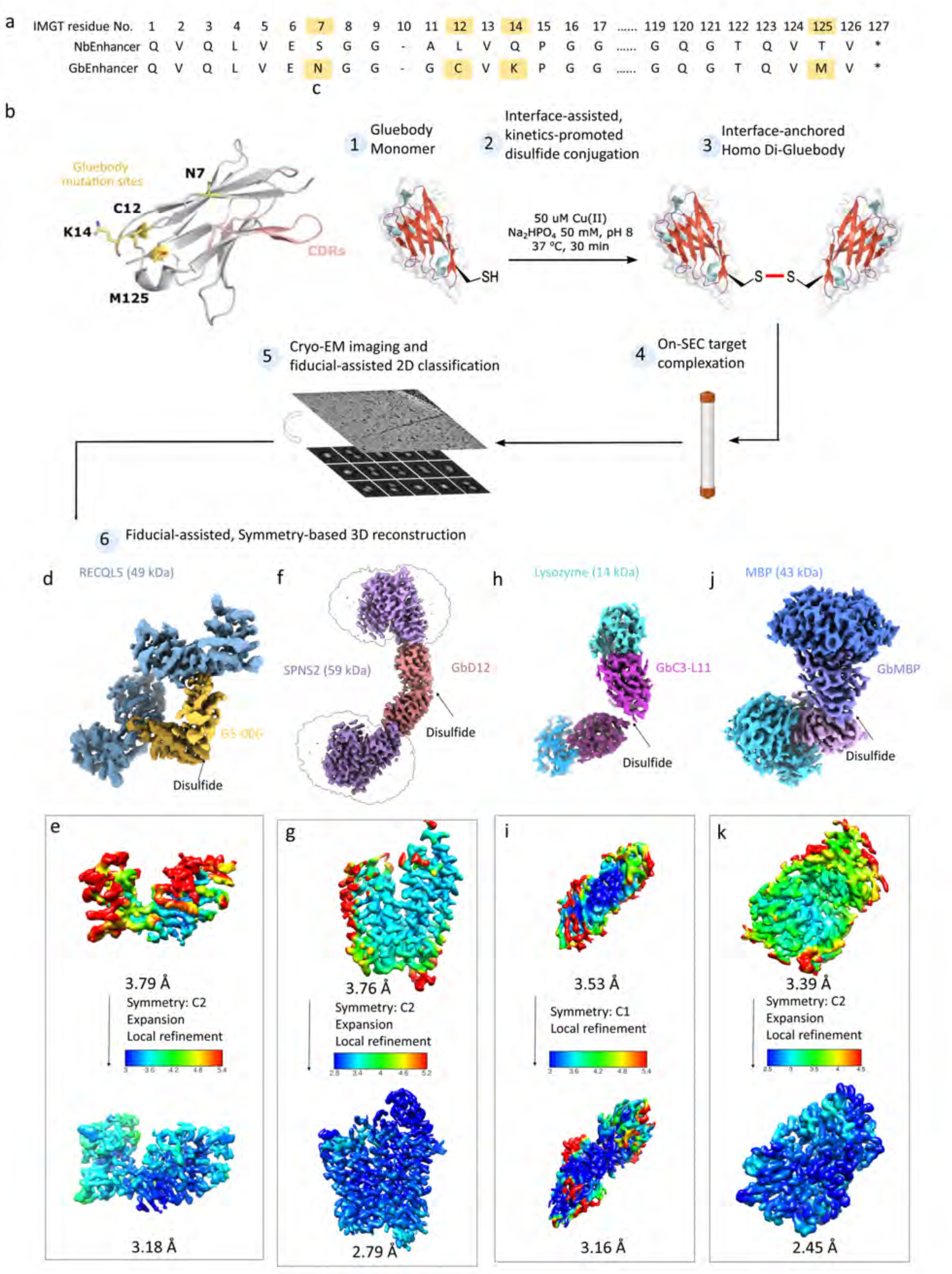
Application of homo Di-Gluebodies (homoDiGb) enables high-resolution cryo-EM structure determination for dual copies of small proteins. (a) Designed Gluebody (Gb) core mutations (S7N/L12C/Q14K/T125M in gold) on the nanobody(Nb) backbone enable chemically-driven, covalent Di-Gluebody generation. The sequences are aligned using the IMGT scheme developed for immunoglobulin folds^18^. The sequence difference between anti-GFP nanobody NbEnhancer (PDB 3K1K)^19^ and its Gluebody equivalent (Extended Data Figure 2) is shown as an example. (b) Schematic cartoon of the anti-GFP nanobody NbEnhancer showing the Gluebody mutation sites(gold) away from CDRs(pink). (c) Construction pipeline for homoDiGb. (d-k) High-resolution cryo-EM reconstruction maps of (d)RECQL5, (f)SPNS2, (h)lysozyme and (j)MBP in complex with their respective homoDiGbs. While also showing the local resolution comparisons before and after local refinement with or without C2 symmetry expansion for (e)RECQL5, (g)SPNS2, (i)lysozyme and (k)MBP

To probe the applicability and value of these homoDiGbs to cryo-EM structure determination, we tested the 49 kDa DNA helicase RECQL5 as a challenging soluble target with both small size and significant inter-domain flexibility.^17^ As an initial experiment, we compared the 2D cryo-EM classes of RECQL5 alone; RECQL5 bound to a wild-type nanobody; and RECQL5 in complex with the corresponding homoDiGb (homoDiGb5-006) generated from our chemically-driven, covalent-dimerization workflow (**Figure 1**). With the same particle-picking strategies, the homoDiGb resulted in many well-resolved classes for different views (**Extended Data Figure 6**), appreciably outperforming the other RECQL5 samples. Next, a larger dataset of this RECQL5:homoDiGb complex on a 300-kV microscope with standard operating parameters (**Extended Data Table 1**) yielded a 3.79 Å reconstruction of the complex after imposing C2 symmetry. Moreover, this initial map could be further improved by foregoing strict C2 symmetry, with symmetry expansion and local refinement of RECQL5 alone improving the resolution to 3.18 Å (**Figures 1d and 1e**, **Extended Data Figure 7**). Similarly, the clinically-relevant, membrane protein Sphingosine-1-phosphate (S1P) transporter Spinster Homolog 2 (SPNS2) was resolved to 2.79 Å locally, when bound to the equivalent homoDiGbD12 derived from the SPNS2-binding nanobody NbD12 ^10^ (**Figures 1f and 1g**, **Extended Data Figure 10**). Notably, the homoDiGb-bound structure of SPNS2 was found in the same inward-facing DDM-bound state as when bound to wild-type nanobody NbD12, indicating minimal perturbation to the structure. Comparison shows clear improvements over the previously published structures of SPNS2 bound to NbD12 (EMD-18668)^10^ (with significantly better defined sidechains, **Extended Data Figure 13b**) and, importantly, over structures determined with NabFab (EMD-34104), or DARPin and MBP fusions (EMD-28650).^3, 7^ This increased resolution usefully revealed interactions at the intracellular loops of SPNS2, including the important PI(4,5)P_2_ pocket (site 2, **Extended Data Figure 13**).^20^

The rapid implementation of the homoDiGb method and the additional insights gained into target structure function led us to consider the limits of this technology. Current cryo-EM structure determination methods typically struggle with targets below 50 kDa, contrasting with our ready solution of the structure of 49 kDa RECQL5. We therefore chose to probe even smaller targets. Strikingly, within weeks, using an essentially identical workflow to that used for RECQ5 and SPNS2, the homoDiGb method was applied to 43 kDa maltose binding protein (MBP) and even very small 14 kDa hen egg white lysozyme (**Extended Data Figures 3 and 4**), achieving reconstructions at 2.45 Å and 3.16 Å, respectively (**Figure 1h, 1i, 1j and 1k**, **Extended Data Figure 8** and 9). Interestingly, for lysozyme, an archetype of X-ray structure determination that we solve here by cryo-EM SPA for the first time, we found that C1 symmetry reconstruction followed by local refinement yielded better resolution than imposing C2 symmetry for the entire particle. This putative observation of nominal asymmetry for such extremely small proteins may offer strategic solutions for other very small targets as this field grows. Together, these data demonstrated that the homo Di-Gluebody approach provides a rapid, generic and efficient way for cryo-EM structural solution now even of small proteins.

### Heteromeric Di-Gluebodies can act as a ‘plug-and-play’ tool for simultaneous cryo-EM structural determination of small proteins

Next we explored bi-specificity of this tool, by testing the extension of our modular DiGb method to the more challenging generation of *heteromeric* Di-Gluebodies (heteroDiGbs). We reasoned that needed kinetic-control of covalency might be readily achieved via oxidative pre-activation of a Gluebody followed by the addition of a second Gluebody, to yield a homogeneous heterodimeric population via a trapped oxidation relay. Validating this notion, quantitative functionalization at site 12 of the first binder with 5,5-dithio-bis-(2-nitrobenzoic acid)^21^, produced a clean, trapped intermediate that was then readily reacted with a second Gluebody to form the desired heteromeric Di-Gluebodies (heteroDiGb) with high selectivities and yields (**Figure 2a**, **Extended Data Figure 4**).

**Figure 2.**
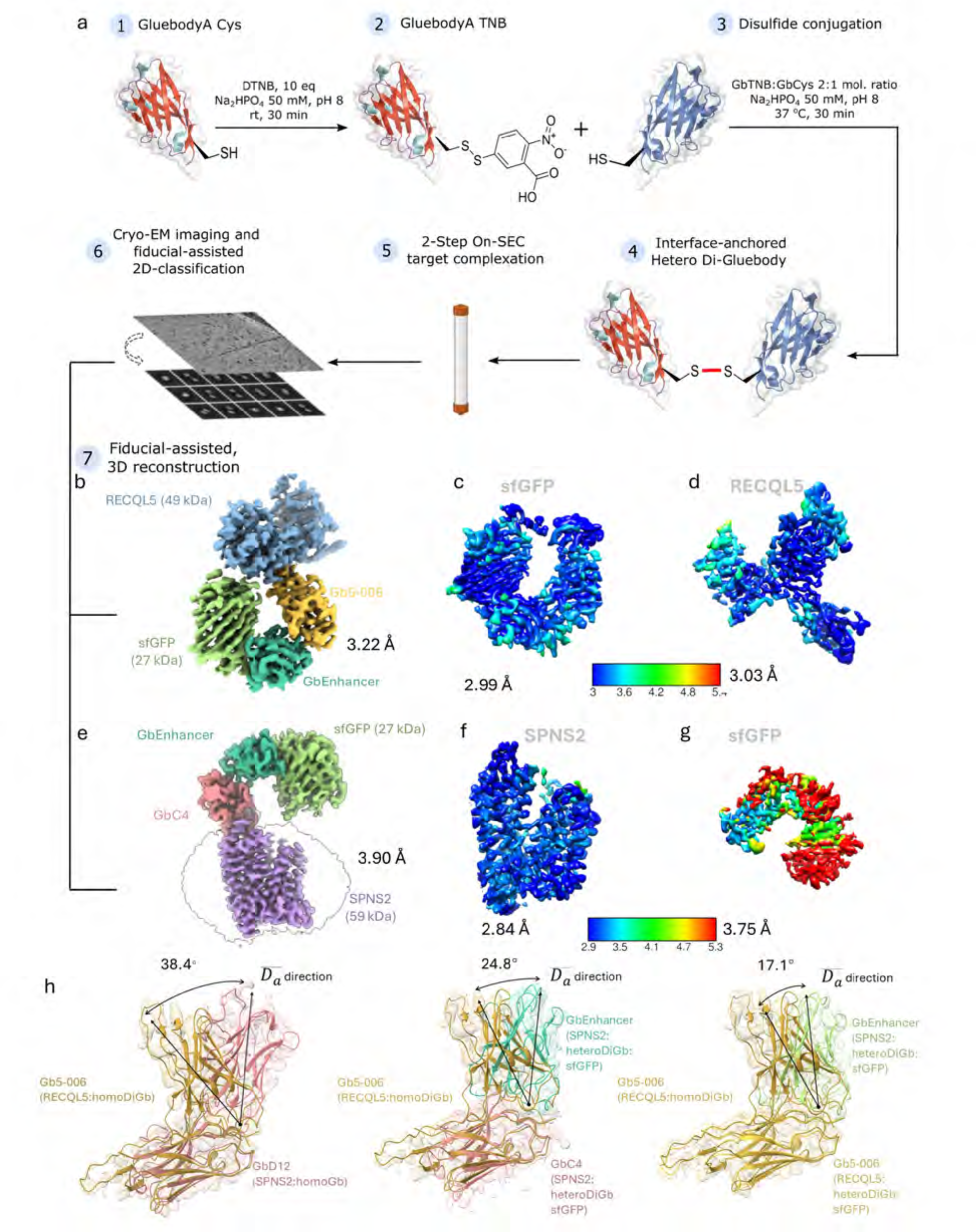
Synthesis of hetero Di-Gluebody (heteroDiGb) enables simultaneous high-resolution structure solution of two different small proteins. (a) Construction pipeline for heteroDiGb, via a trapped intermediate. (b) High-resolution cryo-EM map of RECQL5 and sfGFP in complex with heteroDiGb Gb5-006:GbEnhancer. Individual local refinement resulted in higher resolutions of (c) sfGFP. and (d) RECQL5. (e) High-resolution cryo-EM map of SPNS2 and sfGFP in complex with heteroDiGb GbC4:GbEnhancer. The intra-DiGb angle is indicated. Individual local refinement resulted in higher resolutions of (f) SPNS2 and (g) sfGFP. (h) Structural comparisons of homoDiGbD12, heteroDiGbs GbC4:GbEnhancer and Gb5-006:GbEnhancer with homoDiGb5-006. Intra-DiGb angles are indicated and 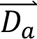 direction is defined in the methods section and Extended Data Figure 14.

Strikingly, using sequential purification, we generated heteroDiGbs bound to two distinct targets. Starting with a heteroDiGb of Gb5-006 and GbEnhancer, the RECQL5:heteroDiGb:sfGFP complex’s structure was determined to an overall resolution of 3.22 Å (**Figure 2b** and **Extended Data Figure 11**). Moreover, when we locally refined each target separately, we found much-improved individual maps of 3.03 Å for RECQL5 and 2.99 Å for sfGFP (**Figures 2c and 2d**, **Extended Data Figure 11**). In an essentially identical workflow using heteroDiGb GbC4:GbEnhancer, the SPNS2:heteroDiGb:sfGFP complex’s structure was determined to 3.90 Å overall (**Figure 2e** and **Extended Data Figure 12**), with 2.84 Å for SPNS2 and 3.75 Å for sfGFP, individually (**Figures 2f and 2g**, **Extended Data Figure 12**). These results clearly demonstrated the efficiency offered by heteromeric Di-Gluebodies in duplexing structure determination. Further validating these methods, all of the structures obtained were largely consistent with deposited structures, with the only changes being small inter-domain movements observed in RECQL5 and its slightly different Gluebody conformations (**Extended Data Figure 13**).

Notably the heteroDiGb strategy can also further overcome limitations of the homodimeric Gluebodies. While homoDiGbs improved resolvability and resolution for the RECQL5 and SPNS2 test cases, this strategy failed with the super-folder GFP (sfGFP)^19^. Alignment of a previously determined NbEnhancer:sfGFP complex to our determined homo Di-Gluebody structures, suggested that the two target molecules would likely clash in the homoDiGb complex despite flexibility in the intra-DiGluebody angle (**Figure 2h** and **Extended Data Figure 15**). By contrast, heteroDiGbs constructed from NbEnhancer and the RECQL5 nanobody yielded high-resolution results.

### Micromolar-affinity nanobodies are applicable for the Di-Gluebody approach

The ready generation of protein complexes under kinetic control that underpins our use of homoDiGbs and heteroDiGbs raises the intriguing question of needed affinities (and associated limits) of homo/heteroDiGbs for their targets. Interestingly, in each case the dissociation constants measured for each were weaker than for corresponding nanobodies (Nbs) or indeed monovalent Gbs (**Figure 3**, *K*_D_ 1.2 μM RECQL5 Nb → 9.0 μM for Gb equivalent → 23.4 μM for the homoDiGb; and from *K*_D_ 0.5 nM^19^ for GFP enhancer Nb → 5 nM for its Gb). Therefore, whilst we did not notice any practical effects on workflows for generating protein complexes (e.g. during size exclusion chromatography), these data importantly challenge the dogma of needed ‘tight binders’ and further highlight the utility of our kinetically-driven method in its accommodation, via such trapping, of complexes with even modest (micromolar) affinities.

**Figure 3.**
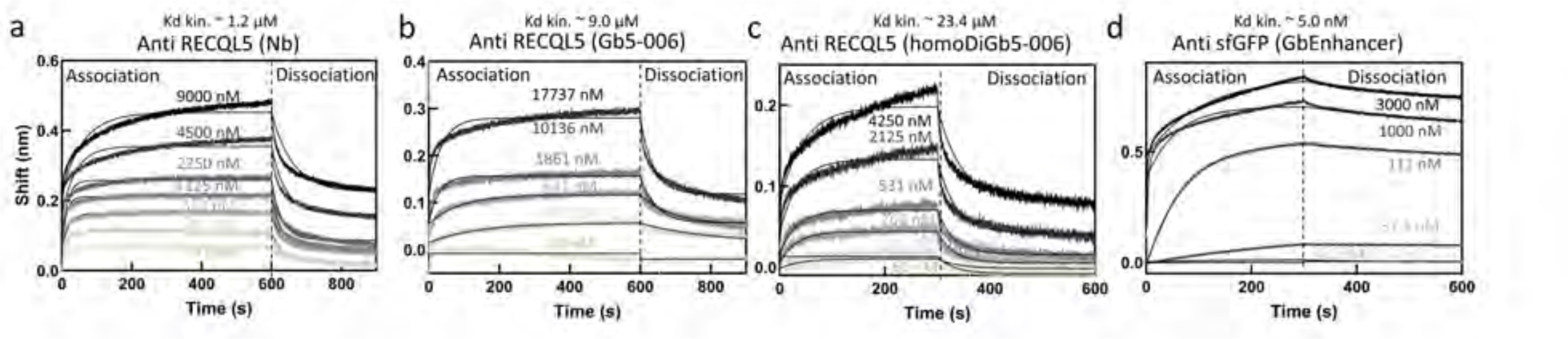
Kinetic trapping allows use of even modest underpinning affinities for protein targets. Bio-layer interferometric (BLI) measurements for RECQL5 Nb/Gb/homoDiGb and sfGFP Gb show affinity reductions after Gluebody mutagenesis and generation of homo Di-Gluebodies. BLI measurement plots with fitted lines are shown in the sequence of (a) anti-RECQL5 wild type nanobody, (b) anti-RECQL5 monomeric Gluebody Gb5-006, (c) anti-RECQL5 homoDiGb5-006 and (d) anti-sfGFP monomeric Gluebody GbEnhancer. The kinetic dissociation constants are indicated above the title of the plots individually. One replicate is shown in each plot.

### Interface-assisted, chemically-driven, and kinetic-controlled disulfide formation enables balanced, rigid protein assemblies

The striking ease of application of the homo/heteroDiGb method to cryo-EM-enabled structure solution led us to probe also its structural as well as biophysical origins, particularly in the non-canonical, sidechain-to-sidechain covalency that we exploit here to link protein modules. These analyses revealed a remarkable balance. All six structures have DiGbs at distinct intra-Gb angles (**Figure 4a**, **Extended Data Table 2**), showing the remarkable compatibility of the sidechain-to-sidechain covalent linkage that is trapped at site 12. Yet, at the same time, within each structure, the DiGb interfaces exhibit <10° of wobble, suggesting this interface is quite rigid and hence enabling refinement with C2 symmetry (**Figure 4b, c, d and e**, **Extended Data Figure 14**, and **Extended Data Table 2**).

**Figure 4.**
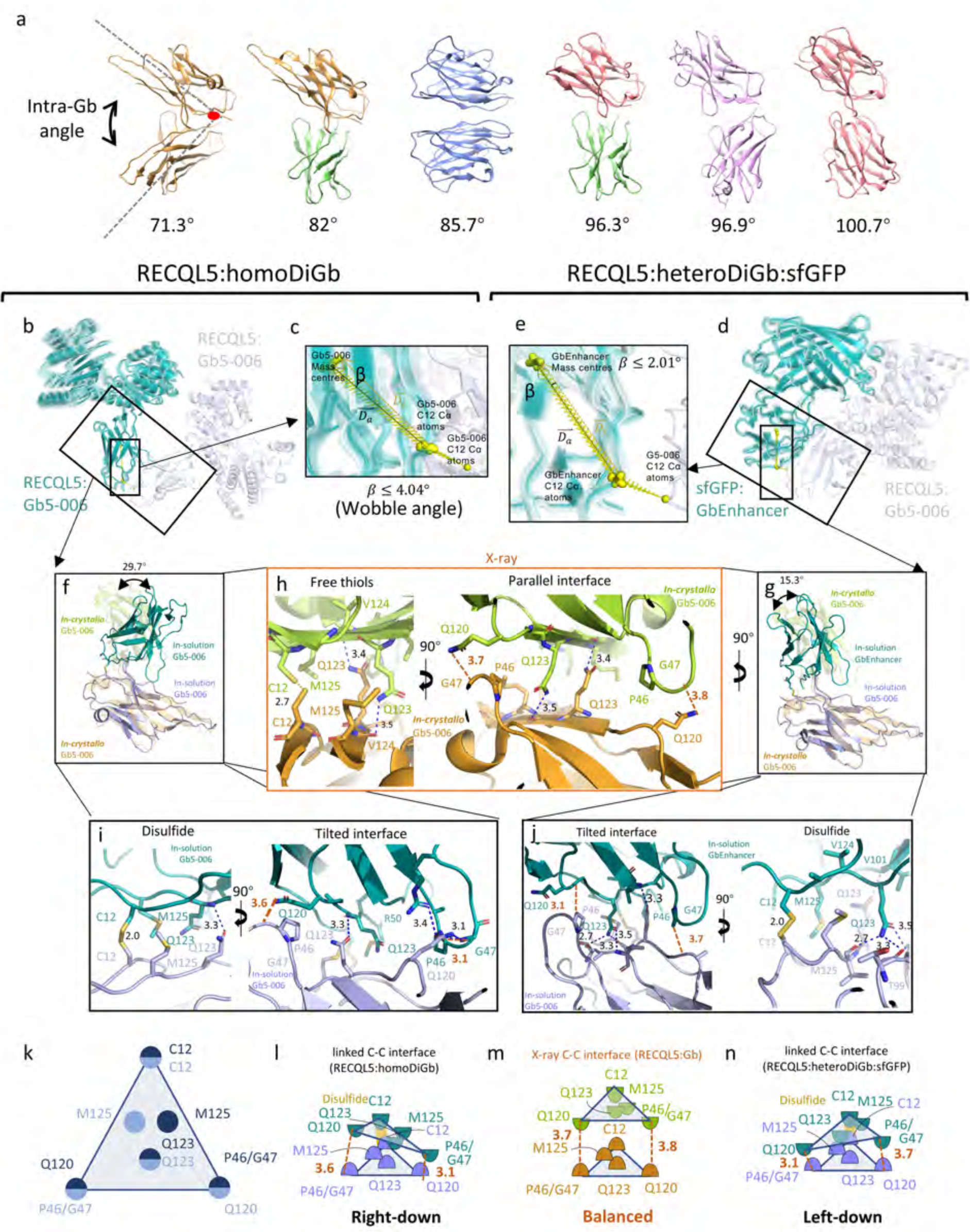
Side-chain-to-side-chain-linked DiGb interfaces are tilted and asymmetric. (a) Different DiGb pairs have unique intra-DiGb angles. The intra-DiGb angle is defined by the angle of the vectors from the Cα atom of Cys12 to the mass centre of respective Gluebodies. The represented structures from left to right are: RECQL5:homoDiGb, RECQL5:heteroDiGb:sfGFP, MBP:homoDiGb, SPNS2:heteroDiGb:sfGFP, Lysozyme:homoDiGb, SPNS2:homoDiGb. Within one structure, 3D variability analysis revealed the flexibility around the DiGb interfaces of (b) RECQL5:homoDiGb and (d) RECQL5:sfGFP:heteroDiGb with the respective insets (c) and (e) showing the degree of flexibility by wobbling angles. (f) Comparison of homoDiGb with the G5-006 crystallographic pattern. The pairs are aligned to one copy of Gb5-006. The angle deviation is indicated. (g) Comparison of heteroDiGb Gb5-006:GbEnhancer with the G5-006 crystallographic pattern. The pairs are aligned to one copy of Gb5-006. The angle deviation is indicated. The interacting residues of the C-C interfaces (h) in crystallo, (i) in homoDiGb5-006 and (j) heteroDiGb Gb5-006:GbEnhancer are demonstrated, with blue dashes showing hydrogen bonds, red dashes showing the minimum distances (not counting hydrogen atoms) between the P46/G47 and Q120 pairs. The (k) top view cartoon of the C-C interface for all three types and side view cartoons of (l) RECQL5:homoDiGb, (m) *in crystallo* RECQL5:Gb5-006 and (n) RECQL5:heteroDiGb:sfGFP are shown with key residues annotated and the minimum distances (not counting hydrogen atoms) between the P46/G47 and Q120 pairs in red dashes identical to (h-j)

By closely inspecting the interfaces of RECQL5:homoDiGb, RECQL5:sfGFP:heteroDiGb that are both trapped by covalency and comparing them with the *in crystallo* interface that is trapped by crystallization, we discovered subtle differences in the interfaces (**Figure 4f-4n**), causing effectively different intra-DiGb angles (**Figure 4f and 4g**). Both in solution and *in crystallo* interfaces are approximately arranged in equilateral triangular form with Cys12 at the top and Gln120 and Pro46/Gly47 pairs at the bottoms (**Figure 4h, 4i, 4j and 4k**). However, comparing the minimum distances between the interacting bottom residue pairs (Pro46/Gly47 and Gln120), the *in crystallo* generated interface is parallel whereas the in solution interfaces are tilted (**Figure 4h, 4i and 4j**). We noticed that in both cases, Gln123 always serves as a supporting residue, forming hydrogen bonds with the backbones on the opposing nanobody/Gluebody (Figure 4h, 4i and 4j), clearly highlighting the vital role of glue-ing in driving trapping. Due to tilt, hydrogen bonds only form in the closer half of the triangular interface (**Figure 4m and 4n**), together with the disulfide forming the interaction network.

The side chains of crucial residues also change the Di-Gluebody interface. Apart from the triangular Cys-Cys interface discussed above (**Figure 4k**, **Extended Data Figure 16**), we observed a much narrower interface in the SPNS2:homoDiGb structure. Both sides of residue 120 on the homoDiGb are lysines instead of the typical glutamine causing the interface to be shifted to a new minimum, with Pro46/Gly47 interacting instead with Gln44 (**Extended Data Figure 16**).

Finally, we also observed clear structural evidence for covalent bond formation as a driving trapping force for formation of the Di-Gluebody complex. In the MBP:homoDiGb structure, there is a unique interface different from those discussed above or any class of nanobody crystal contacts^16^. The sidechain-to-sidechain covalent disulfide bond gives the Gluebodies a ‘U-turn’ in this joint, enabling the first beta-sheet of both Gluebodies to come into contact, but not close enough to form direct hydrogen bonds (**Extended Data Figure 16**). Nonetheless, the resulting assembly proved rigid enough for cryo-EM structural determination, with or without local refinement, further highlighting the value of this kinetically-controlled approach.

## Discussion

Together, our results suggest that the success we see here in forming balanced, rigid assemblies is enabled by (i), homo- and hetero-DiGbs settling into robust conformational minima at non-canonical dimerization interfaces with sufficiently-balanced chemical flexibility and yet structural rigidity and (ii), the validity and robust nature of these interfaces being tested and reinforced by a chemically-driven, kinetically-controlled workflow that samples the equilibria that underpin the covalent interface-assisted events that are feasible in solution. Furthermore, the success of Di-Gluebodies to high-resolution structural solution is enabled by its relatively small mass contribution to the imaged complex while increasing the maximum particle diameter for better particle alignment (**Table 1**).

**Table 1.** DiGbs increase maximum particle diameters while maintaining the target volume proportions within the complex.

| Model | Targets | Target Volume Proportion (%) | Complex Max Diameter (Å) | Target Max Diameter (Å) |
| --- | --- | --- | --- | --- |
| RECQL5:homoDiGb | RECQL5 | 77.8 | 104.7 | 72.3 |
| MBP:homoDiGb | MBP | 75.4 | 108.3 | 64.5 |
| Lysozyme:homoDiGb | Lysozyme | 51.9 | 125.6 | 44.1 |
| SPNS2:homoDiGb | SPNS2 | 80.8 | 178.4 | 65.8 |
| SPNS2:heteroDiGb:sfGFP | SPNS2 | 52.6 | 116.5 | 65.8 |
|  | sfGFP | 24 |  | 49.9 |
| RECQL5:heteroDiGb:sfGFP | RECQL5 | 47.6 | 110.5 | 72.3 |
|  | sfGFP | 26.2 |  | 49.9 |

In summary, our ‘plug-and-play’ Di-Gluebody strategy improves cryo-EM resolvability and resolution of small proteins through rigidity and increased particle size with a simple and robust workflow, and can even enable simultaneous structure determination of two challenging targets in one dataset. Future exploitation of this modular ‘plug-and-play’ tool may allow even higher-order multiGb:target complexes or multi-specific assemblies when combined with other scaffold proteins.

The kinetic interface trapping that underpins this method was discovered *in crystallo* and taken to its current modular form through rare, kinetically-controlled non-canonical sidechain-to-sidechain covalent bond formation in solution. It should be noted therefore that the generic, rapid and modular nature of this method cannot and will not, essentially by definition, be achieved through traditional fusion protein methods. Moreover, our results also highlight that current dogmas of ’tight’ K_D_s, which indeed drive the assessment of most binding proteins, are less relevant in a kinetic scenario and may indeed prove inappropriate / inhibitory.

Finally, our rapid implementation of the homo/heteroDiGb method suggests that this will become a powerful, prospective tool for cryo-EM SPA structure determination.

## Methods

### Conversion of the nanobody numbering to the IMGT scheme

Due to variable lengths of nanobody CDRs, the same positions on the nanobody scaffold have different residue numbers in different nanobodies. Therefore, we used the IMGT scheme^18^ via the server ANARCI^22^ for consistent numbering and clarity in the Gluebody mutations (Figure 1a and Extended Data Figures 2a and 2b).

### Expression and purification of RECQL5, sfGFP, SPNS2, MBP and Lysozyme

RECQL5 was expressed and purified as previously described^16^, and then exchanged into a non-reducing buffer containing 5% Glycerol, 10 mM HEPES (pH7.5) and 500 mM NaCl using a PD10 columns (Cytiva #17-0851-01). SPNS2 was expressed and purified in DDM as described previously^10^.

The sfGFP and MBP protein was expressed using BL21-DE3-pRARE strain in auto-induction TB media (Formedia #AIMTB0260) containing 50 μg/mL Kanamycin and 0.01% antifoam 204 at 37°C for 5.5 hours followed by 40-44 hours at 18°C. Bacteria were harvested by centrifugation at 4000 g and re-suspended in a 3x cell pellet volume of base buffer (5% glycerol, 10 mM HEPES, pH 7.5 and 500 mM NaCl) supplemented with 30 mM imidazole, 1% triton, 0.5 mg/ml lysozyme, 10 μg/ml benzonase, and then stored in a -80°C freezer overnight for freeze-thaw cell lysis. Cell pellets were thawed in a room temperature water bath and then clarified by centrifugation at 5000 g for 1h. The supernatant was then applied to a Ni-NTA pre-packed column (Cytiva #11003399) pre-equilibrated with base buffer supplemented with 30 mM imidazole. After washing the Ni-NTA column with 10x column volume of base buffer, protein was eluted using 2.5 ml of base buffer supplemented with 500 mM imidazole and then immediately loaded onto a PD-10 column (Cytiva #17-0851-01) that had been base buffer-equilibrated. The sfGFP or MBP protein was then eluted from PD-10 columns. TEV protease at a 1:10 mass ratio and 20 mM imidazole were added to protein solution and incubated overnight. The next day, two Ni-NTA columns were pre-equilibrated with base buffer + 20 mM imidazole and the sfGFP or MBP solution with TEV was applied to the columns to remove sfGFP or MBP with uncleaved HIS_6_ tag, TEV protease and contaminants. Eluted fractions were collected, and flash frozen until use.

Lysozyme was purchased from Sigma (#L6876-1G) and dissolved in base buffer (5% glycerol, 10 mM HEPES, pH 7.5 and 500 mM NaCl) before complexation with the binding homo Di-Gluebody.

### Generation of homo Di-Gluebodies

The genes of Gluebodies for homo Di-Gluebody generation were amplified either from synthetic genes from TWIST Biosciences (Gb5-006, GbD12, GbS2A4 ^23^, GbH12 ^10^, GbLysozyme, GbHIV, GbRBD3) or existing clones with Gluebody mutations implemented in the primers (GbEnhancer and GbMBP^24^). GbEnhancer, GbLysozyme, GbHIV and GbMBP had the minimal Gluebody mutations S7N/L12C/Q14K/T125M implemented. Gb5-006 had solubility mutations G40T/Q49E/L52W/I101V plus K84E in addition to S7N/L12C/Q14K/T125M. GbD12 had S7N/L12C/Q14K/K84E/P123Q/T125M implemented. GbS2A4 had Q5V/S7N/L12C/Q14K/Q44R/T125M implemented. GbH12 had L2V/S7N/L12C/Q14K/K84E/P123Q/T125M implemented. GbRBD3 had S7N/L12C/Q14K/T125M/K84E implemented. They all share the core mutations of S7N/L12C/Q14K/T125M. NbH12 (wildtype of GbH12) was generated in the same screening batch as NbD12 (wildtype of GbD12) ^10^. The amplified GbMBP gene fragment was cloned onto the pNIC-NHStIIT vector, GbGFP gene fragment was cloned onto the pNIC-GST-bio vector and all other amplified genes were subsequently cloned onto the pNIC-MBP2-LIC vector, by ligation-^25^1. The Gluebodies were produced as previously described except that no reducing agent was used throughout^16^1. The purified monomeric Gluebodies were exchanged into 50 mM sodium phosphate (Na_2_HPO_4_) buffer (pH 8.0) using a PD M-iniTrap G25 columns (Cytiva #28918007), snap-frozen and stored at -80 °C until further use.

To generate homo Di-Gluebodies, purified Gluebody at 2 mg/ml in 50 mM Na_2_HPO_4_, pH 8, was treated with 50 uM of Cu(II) acetate, and the resulting solution was incubated at 37 °C temperature for 30 minutes. Conversion was monitored by intact mass spectrometry on Waters G2-XS QToF mass spectrometers equipped with a Waters Acquity UPLC. Separation was achieved using a Thermo Scientific ProSwift RP-2H monolithic column (4.6 mm × 50 mm) using water + 0.1% formic acid (solvent A) and acetonitrile + 0.1% formic acid (solvent B) as mobile a 5-min linear gradient. Spectra were deconvoluted using MassLynx 4.1(Waters). Upon completion, the Di-Gluebody was purified by SEC chromatography using a Superdex Increase 75 pg 16/600 column (Cytiva #28-9893-33). Detailed data for Gb5-006, GbD12, GbLysozyme and GbMBP are shown in Extended Data Figures 3a-d. Additional pairs of homoDiGbs are shown in Figures 3e-3l.

### Generation of hetero Di-Gluebodies

The genes of Gluebodies for hetero Di-Gluebody generation were amplified either from synthetic genes from TWIST Biosciences (Gb5-006, GbH12 ^10^, GbRBD1 ^26^ and GbRBD6 ^26^) or existing clones with Gluebody mutations implemented in the primers (GbEnhancer and GbC4 ^10^). GbC4 had L2G/S7N/L12C/Q14K/T125M implemented. GbRBD1 and GbRBD6 had S7N/L12C/Q14K/T125M/K84E implemented. They all share the core mutations of S7N/L12C/Q14K/T125M. NbC4 (wildtype of GbC4) was generated in the same screening batch as NbD12 (wildtype of GbD12) ^10^. All amplified genes were subsequently cloned onto the pNIC-MBP2-LIC vector, by ligation-independent cloning^25^. The Gluebodies were produced as previously described except that no reducing agent was used throughout purification^16^. The purified monomeric Gluebodies were exchanged into 50 mM sodium phosphate (Na_2_HPO_4_) buffer (pH 8.0) using a PD MiniTrap G-25 columns (Cytiva #28918007), snap-frozen and stored at -80 °C until further use.

For generation of hetero Di-Gluebodies, the first Gluebody (GluebodyA) at 1.0 mg/mL in 50 mM Na_2_HPO_4_ (pH 8.0), was treated with DTT at 10x molar concentration of GluebodyA at room temperature. After 15 min, DTT was removed from GluebodyA using a PD MiniTrap G-25 columns (Cytiva #28918007) equilibrated in Na_2_HPO_4_ 50 mM, pH 8.0. GluebodyA was then treated with 10x the molar concentration of 5,5ʹ-dithiobis-2-nitrobenzoic acid (DTNB) at room temperature. After 30 minutes, DTNB was removed by buffer exchange using G-25 columns in 50 mM Na_2_HPO_4_, pH 8 and concentrated with Ultra-0.5 3 kDa centrifugal filters (Amicon #UFC500308). Conversion of GlueBodyA’s Cys12 to Cys12TNB was confirmed on Waters G2-XS QToF mass spectrometers equipped with a Waters Acquity UPLC. Separation was achieved using a Thermo Scientific ProSwift RP-2H monolithic column (4.6 mm × 50 mm) using water + 0.1% formic acid (solvent A) and acetonitrile + 0.1% formic acid (solvent B) as mobile a 5-min linear gradient. Spectra were deconvoluted using MassLynx 4.1 (Waters). Samples of GluebodyB at 1.0 mg/ml were reduced with DTT using the same protocol as for GluebodyA, desalted using minitrap G-25 columns, and then mixed with GluebodyA-Cys12TNB at 2x the molar concentration and incubated for 30 minutes at 37 °C. Small molecule byproducts from the resulting mixture were removed using a G-25 column in a buffer containing 50 mM Na_2_HPO_4_ (pH 8.0), flash frozen in liquid nitrogen and kept at –80 °C until further use. Detailed data for Gb5-006:GbEnhancer, GbC4:GbEnhancer and additional pairs of hetero Di-Gluebodies are shown in Extended Data Figures 4a and 4b.

### In vitro reconstitution of Di-Gluebody:target complexes

All proteins prior to this stage are in a buffer without reducing agents as described in the previous sections, and thus all proteins are under non-reducing conditions thereafter.

To prepare complexes of RECQL5 with the homo Di-Gluebody Gb5-006, the Di-Gluebody was mixed with RECQL5 at a molar ratio of 1:2 and subsequently purified by size exclusion chromatography on a Superdex 200 10/300 GL Increase column (Cytiva #28-9909-44), pre-equilibrated in 20 mM HEPES (pH 7.5), 150 mM NaCl. The dimer peak was pooled, concentrated by 10 kDa Vivaspin 20 centrifugal concentrators (Vivaproducts #VS2001) and used for cryo-EM specimen preparation (Extended Data Figure 5a). The same procedure applies to Lysozyme (Extended Data Figure 5c) and MBP (Extended Data Figure 5d).

To prepare complexes of sfGFP with the homo Di-Gluebody GbEnhancer, the Di-Gluebody was mixed with sfGFP at a molar ratio of 1:2 and subsequently purified by size exclusion chromatography on a Superdex 200 10/300 GL Increase column (Cytiva #28-9909-44), pre-equilibrated in 20 mM HEPES (pH 7.5), 150 mM NaCl. The dimer peak was pooled, concentrated by 10 kDa Vivaspin 20 centrifugal concentrators (Vivaproducts #VS2001) and used for cryo-EM specimen preparation.

To prepare complexes of SPNS2 with the homo Di-Gluebody GbD12, the purified Di-Gluebody was first supplemented with 0.025% DDM (Anatrace #D310LA) and then mixed with SPNS2 at a molar ratio of 1:2. The complex was then purified by size exclusion chromatography on a Superdex Increase 200 10/300 GL column (Cytiva #28-9909-44) pre-equilibrated in 20 mM HEPES (pH 7.5), 150 mM NaCl, 0.025% DDM. The dimer peak was pooled, concentrated by 100 kDa Vivaspin 20 centrifugal concentrators (Vivaproducts #VS2041), and used for cryo-EM specimen preparation (Extended Data Figure 5b).

The RECQL5:heteroDiGb:sfGFP complex was prepared by first mixing Gb5-006:GbEnhancer hetero Di-Gluebody with RECQL5 at a molar ratio of 1:1.5. The RECQL5:heteroDiGb complex was then purified by size exclusion chromatography on the Superdex Increase 75 10/300 GL column (Cytiva #29-1487-21) pre-equilibrated in a buffer condition of 20 mM HEPES (pH 7.5), 150 mM NaCl. The peak corresponding to the hetero Di-Gluebody and RECQL5 complex was pooled (Extended Data Figure 5f left panel), mixed with sfGFP at a molar ratio of 1:1.5, and further purified by size exclusion chromatography on the Superdex Increase 75 10/300 GL column. The peak containing both targets was then pooled, concentrated by 10 kDa Vivaspin 20 centrifugal concentrators (Vivaproducts #VS2001), and used for cryo-EM specimen preparation (Extended Data Figure 5f right panel).

The SPNS2:heteroDiGb:sfGFP complex was prepared by first mixing GbC4:GbEnhancer hetero Di-Gluebody with sfGFP at a molar ratio of 1:1.5. The sfGFP:heteroDiGb complex was then purified by size exclusion chromatography on the Superdex Increase 200 10/300 GL column (Cytiva #28-9909-44) pre-equilibrated in a buffer condition of 20 mM HEPES (pH 7.5), 150 mM NaCl, 0.025% DDM. The peak corresponding to the hetero Di-Gluebody and sfGFP complex was pooled (Extended Data Figure 5e left panel), mixed with SPNS2 at a molar ratio of 1:1, and further purified by size exclusion chromatography on the Superdex Increase 200 10/300 GL column. The peak containing both targets was then pooled, concentrated by 100 kDa Vivaspin 20 centrifugal concentrators (Vivaproducts #VS2041), and used for cryo-EM specimen preparation (Extended Data Figure 5e right panel).

### Cryo-EM specimen preparation and data acquisition

Cryo-EM grids of the RECQL5:homoDiGb, sfGFP:homoDiGb, MBP:homoDiGb, Lysozyme:DiGb and RECQL5:heteroDiGb:sfGFP complexes were prepared by applying freshly purified complexes to Au C-flat, 2/1, 200 mesh (Jena Bioscience #X-302-AU200) at 0.8 mg/ml, 0.75 mg/ml, 2.5 mg/ml, 0.85 mg/ml and 1.1 mg/ml, respectively. Grids were blotted using a Vitrobot (FEI) at 4 °C and 100 % humidity for 4 seconds with the force of -6 and 5-second waiting time, followed by plunging into liquid ethane.

Cryo-EM grids of the SPNS2:homoDiGb and SPNS2:heteroDiGb:sfGFP complexes were prepared by applying freshly purified complex to Quantifoil Copper, 1.2/1.3, 300 mesh (Quantifoil) at 9.0 mg/ml and 4.2 mg/ml, respectively. Grids were blotted using a Vitrobot (FEI) at 4 °C and 100 % humidity for 8 seconds with force of -10 and 5-second waiting time, followed by plunging into liquid ethane.

Cryo-EM data were collected using a FEI Titan Krios operating at 300 kV with a Gatan K3 with GIF Quantum camera or Falcon 4 with GIF Quantum camera. All data were automatically collected using EPU software (Thermo Fisher) with defocus ranging targeting -1.5 µM to -2.5 µM for SPNS2:homoDiGb and SPNS2:heteroDiGb:sfGFP complexes, -1.2 µM to -2.6 µM MBP:homoDiGb, Lysozyme:homoDiGb or -1.5 µM to -3 µM for RECQL5:homoDiGb, and RECQL5:heteroDiGb:sfGFP complexes. Other parameters such as magnification, total dose and frames used varied between different sample collections and are provided in Extended Data Table 1.

### Image processing of electron micrographs

All datasets were subject to a similar protocol for image processing and reconstruction via CryoSPARC^27^. Raw micrographs were motion corrected, contrast transfer function (CTF) estimated, and curated manually to remove those images with poor image quality (i.e. CTF fit > 4 Å, ice thickness > 1.1 and astigmatism > 1000). For each dataset, 500 micrographs were then used for blob picking and several rounds of 2D classifications. The particles with the best 2D class averages were selected for Topaz training and particle picking using all micrographs. After 2D classifications of Topaz picked particles, all particles from well-resolved 2D classes were merged after removing duplicated particles. The output particles were used for *ab-initio* reconstruction, followed by heterogeneous refinement. Well-resolved classes were selected for non-uniform refinement (with C2 symmetry applied to RECQL5:homoDiGb, SPNS2:homoDiGb and MBP:homoDiGb. C1 symmetry was applied for Lysozyme:DiGb, SPNS2:heteroDiGb:sfGFP complexes and RECQL5:heteroDiGb:sfGFP complexes), followed by local refinement with target-specific masks. The masks were generated using the ‘molmap’ command in UCSF Chimera^28^, based on roughly docked atomic models for the target proteins. For the homo Di-Gluebody complexes (except Lysozyme:homoDiGb), the particles were symmetry expanded, followed by further rounds of 3D classifications. The particles from best-resolved classes were then merged for a final round of local refinement. Maps were further sharpened by DeepEMhancer^29^.

In all cases, the resolution was determined by GS-FSC. The local resolution estimation was calculated via cryoSPARC and presented using UCSF Chimera^28^ based on the output maps from CryoSPARC. 3D conformational variability analysis was carried out using 3DVA^30^. The processing details for each dataset is further shown systematically in Extended Data Figures 7c, 8c, 9c, 10c, 11c and 12c.

### Model building and refinement

Initial models of RECQL5 and Gb5-006 were obtained from deposited crystal structures (PDB: 7ZMV)^16^. Those of sfGFP and GbEnhancer were obtained by modifying the existing models (PDB: 3K1K). For Lysozyme and GbLysozyme, the initial model was obtained from a deposited structure (PDB:6JB2). For MBP and GbMBP, the initial model was obtained from a deposited structure (PDB:5M13). Models of SPNS2, GbD12 and GbC4 used the SPNS2:NbD12 structure published previously (PDB: 8QV6)^10^ as starting models. Model were rebuilt as necessary in COOT^31^, and then refined with PHENIX real-space refinement^32^.

### Affinity measurement of Nanobodies/Gluebodies

Purified RECQL5 and sfGFP proteins were biotinylated using EZ-Link Sulfo-NHS-LC-Biotinylation Kit (Thermo Scientific #21435) following the manufacturer’s instructions. Bio-layer Interometry was then performed Using Octet RED384. All experiments were performed using buffer containing 10 mM HEPES, 150 mM NaCl at pH 7.5 for dilution and incubation. All incubation was done while shaking at 1000 rpm at room temperature. All experiments were done in triplicates.

For Affinity measurements of anti-RECQL5 Nb, Gb5-006 (monomer), the biotinylated target protein solution RECQL5 was diluted to 3 μM. Streptavidin coated biosensors (ForteBio Lot#2104023111) were first incubated in buffer for 1 minute, and then dipped into protein solution for 2 minutes, and back to buffer again for 1 minute. The arrays of sensors were then dipped into 8 concentrations (including a 0 concentration for subtraction) of Nb, Gb5-006 and homoDiGb5-006, respectively for 10 minutes for the association step. Finally the sensors were transferred to buffer for the dissociation step for 5 minutes to complete the measurement. The concentration gradients for anti-RECQL5 Nb is 9 μM, 4.5 μM, 2.250 μM, 1.125 μM, 562.5 nM, 281.25nM, 140.625nM and 0. The concentration gradients for anti-RECQL5 Gb5-006 is 17.7 μM, 10.1 μM, 4.3 μM, 1.9 μM, 620.6 nM, 206.9 nM, 69.0 nM and 0.

For Affinity measurements of anti-RECQL5 homoDiGb, the biotinylated target protein solution RECQL5 was diluted to 1.62 μM. Streptavidin coated biosensors (ForteBio) were first incubated in buffer for 1 minute, and then dipped into protein solution for 2 minutes, and back to buffer again for 2 minute. The arrays of sensors were then dipped into 8 concentrations (including a 0 concentration for subtraction) of GbEnhancer for 5 minutes for the association step. Finally the sensors were transferred to buffer for the dissociation step for 5 minutes to complete the measurement. The concentration gradients for anti-RECQL5 DiGb5-006 is 4.3 μM, 2.1 μM, 1.1 μM, 531.25 nM, 265.6 nM, 132.8 nM, 66.4nM and 0.

For Affinity measurements of anti-sfGFP GbEnhancer, the biotinylated target protein solution sfGFP was diluted to 4.5 μM. Streptavidin coated biosensors (ForteBio #18-5019) were first incubated in buffer for 1 minute, and then dipped into protein solution for 2 minutes, and back to buffer again for 2 minutes. The arrays of sensors were then dipped into 8 concentrations (including a 0 concentration for subtraction) of GbEnhancer for 5 minutes for the association step. Finally the sensors were transferred to buffer for the dissociation step for 5 minutes to complete the measurement. The concentration gradients for anti-GFP GbEnhancer is 3μM, 1μM, 333nM, 111nM, 37.3nM, 12.4nM, 4.1nM and 0.

For data processing, all traces with Gluebody/Nanobody concentrations above 0 were subtracted by the values from the 0 concentration trace, and aligned to the beginning of the association step. Non-linear regression was performed using Graphpad Prism 10.2.3 to estimate the dissociation constants of the Gluebodies/Nanobodies.

### Dynamic analysis of the Di-Gluebody interface

Selected particles used for the final rounds of non-uniform refinement^33^ for the dimer structures of RECQL5:homoDiGb, SPNS2:homoDiGb, RECQL5:heteroDiGb:sfGFP, SPNS2:heteroDiGb:sfGFP structures were used as inputs of 3D variability analysis on CryoSparc^27^, yielding 60 density maps. The atomic models for RECQL5:homoDiGb, SPNS2:homoDiGb, RECQL5:heteroDiGb:sfGFP, SPNS2heteroDiGb:sfGFP were then split at the Di-Glubody interface, and each model half docked into the 3D variability density maps using ChimeraX^34^.

The movements of the Di-Gluebody interfaces were analyzed by first aligning a reference (static) Gluebody. Vectors 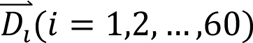 and its average 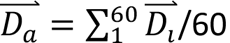 defined by each substituent Gluebody’s center of mass and the Cα atoms of Cys12, were then calculated for both static and the unaligned (mobile) Gluebody. Motion within a single Di-Gluebody complex was quantified as the wobble angle 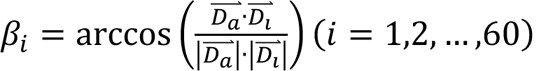 using the mobile Gluebody vectors, while the intra-DiGluebody angle was calculated between the 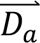 of the static and mobile Gluebodies.

## Supporting information

Supplementary Methods and Discussion

## Acknowledgments

We acknowledge Diamond Light Source for access and support of the cryo-EM facilities at the UK national Electron Bio-imaging Center (eBIC, proposal EM20223 and NT21004), funded by the Wellcome Trust, MRC, and BBSRC. Chemistry at The Rosalind Franklin Institute is supported by the EPSRC (EP/V011359/1). The Division of Structural Biology is a part of the Centre for Human Genetics at the University of Oxford, which was funded by Wellcome Trust Core Grant Number 090532/Z/09/Z. Electron microscopy provision was provided through the OPIC electron microscopy facility, a UK Instruct-ERIC Centre, which was founded by a Wellcome JIF award (060208/Z/00/Z) and was supported by a Wellcome equipment grant (093305/Z/10/Z). Computation was performed at the Oxford Biomedical Research Computing (BMRC) facility, a joint development between the Wellcome Centre for Human Genetics (Wellcome Trust Core Award Grant Number 203141/Z/16/Z) and the Big Data Institute (BDI) supported by Health Data Research UK and the NIHR Oxford Biomedical Research Centre. P.Z. was supported by ERC AdG (101021133) and Wellcome Investigator awards (206422/Z/17/Z). D.B.S. was supported by the Innovative Medicines Initiative 2 Joint Undertaking (JU) under grant agreement No 875510. The JU receives support from the European Union’s Horizon 2020 research and innovation program and EFPIA and Ontario Institute for Cancer Research, Royal Institution for the Advancement of Learning McGill University, Kungliga Tekniska Hoegskolan, Diamond Light Source Limited.

## Data availability

Cryo-EM density maps have been deposited in the EMDB under accession codes EMD-19331, EMD-19332, EMD-19333, EMD-19334, EMD-19335, EMD-19336, EMD-19337, EMD-19338, EMD-19339, EMD19340, EMD-50430, EMD-50432, EMD-50433, EMD-50525 and corresponding coordinate files have been deposited in PDB under accession codes 8RL5, 8RL6, 8RL7, 8RL8, 8RL9, 8RLA, 8RLB, 8RLC, 8RLD, 8RLE, 9FGV, 9FGX,9FGY, 9FQK. The map and model IDs are detailed in Extended Data Table 1. All other data are available upon request.

## Author contributions

M.Y., G.Y., D.M., F.V.D, B.G.D. and R.J.C.G. conceived the project. M.Y and G.Y. cloned, expressed and purified Gluebodies. D.M. generated and characterised Di-Gluebodies. H.L. expressed and purified SPNS2. M.F. cloned the sfGFP construct. M.Y. expressed and purified RECQL5 and sfGFP. M.Y., G.Y. and D.M. performed *in vitro* complex reconstitution. G.Y. and M.Y. performed cryo-EM grid freezing. G.Y. acquired and processed the cryo-EM data. M.Y. built and refined the atomic models. M.Y. performed 3D variability analyses. M.Y. performed the BLI affinity measurements. M.Y., P.Z., G.Y., D.M., D.B.S. and R.J.C.G. wrote the initial manuscript. All authors discussed and edited the manuscript. M.Y., D.B.S, F.V.D., B.G.D, and R.J.C.G. supervised the research.

## Competing interests

The authors declare no competing interests.

**Supplementary Materials**

---

**Extended Data Figure 1.**
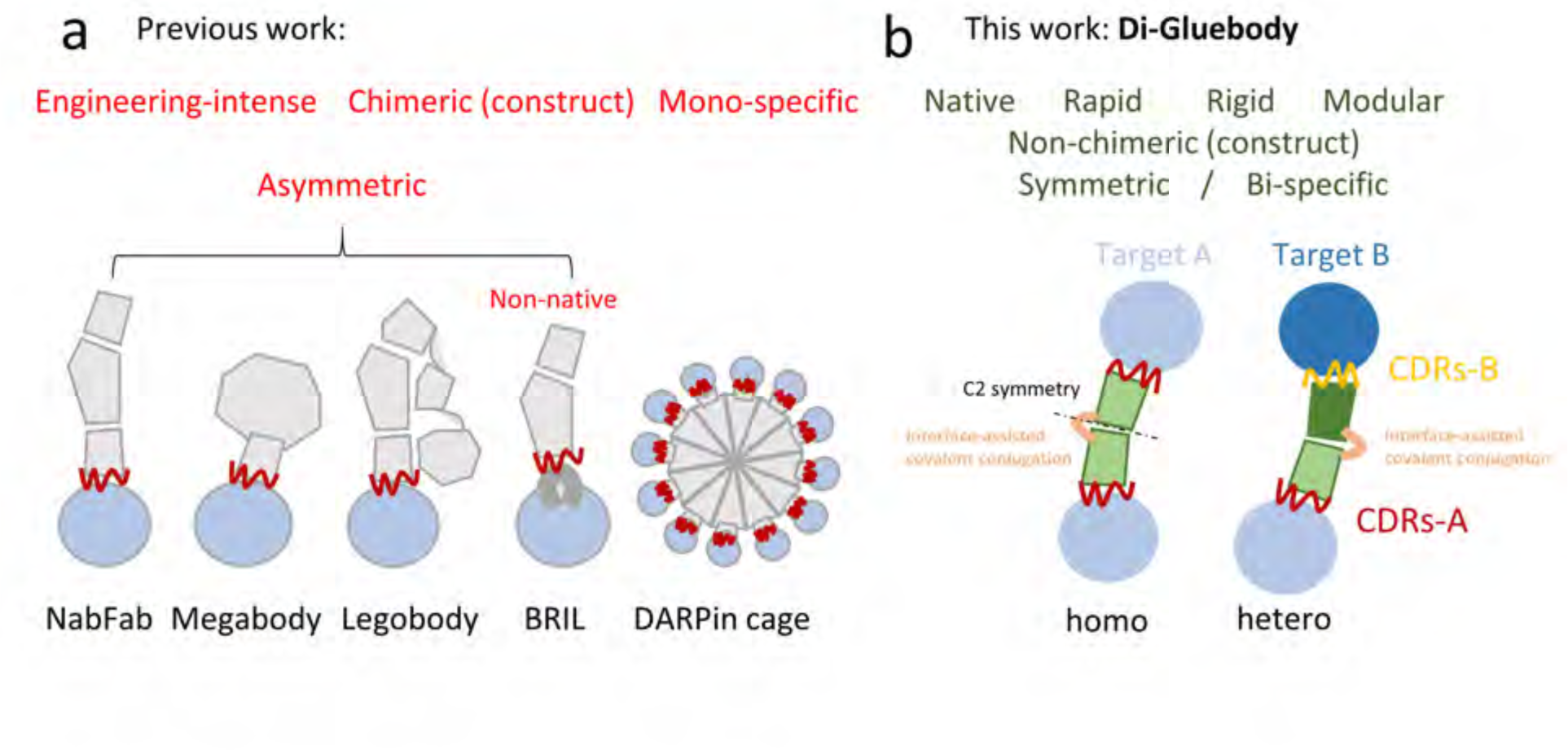
Comparison of Di-Gluebody and existing tools to make small proteins into larger complexes. (a) Previous work. Scaffold proteins are coloured grey, target proteins are in blue and CDRs are in red. (b) Di-Gluebody (this work). Monomeric Gluebodies are coloured deep and light green, target proteins are in deep and light blue, disulfides are in orange, and CDRs are in red and yellow.

**Extended Data Figure 2.**
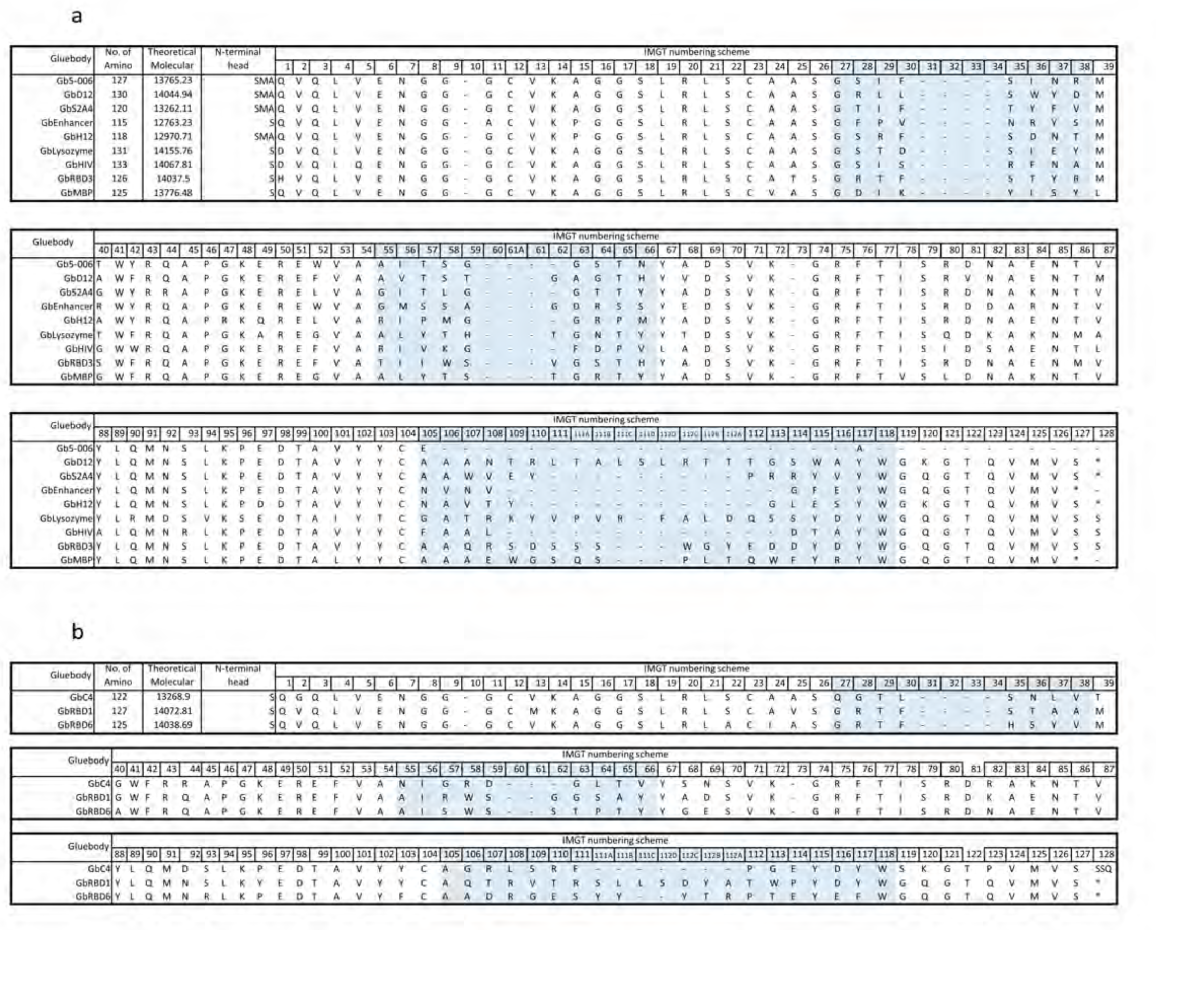
Gluebody sequences aligned in the IMGT scheme. (a) Sequence information of 10 Gluebodies used in homoDiGbs. (b) Sequence information of 3 additional Gluebodies used in heteroDiGbs. CDRs of the Gluebodies are highlighted in cyan.

**Extended Data Figure 3.**
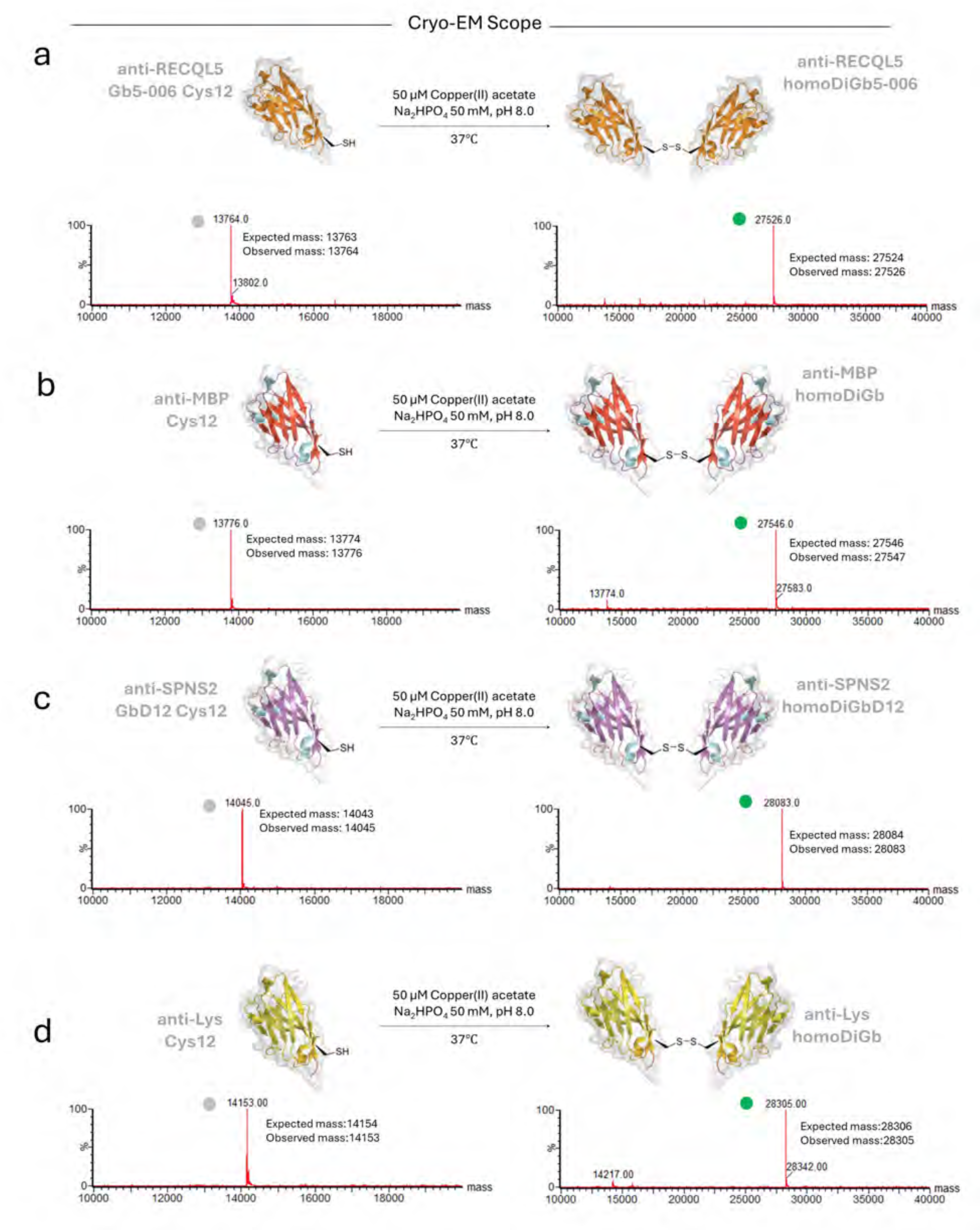
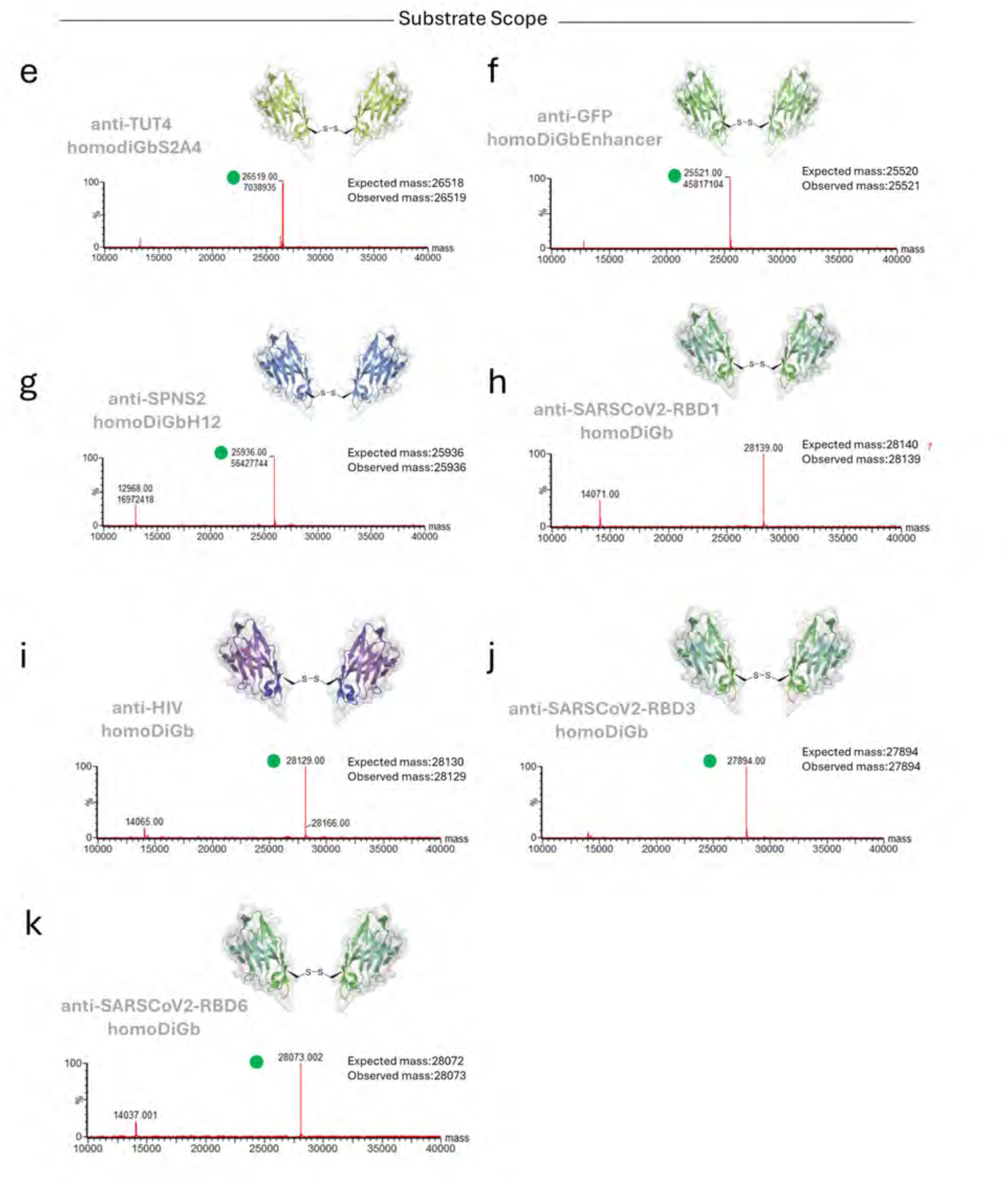
Modular generation of homo Di-Gluebodies (homoDiGbs) for cryo-EM. (a) Gb5-006 (b) GbMBP (c) GbD12 (d) GbLysozyme. Mass Spec peaks for Gluebody monomers and homoDiGb are indicated by blobs in grey and green, respectively.

**Extended Data Figure 4.**
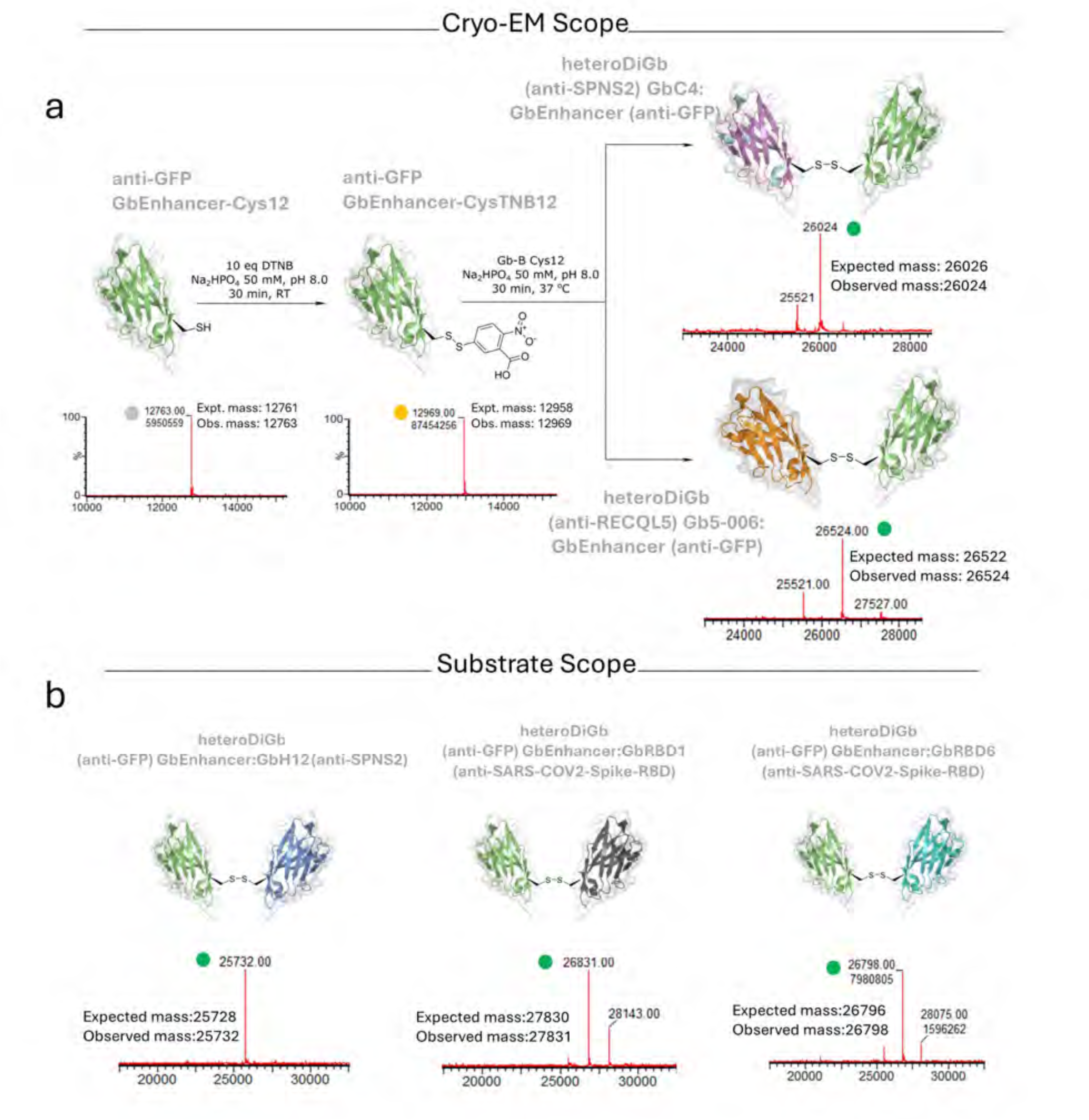
Modular generation of hetero Di-Gluebodies (heteroDiGbs). (a) Generation process of heteroDiGbs used in the cryo-EM studies (GbC4:GbEnhancer and Gb5-006:GbEnhancer) (b) More pairs of homoDiGbs generated demonstrating its modularity. Mass Spec peaks for Gluebody monomers, Gluebody-TNB conjugates and heteroDiGbs are indicated by blobs in grey, yellow and green, respectively.

**Extended Data Figure 5.**
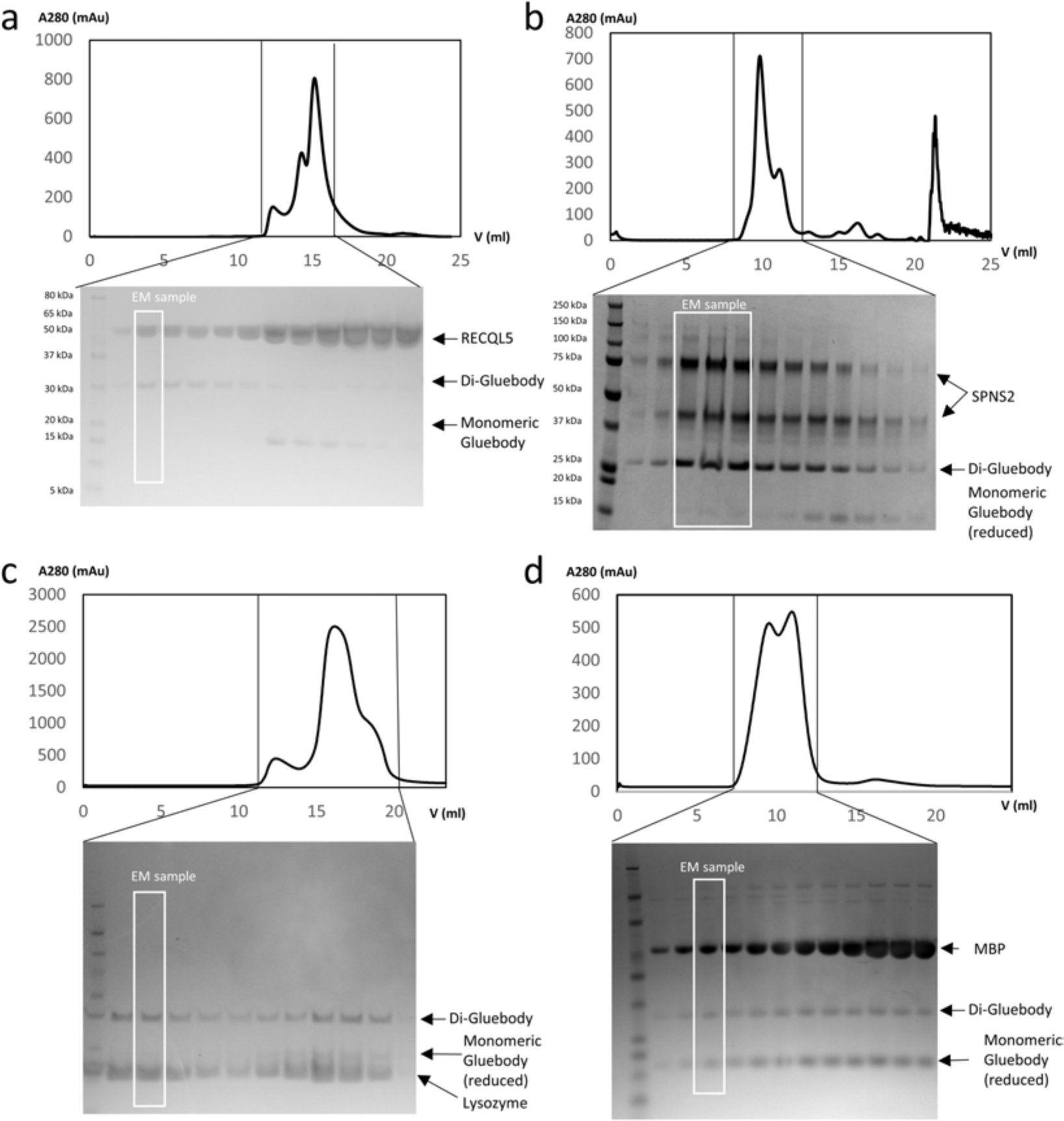
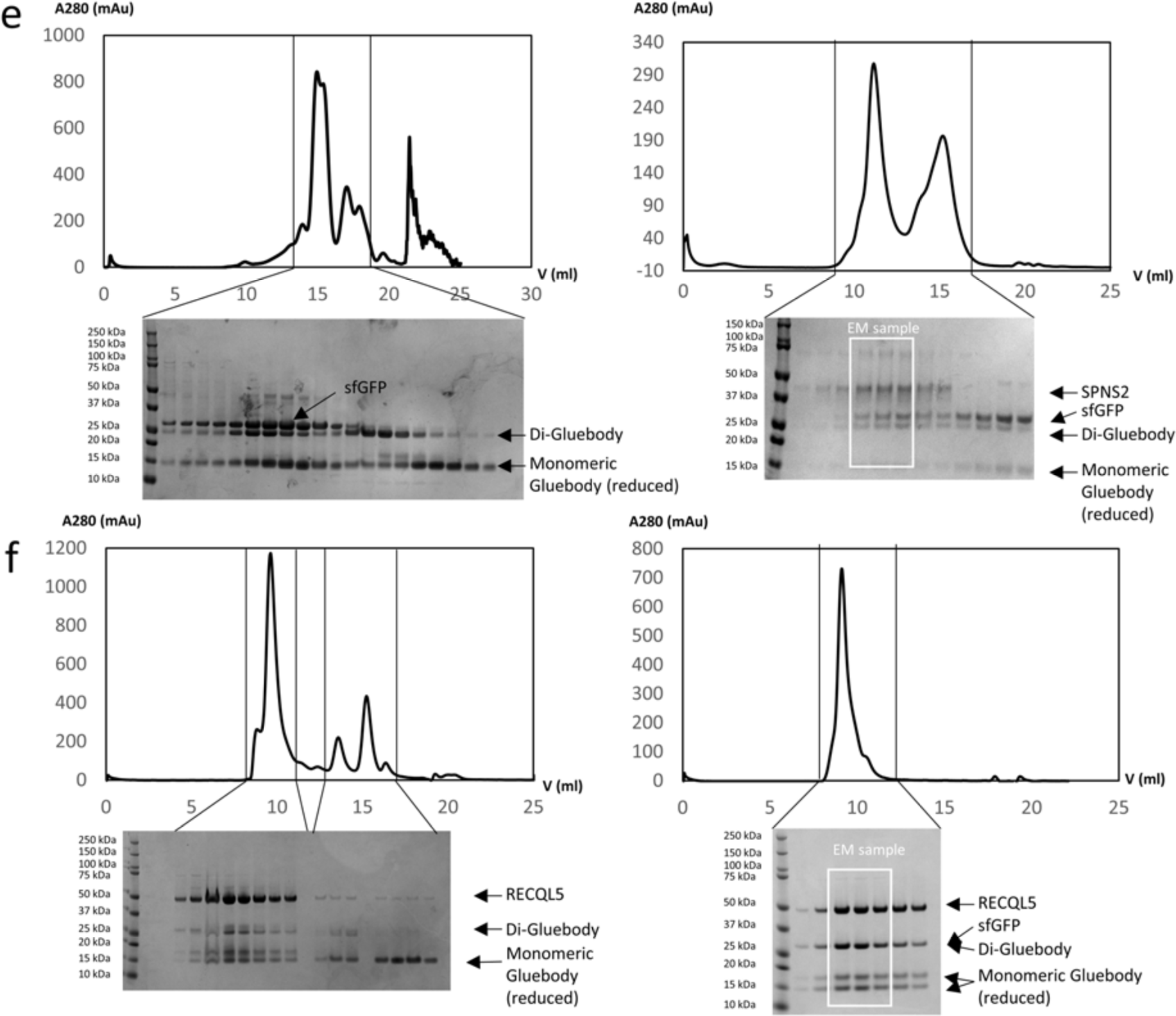
Homo and hetero dimer complex reconstitutions with size exclusion chromatography (SEC). (a) Purification of RECQL5 in complex with the homoDiGb5-006. (b) Purification of SPNS2 in complex with the homoDiGbD12. (c) Purification of lysozyme in complex with the homoDiGbLysozyme. (d) Purification of MBP in complex with the homoDiGb MBP.

**Extended Data Figure 6.**
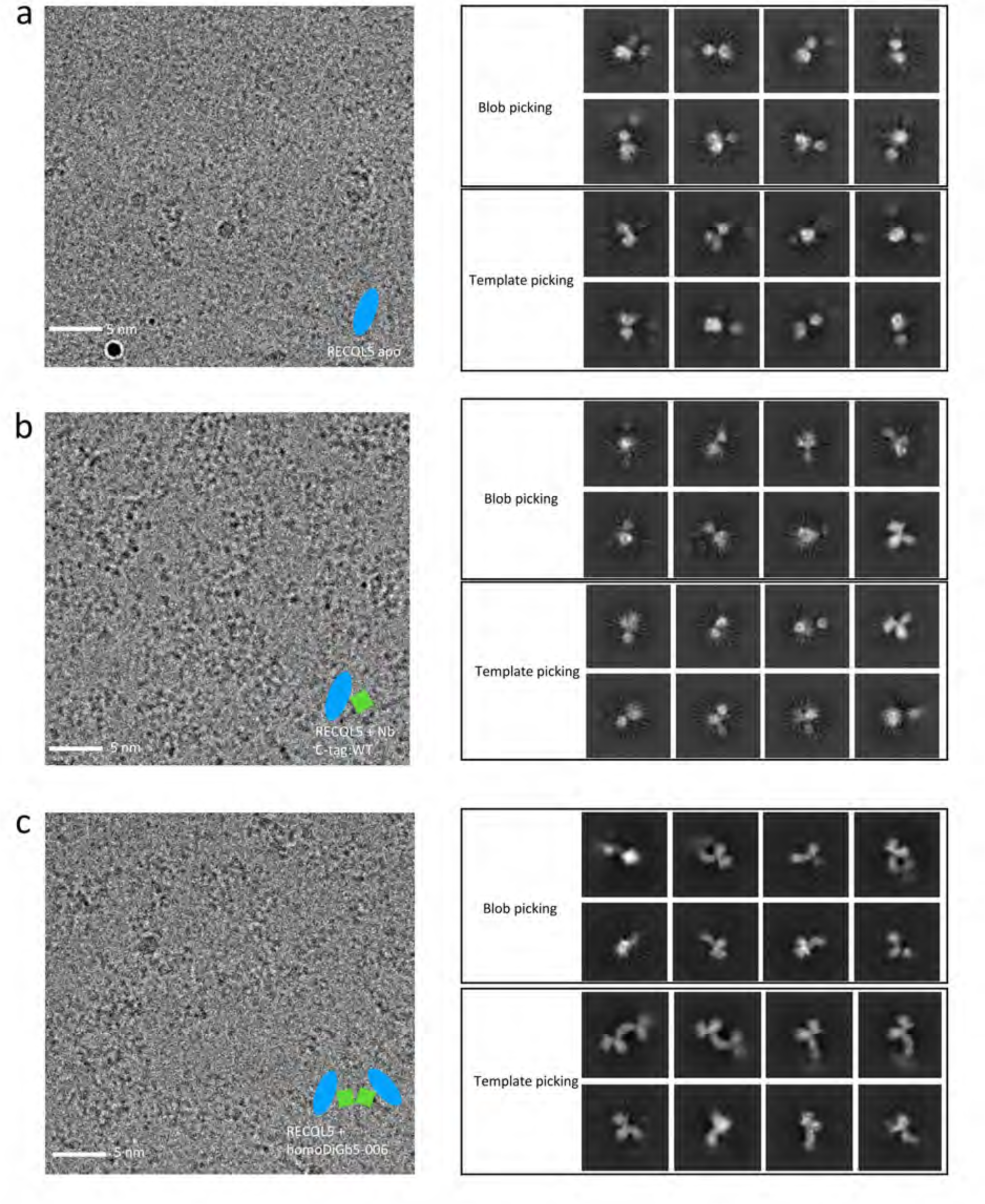
The homo Di-Gluebody improves the alignment of RECQL5. Cryo-EM data and 2D classification results for (a) RECQL5 alone, (b) RECQL5 in complex with nanobody, and (c) RECQL5 complexed with homo Di-Gluebody Gb5-006 with Data collected from 200 kV Glacios microscope. Exemplar raw images, schematics, and 2D classification of particles picked using blob and template-based picking are shown for each sample.

**Extended Data Figure 7.**
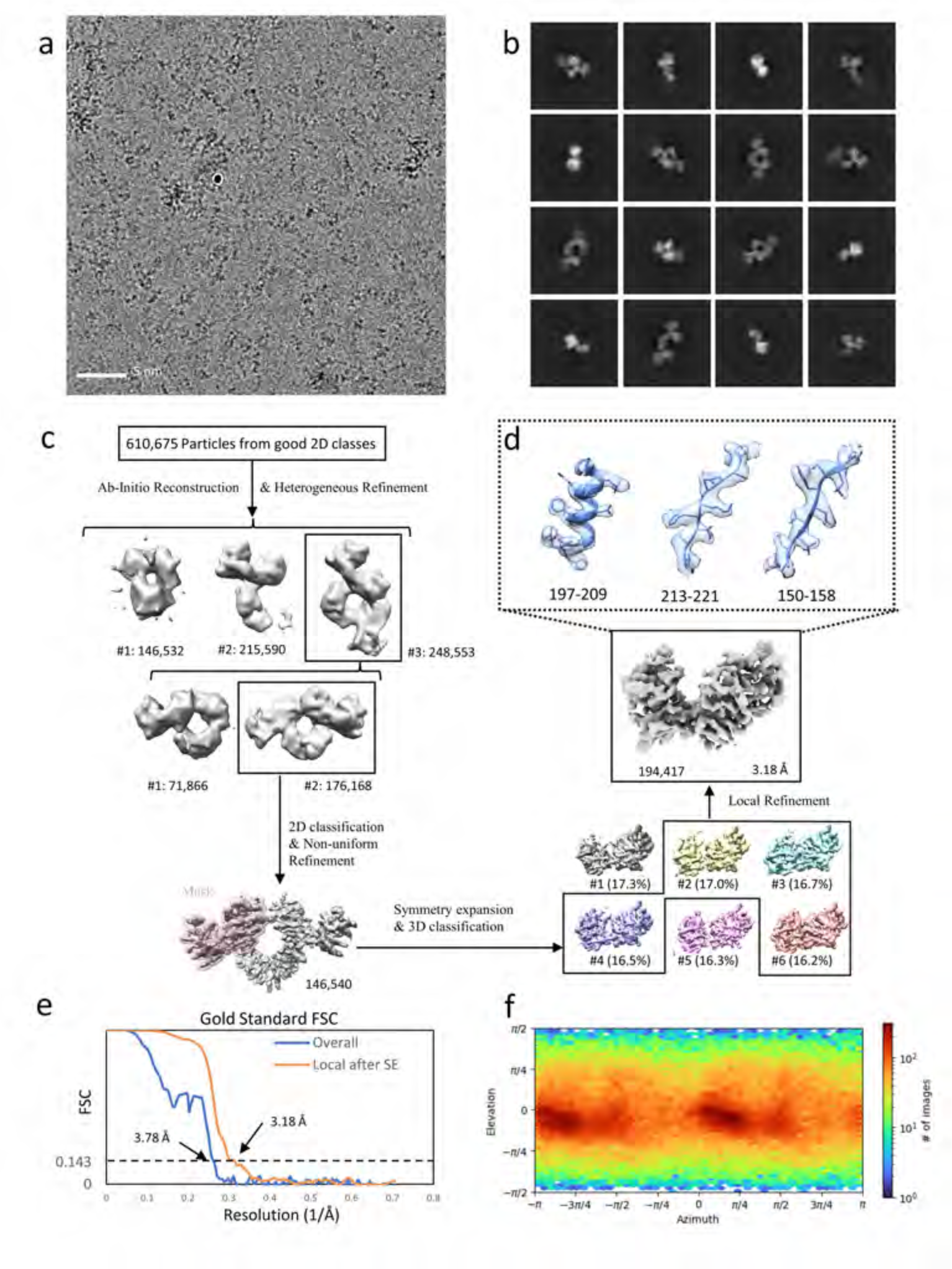
Cryo-EM structural determination of RECQL5 in complex with a homoDiGb5-006. (a) Exemplar raw micrographs. (b) 2D classification results of initial particles after Topaz picking. (c) Cryo-EM 3D reconstruction pipeline. (d) Map details of RECQL5’s residues 197-209, 213-221, and 150-158, containing helical, beta sheet, and loop secondary structures, respectively. (e) The Fourier Shell Correlation curves of the overall complex and locally refined RECQL5 after symmetry expansion (SE). (f) Particle orientation distribution of the final locally refined map.

**Extended Data Figure 8.**
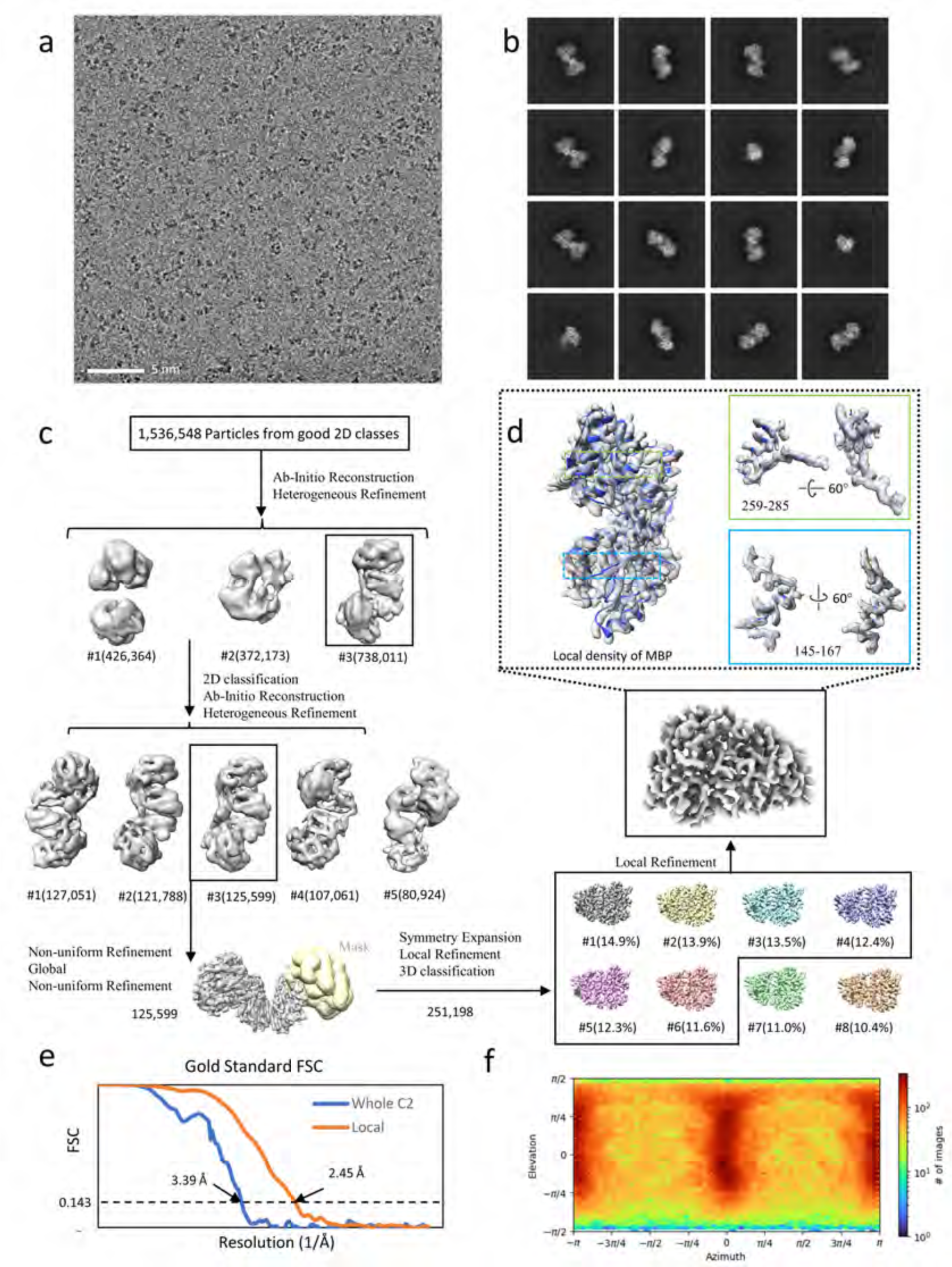
Cryo-EM structural determination of MBP in complex with a homoDiGbMBP. (a) Exemplar raw micrograph. (b) 2D classification of Topaz-picked particles. (c) Cryo-EM 3D reconstruction pipeline. (d) Cryo-EM map of MBP monomer after symmetry expansion, with map-and-model overlays for residues 145-167 and 259-285. (e) The Fourier Shell Correlation curves of the overall dimer complex and locally refined MBP after symmetry expansion (SE). (f) Particle orientation distribution of the final locally refined map.

**Extended Data Figure 9.**
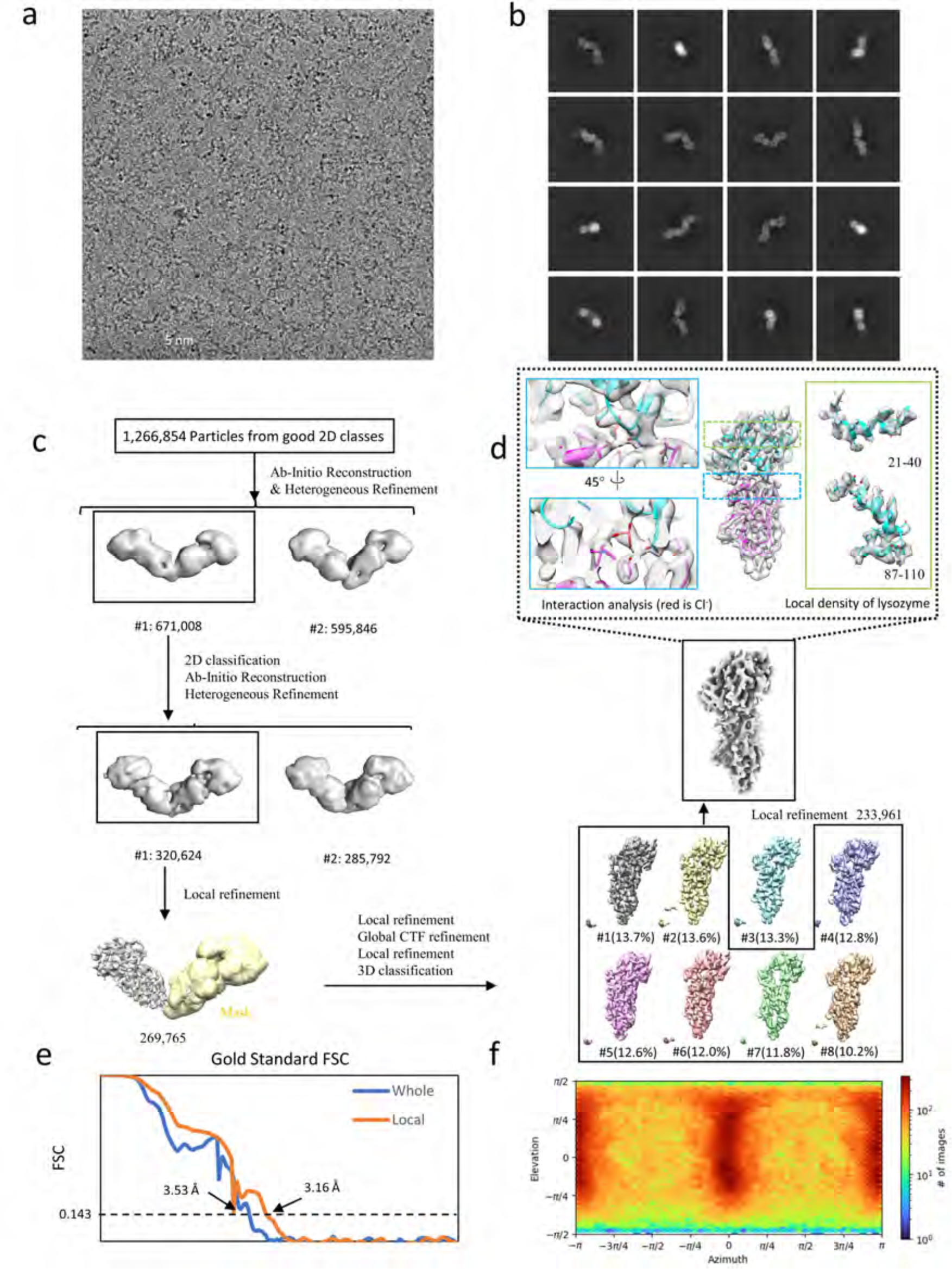
**Cryo-EM structural determination of lysozyme in complex with a homo Di-Gluebody GbLysozyme**. (a) Exemplar raw micrograph. (b) 2D classification of Topaz-picked particles. (c) Cryo-EM 3D reconstruction pipeline. (d) Cryo-EM map of lysozyme monomer after symmetry expansion, with map-and-model overlays for residues 21-40 and 87-110 and the binding site with the bound chloride. (e) The Fourier Shell Correlation curves of the overall dimer complex and locally refined lysozyme. (f) Particle orientation distribution of the final locally refined map.

**Extended Data Figure 10.**
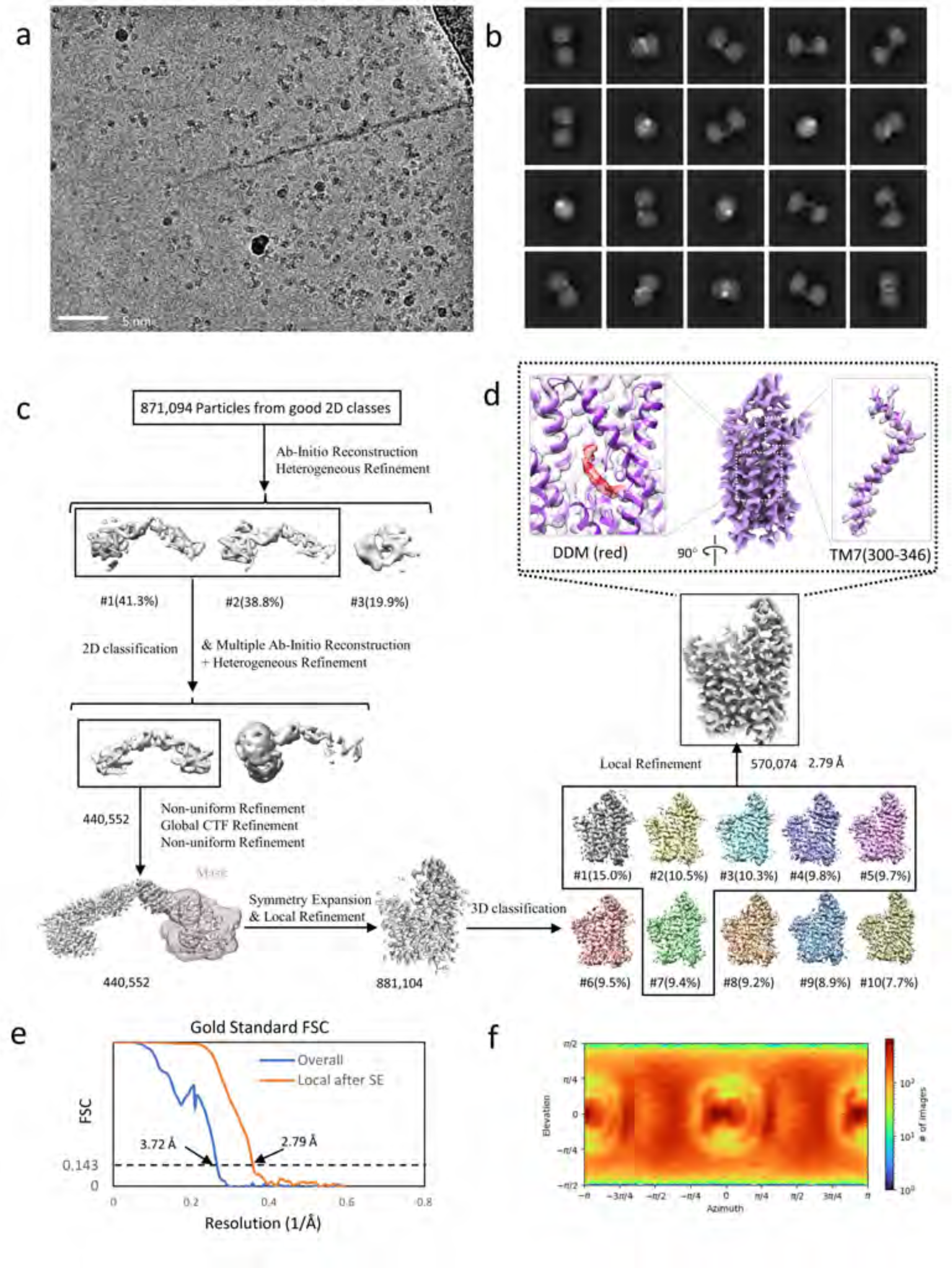
Cryo-EM structural determination of SPNS2 in complex with a homo Di-Gluebody GbD12. (a) Exemplar raw micrograph. (b) 2D classification of Topaz-picked particles. (c) Cryo-EM 3D reconstruction pipeline. (d) Cryo-EM map of SPNS2 monomer after symmetry expansion, with map-and-model overlays for TM7 residues 300-346 and central binding site with bound DDM. (e) The Fourier Shell Correlation curves of the overall dimer complex and locally refined SPNS2 after symmetry expansion (SE). (f) Particle orientation distribution of the final locally refined map.

**Extended Data Figure 11.**
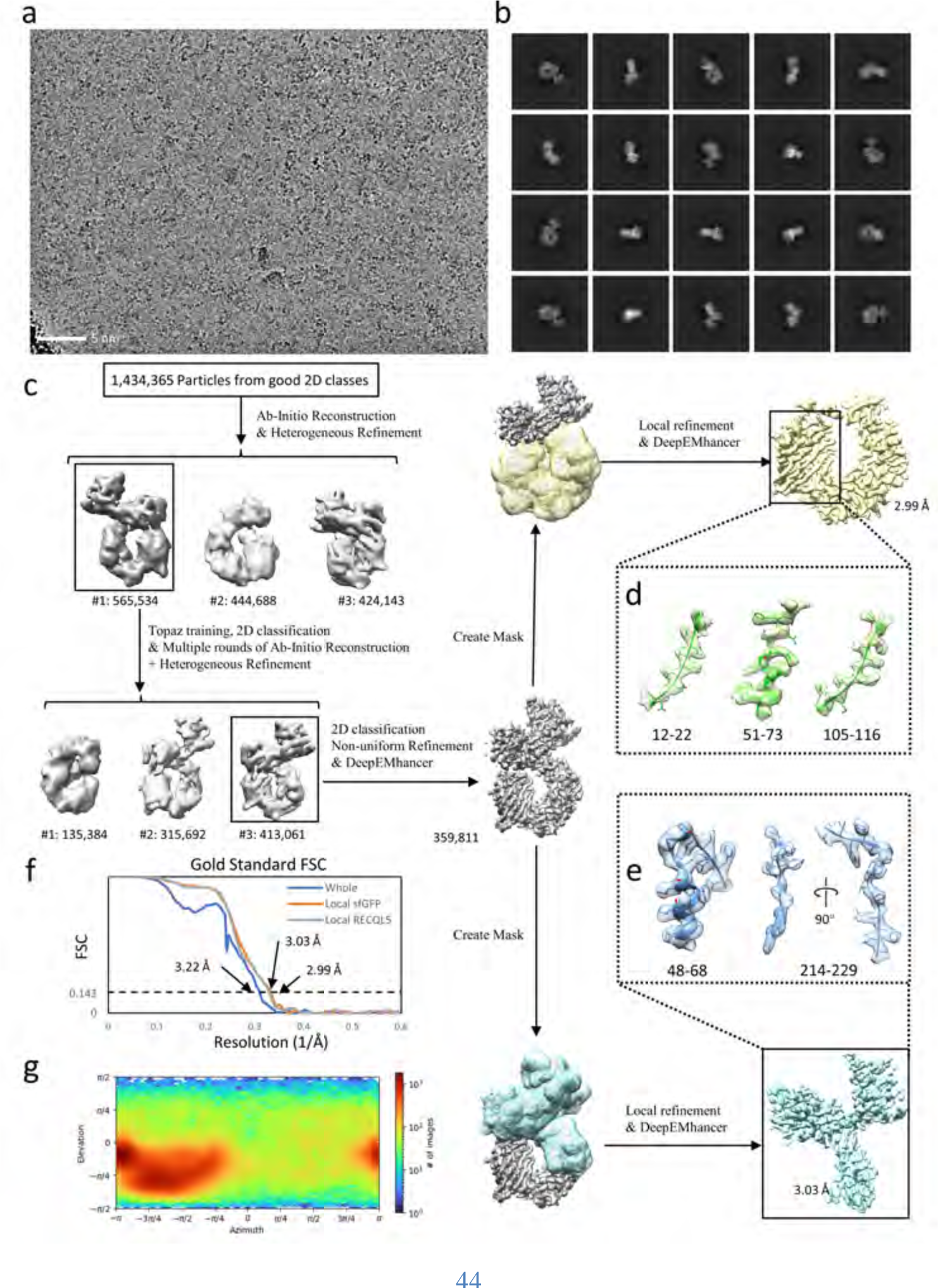
Cryo-EM structural determination of the RECQL5:heteroDiGb:sfGFP complex. (a) Exemplar raw micrograph. (b) 2D classification of particles selected by Topaz picking. (c) Cryo-EM 3D reconstruction pipeline. (d) Overall view of sfGFP after local refinement, and structural details of peripheral regions of the central helix (51-73) and two beta sheets (12-22 and 105-116). (e) Map details of RECQL5 sub regions of the helix (48-68) and the region containing a beta sheet and a loop (214-229). (f) The Fourier Shell Correlation curves of the overall complex, the locally refined map of sfGFP, and the locally refined map of RECQL5. (g) Particle orientation distribution of the overall complex before local refinement of individual targets.

**Extended Data Figure 12.**
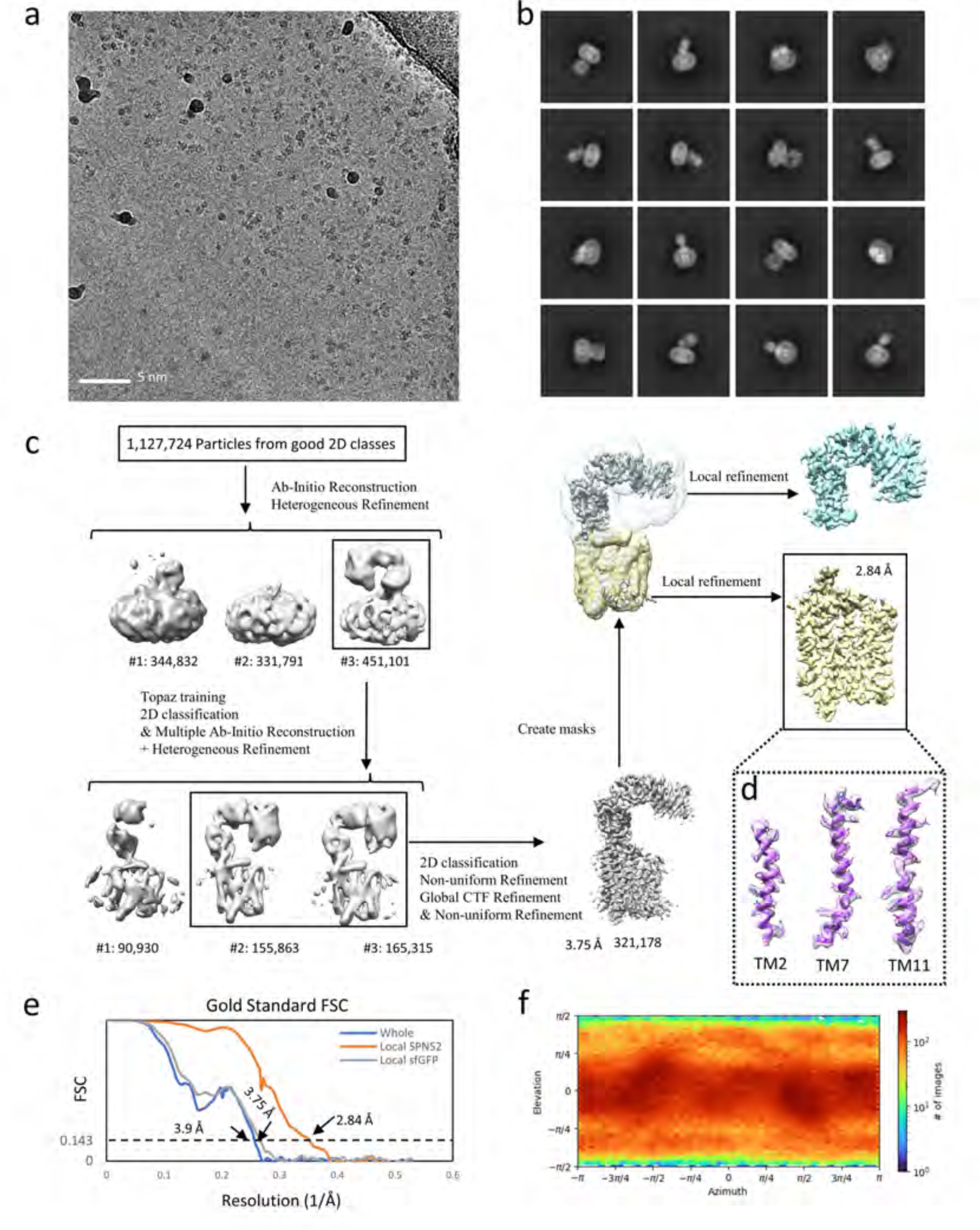
Cryo-EM structural determination of the SPNS2:heteroDiGb:sfGFP complex. (a) Exemplar raw micrograph. (b) 2D classification of particles selected by Topaz picking. (c) Cryo-EM 3D reconstruction pipeline. (d) Map and model of SPNS2’s transmembrane helices of TM2, TM7 and TM11. (e) The Fourier Shell Correlation curves of the overall complex before local refinement, the locally refined map of SPNS2, and the locally refined map of sfGFP. (f) Particle orientation distribution of the dimer complex before local refinement.

**Extended Data Figure 13.**
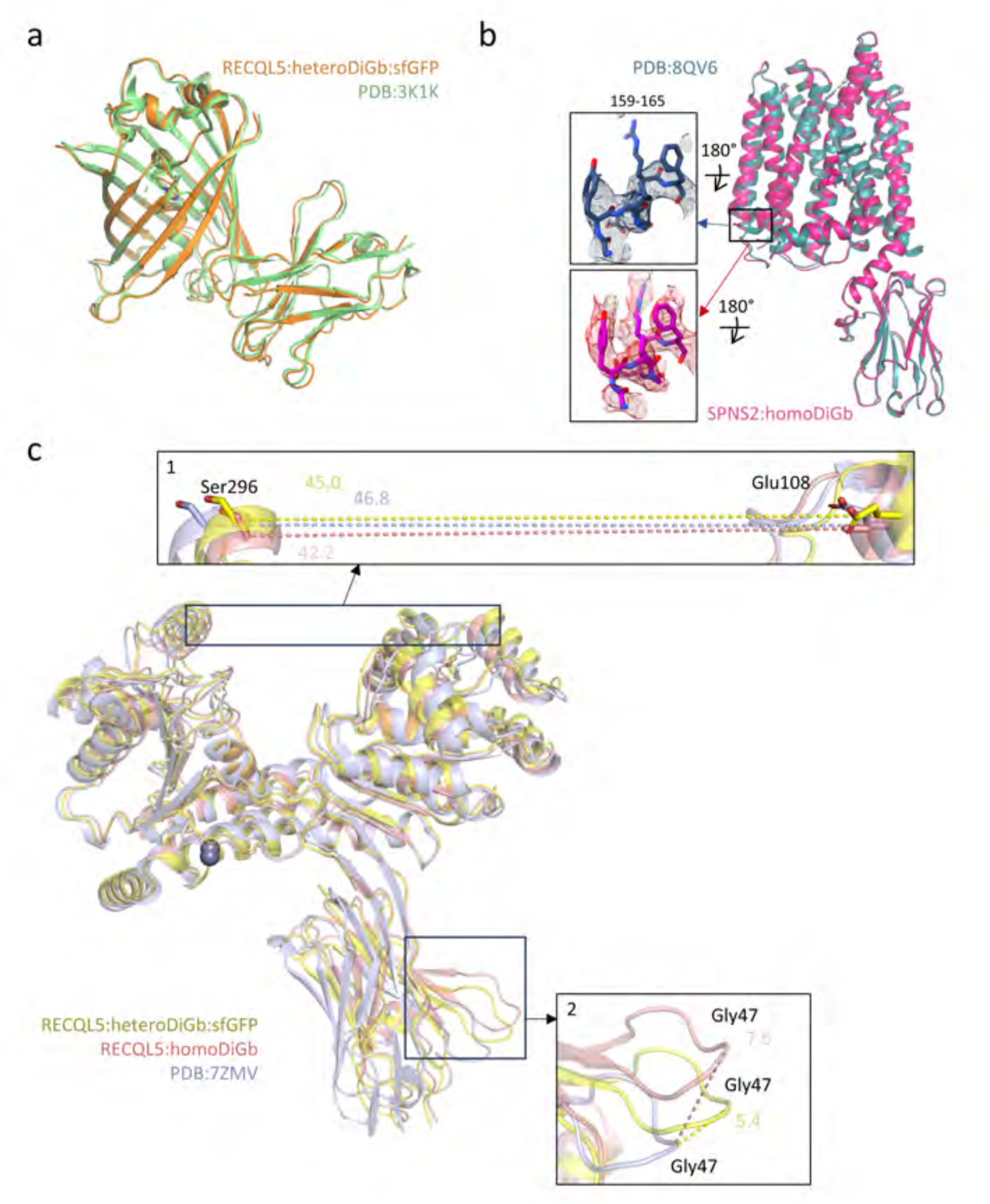
Comparison of the solved structures using Di-Gluebodies and previously determined structures. (a) Structural 3D alignment of the sfGFP:GbEnhancer complex in the RECQL5:sfGFP:hetDiGb(Gb5-006:GbEnhancer) complex structure and the published non-Gluebody structure (3K1K). (b) Structural 3D alignment of the SPNS2:GbD12 complex in the SPNS2:homoDiGb structure and the published SPSNS2:NbD12 structure (8QV6) with inset showing sidechain density improvements. (c) Structural 3D alignment of the RECQL5:Gb5-006 complex in the RECQL5:homoDiGb complex and RECQL5:heteroDiGb:sfGFP complex structures and the published RECQL5:Gb5-006 structure (7ZMV). Inset 1 distance between Cα atoms of RECQL5’s Glu108 in D1 domain and Ser296 in D2 domain. Inset 2 shows the distance comparison between Cα atoms of Gly47 of the Gluebodies. Distances are shown in Å.

**Extended Data Figure 14.**
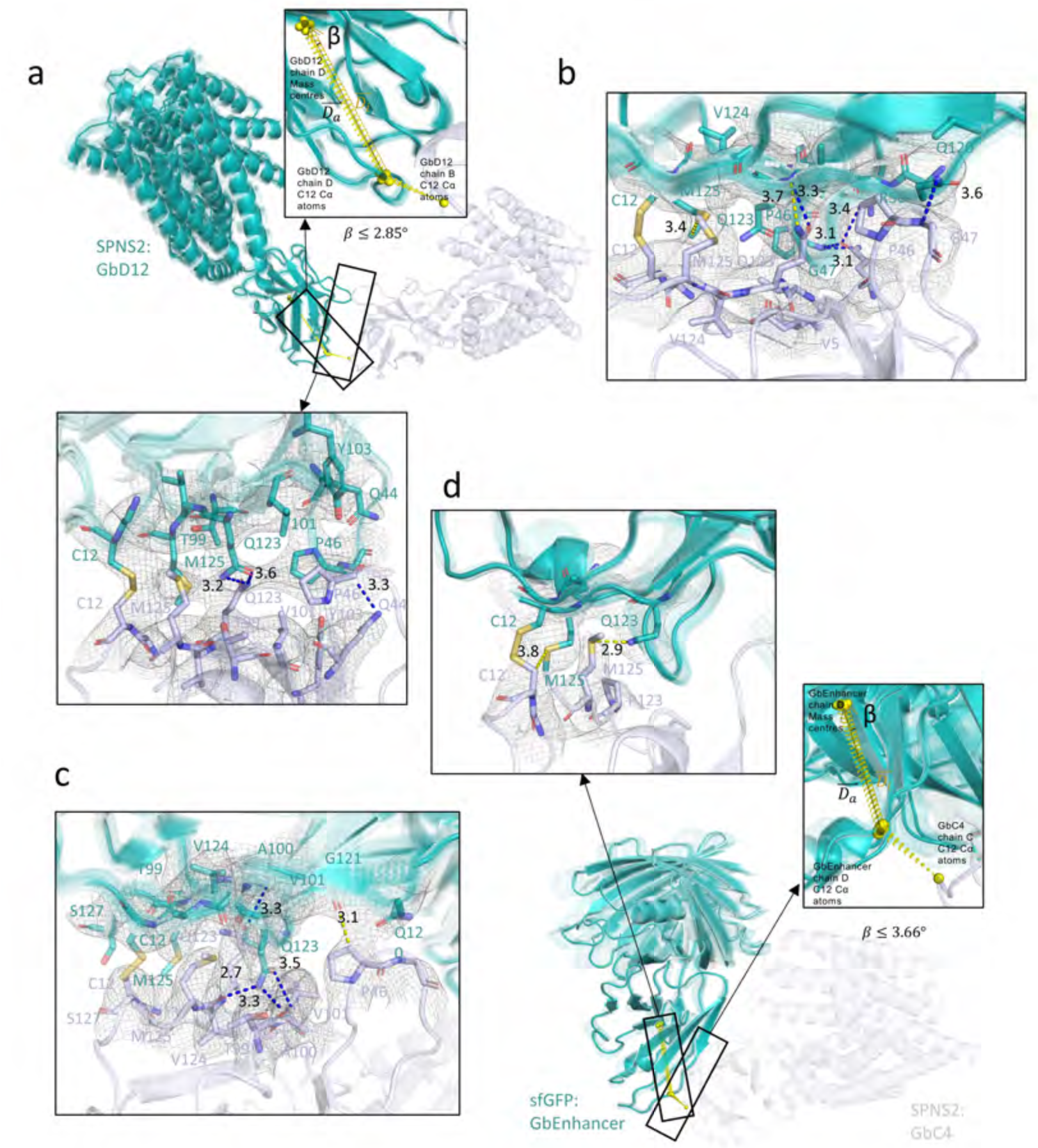
3D variability analyses of Di-Gluebody structures. The 60 models of each structure are overlaid and aligned using the static Gluebody. Static and moving Gluebodies are shown in light blue and teal, respectively. The left insets indicate the reference points (yellow spheres), vectors (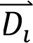 shown as a yellow arrow) and angle β for the moving Gluebody used in the analyses. 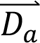 shown as a black arrow is the average vector of 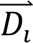. The right insets show the Di-Gluebody interfaces with the local density shown in mesh, interacting residues shown in stick, and hydrogen bonds shown in blue dashes. Distances are shown in Å. The analyses include (a) SPNS2:homoDiGb (b) RECQL5:homoDiGb (whole map and wobbling angle shown in Figure 3a) (c) RECQL5:heteroDiGb:sfGFP (whole map and wobbling angle shown in Figure 3c) and (d) SPNS2:heteroDiGb:sfGFP.

**Extended Data Figure 15.**
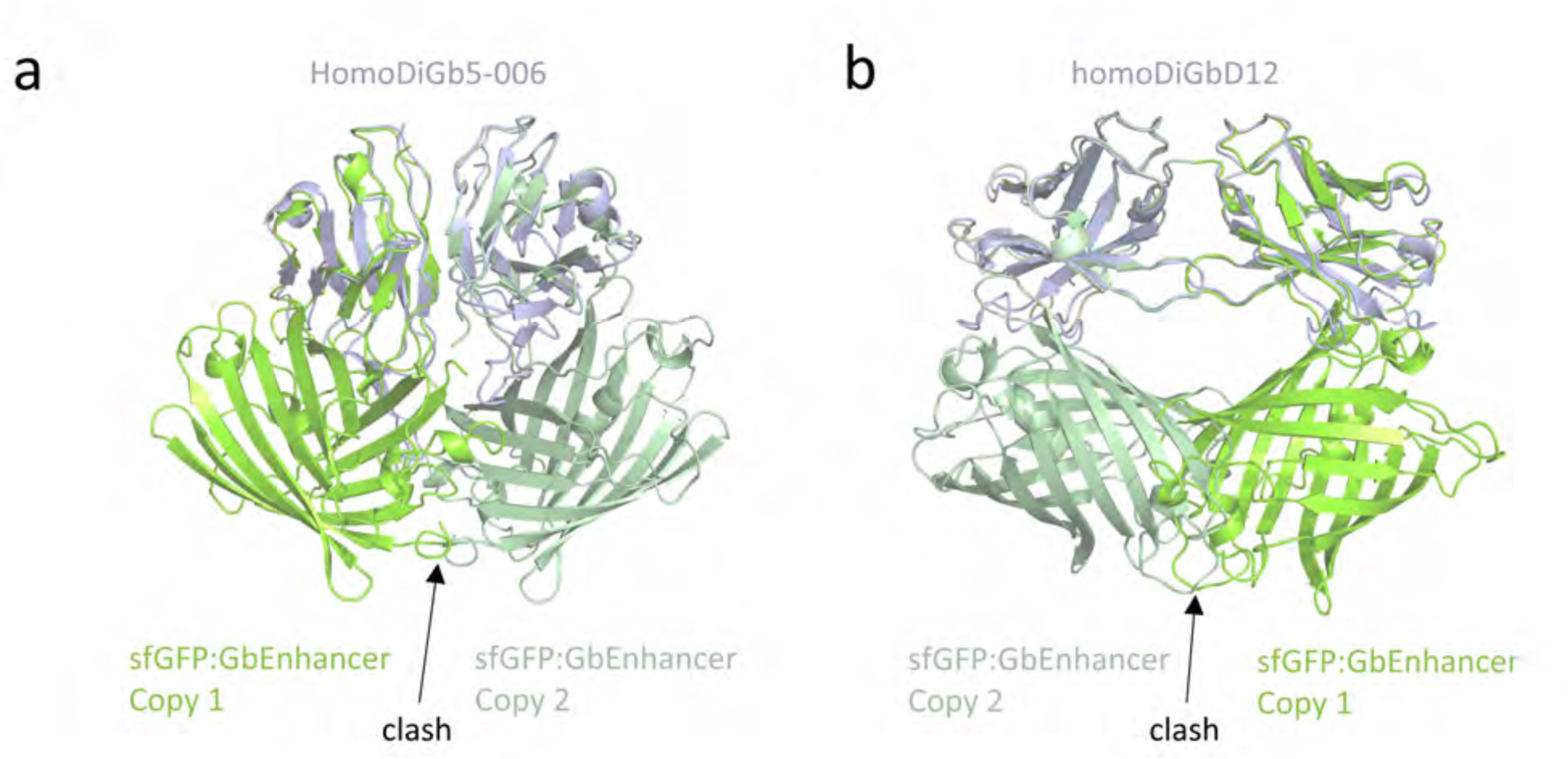
The GbEnhancer homo Di-Gluebody is expected to introduce clash when the target GFP is attached. (a) The GFP:NbEnhancer structure (PDB: 3K1K) modeled on the homo Di-Gluebody Gb5-006. (b) The 3K1K structure modeled on the homo Di-Gluebody GbD12. Clash is indicated by the arrow.

**Extended Data Figure 16.**
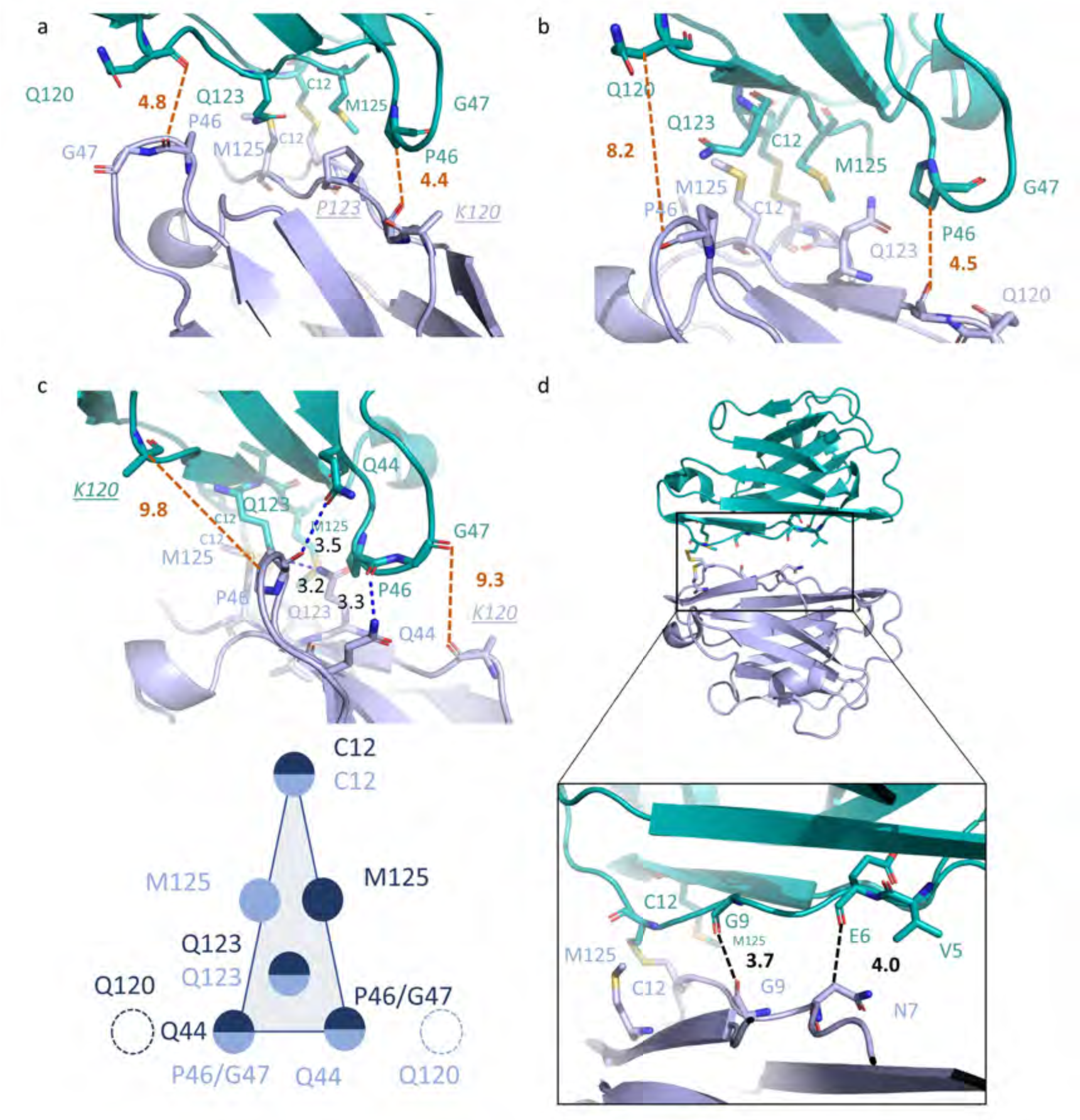
The DiGb interfaces are influenced by the interacting residue side chains and driven by the disulfide. Two Gluebodies are shown in lightblue and teal. Typical Cys-Cys interfaces were present in structures of (a) SPNS2:heteroDiGb:sfGFP and (b) Lysozyme:homoDiGb. Narrower Cys-Cys interface was present in the structure of (c) SPNS2:homoDiGb with P46/G47 interacting with Q44 instead of residue 120, which has a Lysine instead of Glutamine at this position. Interface cartoon is drawn below. The MBP:homoDiGb presents an even more unique DiGb interface that is chemically driven by the disulfide and assisted by the first beta-sheets of the Gluebodies. Blue dashes show hydrogen bonds, and dashes show the minimum distances (not counting hydrogen atoms) between the P46/G47 and Q/K120 pairs, and black ones show distances between two atoms. Atypical residue types are shown in italics underlined.

**Extended Data Table 1.**
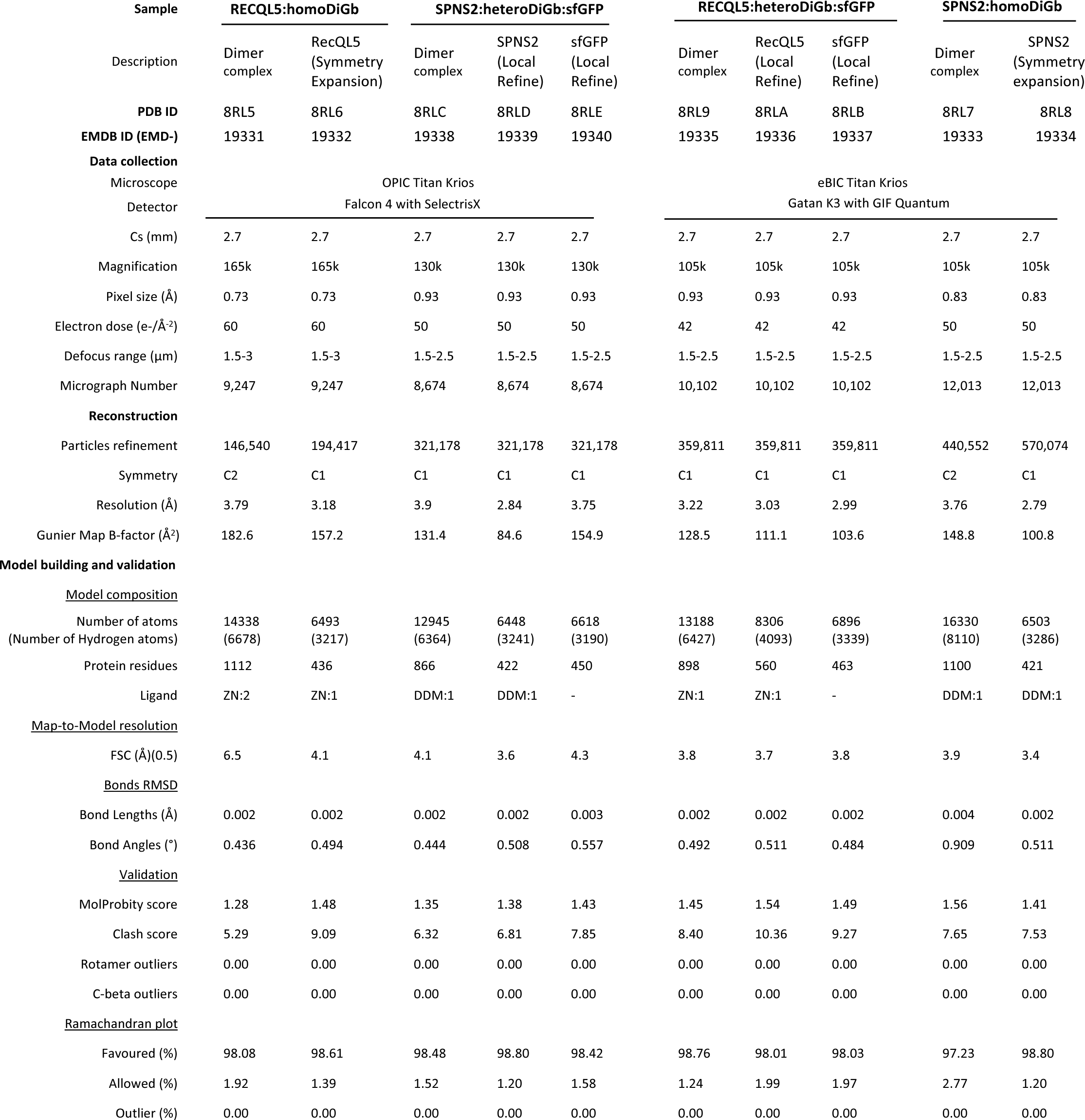
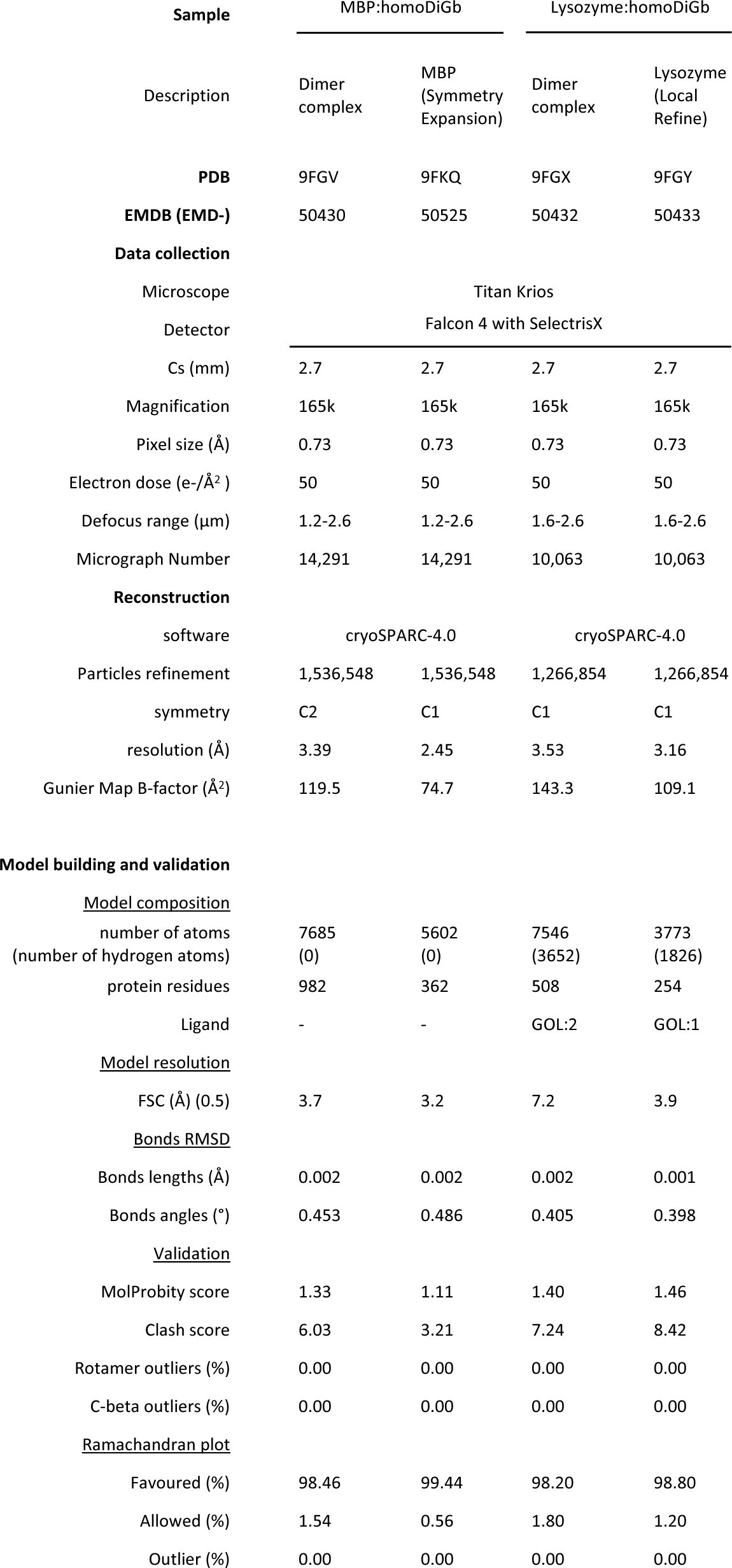
Cryo-EM Data Collection, Processing, Refinement and Model Building and Validation.

**Extended Data Table 2.** 3D Variability Analyses Summary.

| Structure | No. of interacting residues (<4Å) in the Gluebody interface | No. of hydrogen bonds in the Gluebody interface | Moving Gluebody chain ID | Static Gluebody chain ID | Intra-Glebody angle (°) |
| --- | --- | --- | --- | --- | --- |
| RECQL5:homoDiGb | 28 | 5 | L (Gb5-006) | K (Gb5-006) | 71.32 |
| SPNS2:homoDiGb | 21 | 3 | D (GbD12) | B (GbD12) | 100.70 |
| SPNS2:heteroDiGb:sfGFP | 8 | 0 | D (GbEnhancer) | C (GbC4) | 96.34 |
| RECQL5:heteroDiGb:sfGFP | 26 | 4 | D (GbEnhancer) | K (G5-006) | 82.00 |

b
| Structure | Max $\overrightarrow{D_a}$ wobble angle $\beta$ (°) | 75 quartile $\overrightarrow{D_a}$ wobble angle $\beta$ (°) | Median $\overrightarrow{D_a}$ wobble angle $\beta$ (°) |
| --- | --- | --- | --- |
| RECQL5:homoDiGb | 4.04 | 1.22 | 0.78 |
| SPNS2:homoDiGb | 2.85 | 1.21 | 0.47 |
| SPNS2:heteroDiGb:sfGFP | 3.66 | 0.61 | 0.40 |
| RECQL5:heteroDiGb:sfGFP | 2.01 | 1.24 | 0.84 |

## Notes

### Competing Interest Statement

The authors have declared no competing interest.

