## Supplementary Methods and Discussion for "Di-Gluebodies as covalently-rigidified, modular protein assemblies enable simultaneous determination of high-resolution, low-size, cryo-EM structures"

### SUPPLEMENTARY INFORMATION

### Supplementary Methods

#### General Experimental Procedures

Unless otherwise noted, chemical reagents, media, and Escherichia coli cell stocks were obtained from commercial suppliers (Sigma-Aldrich, Fluorochem, Carbosynth, VWR, Alfa Aesar, Fisher Scientific) and used without further purification.

#### Protein Concentration Measurement

Protein concentration was measured using the Implen C40 NanoPhotometer UV/Vis Spectrophotometer, using the custom function using the molecular weight and Ext. coefficient, as calculated by the ProtParam tool, Expasy.

#### Protein Mass Spectrometry

Protein samples were analysed on Waters Xevo G2-XS QTof mass spectrometers equipped with a Waters Acquity UPLC. Separation was achieved using a Thermo Scientific ProSwift RP-2H monolithic column (4.6 mm × 50 mm) using water + 0.1% formic acid (solvent A) and acetonitrile + 0.1% formic acid (solvent B) as mobile phase at a flow rate of 0.4 ml/min and running a 5-min linear gradient as follows: 5% solvent B for 0.5 min, 5 to 95% solvent B over 3.0 min, 95 to 5% solvent B over 0.5 min, and 5% solvent B for 1 min. Spectra were deconvoluted using MassLynx 4.1 (Waters) and the "MaxEnt1" deconvolution algorithm with the following settings: resolution: 1.0 Da per channel; damage model: uniform Gaussian; width at half height: 0.4 Da; minimum intensity ratios: 33% (left) and 33% (right); and iterate to convergence. Conversions were calculated from peak intensities.

#### Protein Constructs

##### Anti-SPNS2 Gluebody GbH12

```
        10         20         30         40         50         60
SMAQVQLVEN GGGCVKPGGS LRLSCAASGS RFSDNTMAWY RQAPRKQREL VARIPMGGRP

        70         80         90        100        110
MYADSVKGRF TISRDNAENT VYLQMNSLKP DDTAVYYCNA VTYGLESYWG KGTQVMVS
```

**Molecular weight:** 12970.71 g/mol

**Ext. coefficient:** 21430 $M^{-1}$ $cm^{-1}$

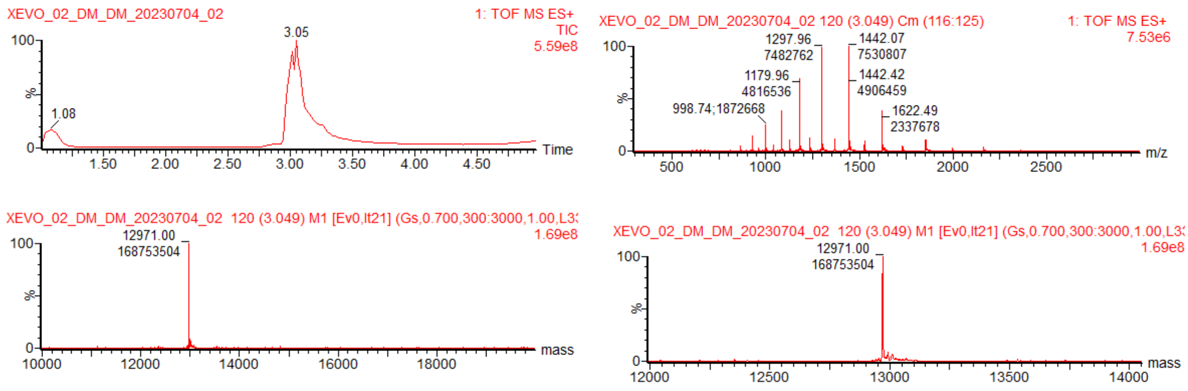


Figure 1. Calculated mass of Gluebody GbH12: 12969, Observed mass: 12971

#### Anti-GFP Gluebody

```
         10         20         30         40         50         60
SQVQLVENGG ACVKPGGSLR LSCAASGFPV NRYSMRWYRQ APGKEREWVA GMSSAGDRSS

         70         80         90        100        110
YEDSVKGRFT ISRDDARNTV YLQMNSLKPE DTAVYYCNVN VGFEYWGQGT QVMVS
```

**Molecular weight:** 12763.23 g/mol

**Ext. coefficient:** 26930 $M^{-1}$ $cm^{-1}$

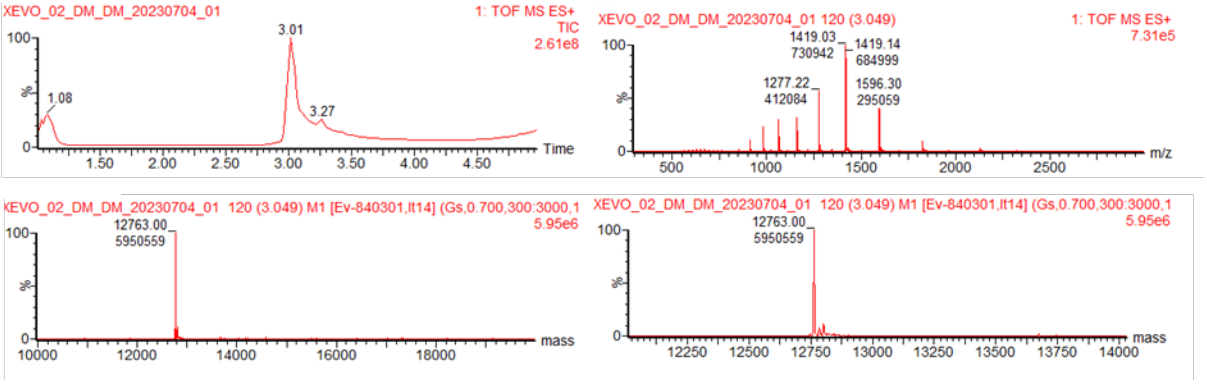


Figure 2. Calculated mass of Gluebody GbEnhancer: 12761, Observed mass: 12763

Anti-RECQL5 *Gluebody* Gb5-006

```
         10         20         30         40         50         60
SMAQVQLVEN GGGCVKAGGS LRLSCAASGS IFSINRMTWY RQAPGKEREW VAAITSGGST

         70         80         90        100        110        120
NYADSVKGRF TISRDNAENT VYLQMNSLKP EDTAVYYCEA YGTYTLAPTG EGEYDDYWGQ

GTQVMVS
```

**Molecular weight:** 13765.23 g/mol

**Ext. coefficient:** 29910 $M^{-1}$ $cm^{-1}$

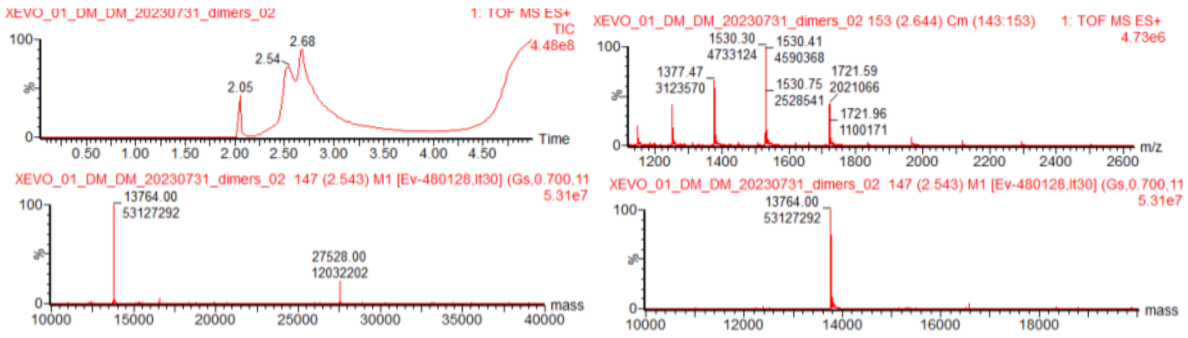


Figure 3. Calculated mass of Gluebody Gb5-006: 13763, Observed mass: 13764

#### Anti-TUT4 Gluebody GbS2A4

```
         10         20         30         40         50         60
SMAQVQLVEN GGGCVKAGGS LRLSCAASGT IFTYFVMGWY RRAPGKEREL VAGITLGGTT

         70         80         90        100        110        120
YYADSVKGRF TISRDNAKNT VYLQMNSLKP EDTAVYYCAA WVEYPRRYVY WGQGTQVMVS
```

**Molecular weight:** 13262.11 g/mol

**Ext. coefficient:** 31400 $M^{-1}$ $cm^{-1}$

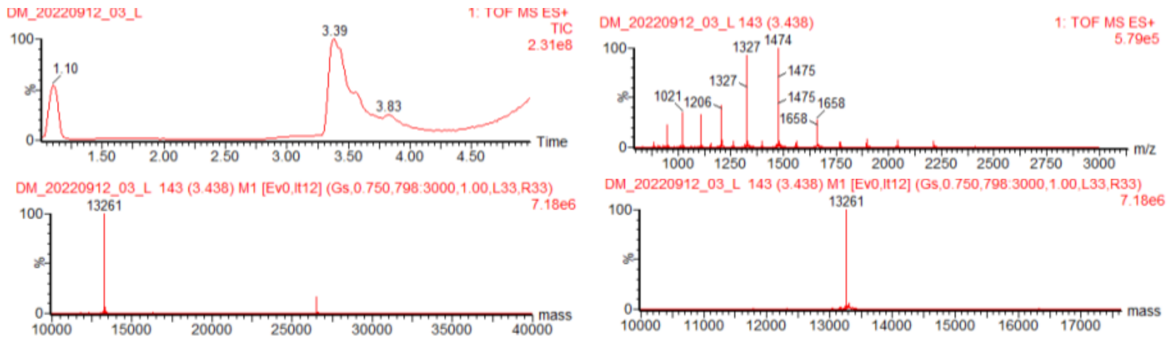


Figure 4. Calculated mass of Gluebody GbS2A4: 13260, Observed mass: 13261

#### Anti-MBP Gluebody GbMBP

```
         10         20         30         40         50         60
SQVQLVENGG GCVKAGGSLR LSCVASGDIK YISYLGWFRQ APGKEREGVA ALYTSTGRTY

         70         80         90        100        110        120
YADSVKGRFT VSLDNAKNTV YLQMNSLKPE DTALYYCAAA EWGSQSPLTQ WFYRYWGQGT


QVMVS
```

**Molecular weight:** 13776.48 g/mol

**Ext. coefficient:** 36900 $M^{-1}$ $cm^{-1}$

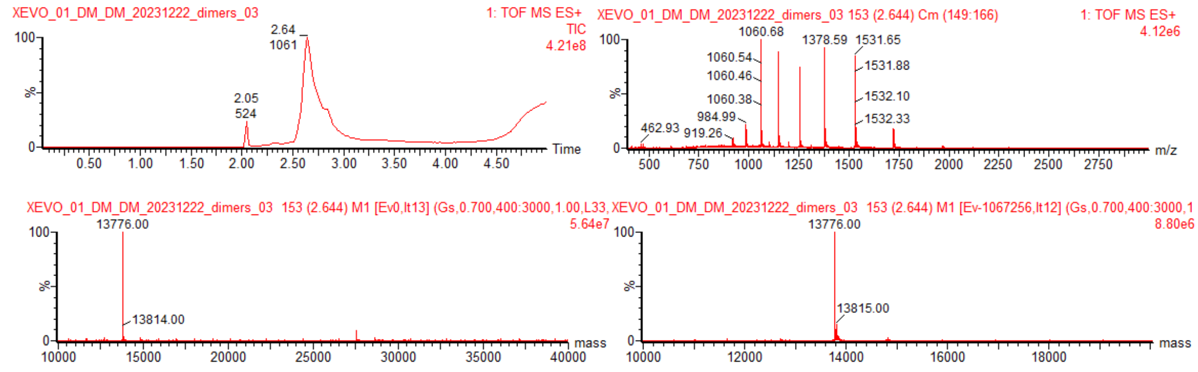


Figure 5. Calculated mass of Gluebody GbMBP: 13774, Observed mass: 13776

#### Anti-SPNS2 Gluebody GbD12

```
10         20         30         40         50         60
SMAQVQLVEN GGGCVKAGGS LRLSCAASGR LLSWYDMAWF RQAPGKEREF VAAVTSTGAG

        70         80         90        100        110        120
THYVDSVKGR FTISRVNAEN TMYLQMNSLK PEDTAVYYCA AANTRLTALS LRTTTGSWAY

       130
WGKGTQVMVS
```

**Molecular weight:** 14044.94 g/mol

**Ext. coefficient:** 30940 $M^{-1}$ $cm^{-1}$

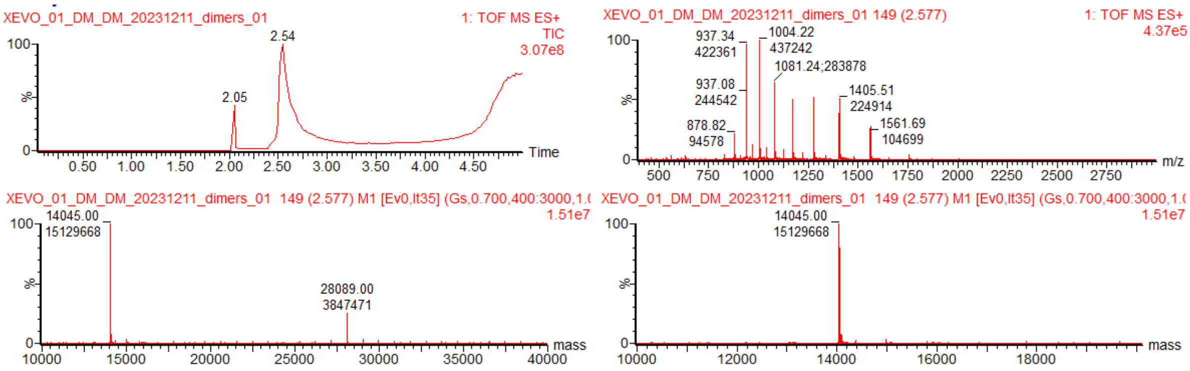


Figure 6. Calculated mass of Gluebody GbD12: 14043, Observed mass: 14045

#### Anti-SPNS2 Gluebody GbC4

```
        10         20         30         40         50         60
SQGQLVENGG GCVKAGGSLR LSCAASQGTL SNLVTGWFRR APGKEREFVA NIGRDGLTVY

        70         80         90        100        110        120
SNSVKGRFTI SRDRAKNTVY LQMDSLKPED TAVYYCAGRL SRFPGEYDYW SKGTPVMVSS

SQ
```

**Molecular weight:** 13268.90 g/mol

**Ext. coefficient:** 19940 $M^{-1}$ $cm^{-1}$

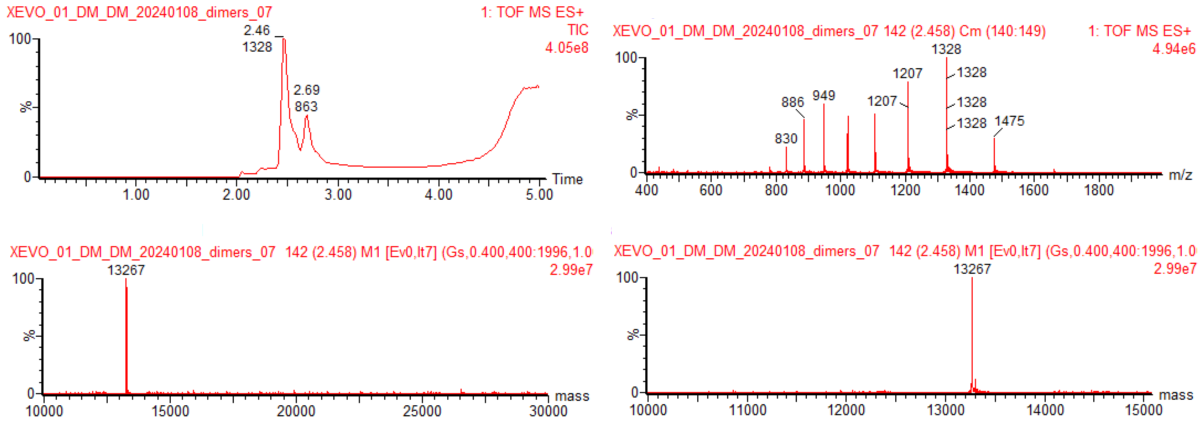


Figure 7. Calculated mass of Gluebody F9 12Cys: 13267, Observed mass: 13267

#### Anti-SARS-COV2-SpikeRBD Gluebody GbRBD1

```
        10         20         30         40         50         60
SQVQLVENGG GCMKAGGSLR LSCAVSGRTF STAAMGWFRQ APGKEREFVA AIRWSGGSAY

        70         80         90        100        110        120
YADSVKGRFT ISRDKAENTV YLQMNSLKYE DTAVYYCAQT RVTRSLLSDY ATWPYDYWGQ

GTQVMVS
```

**Molecular weight:** 14072.81 g/mol

**Ext. coefficient:** 35410 $M^{-1}$ $cm^{-1}$

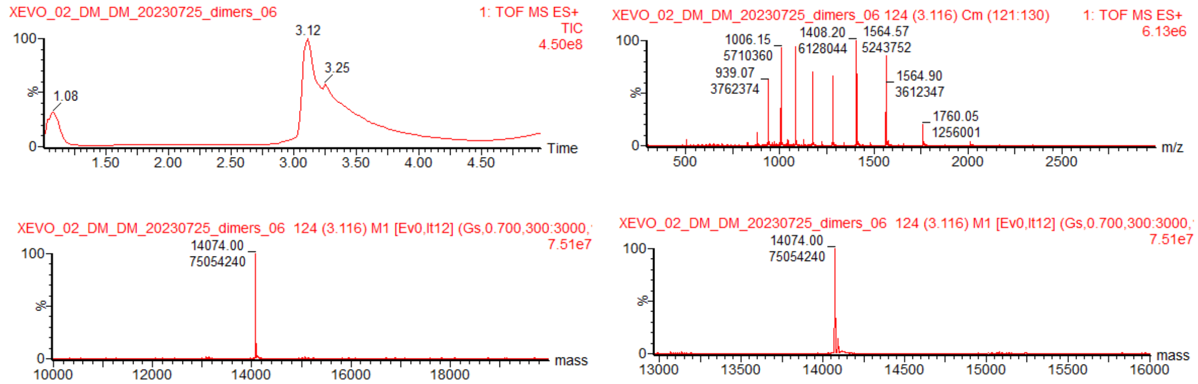

Figure 8. Calculated mass of Gluebody GbRBD1: 14071, Observed mass: 14074

#### Anti-SARS-COV2-SpikeRBD Gluebody GbRBD3

```
        10         20         30         40         50         60
HVQLVENGGG CVKAGGSLRL SCATSGRTFS TYRMSWFRQA PGKEREFVAT IIWSVGSTHY

        70         80         90        100        110        120
ADSVKGRFTI SRDNAENMVY LQMNSLKPED TAVYYCAAQR SDSSSWGYED DYDYWGQGTQ
```

VMVSS

**Molecular weight:** 13950.42

**Ext. coefficient:** 33920 $M^{-1}$ $cm^{-1}$

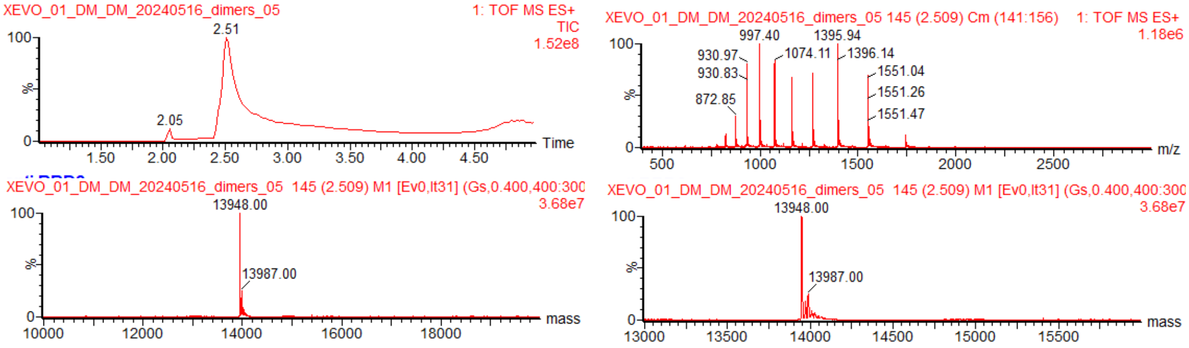


Figure 9. Calculated mass of Gluebody GbRBD3: 13948, Observed mass: 13948

**Anti-SARS-COV2-SpikeRBD Gluebody GbRBD6**

```
        10         20         30         40         50         60
SQVQLVENGG GCVKAGGSLR LACIASGRTF HSYVMAWFRQ APGKEREFVA AISWSSTPTY

        70         80         90        100        110        120
YGESVKGRFT ISRDNAENTV YLQMNRLKPE DTAVYFCAAD RGESYYYTRP TEYEFWGQGT
```

QVMVS

**Molecular weight:** 14038.69 g/mol

**Ext. coefficient:** 29910 $M^{-1}$ $cm^{-1}$

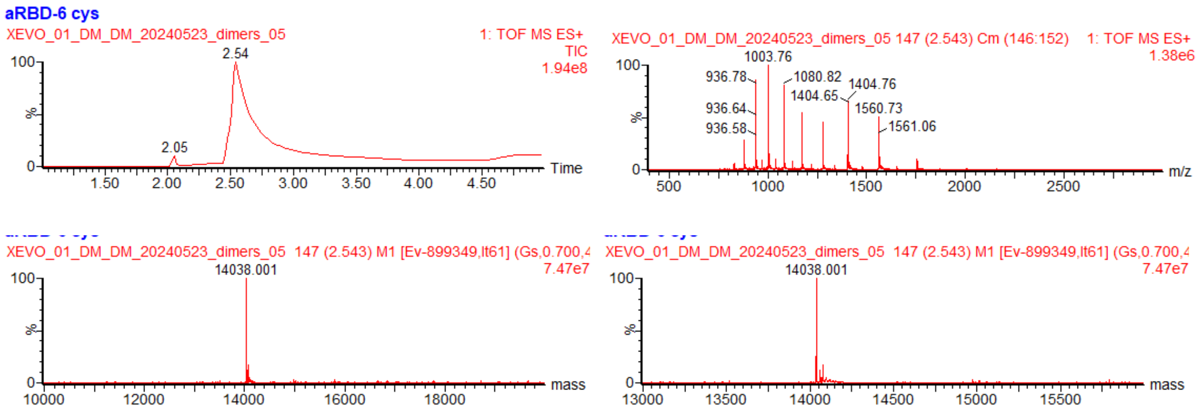


Figure 10. Calculated mass of Gluebody GbRBD6: 14037, Observed mass: 14038

#### Anti-Lysozyme Gluebody GbLys

```
         10         20         30         40         50         60
SDVQLVENGG GCVKAGGSLR LSCAASGSTD SIEYMTWFRQ APGKAREGVA ALYTHTGNTY

         70         80         90        100        110        120
YTDSVKGRFT ISQDKAKNMA YLRMDSVKSE DTAIYTCGAT RKYVPVRFAL DQSSYDYWGQ

        130
GTQVMVSSSA G
```

**Molecular weight:** 14155.76 g/mol
**Ext. coefficient:** 24410 $M^{-1}$ $cm^{-1}$

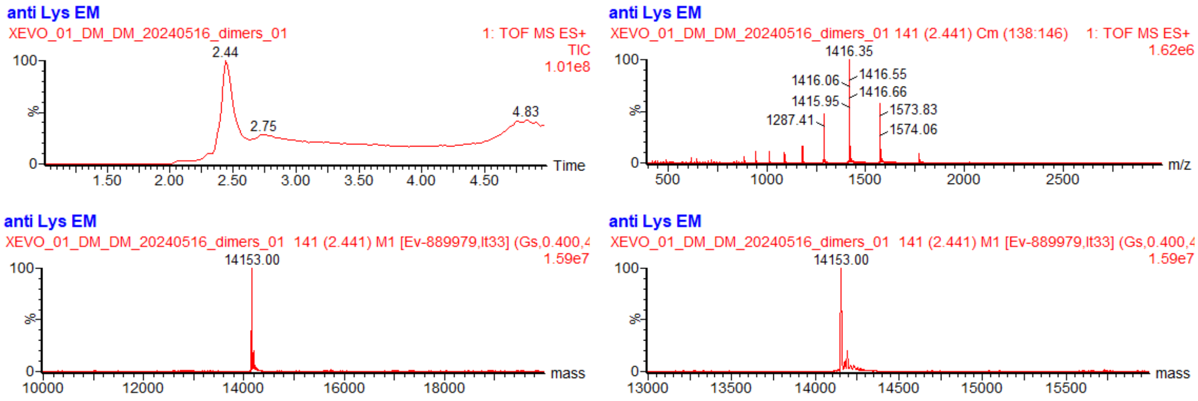


Figure 11. Calculated mass of Gluebody GbLys: 14154, Observed mass: 14153

#### Anti-HIV Gluebody GbHIV

```
        10         20         30         40         50         60
SDVQLQENGG GCVKAGGSLR LSCAASGSIS RFNAMGWWRQ APGKEREFVA RIVKGFDPVL

        70         80         90        100        110        120
ADSVKGRFTI SIDSAENTLA LQMNRLKPED TAVYYCFAAL DTAYWGQGTQ VMVSSAAADY

       130
KPGGGKPGGE PEA
```

**Molecular weight:** 14067.81 g/mol
**Ext. coefficient:** 22460 $M^{-1}$ $cm^{-1}$

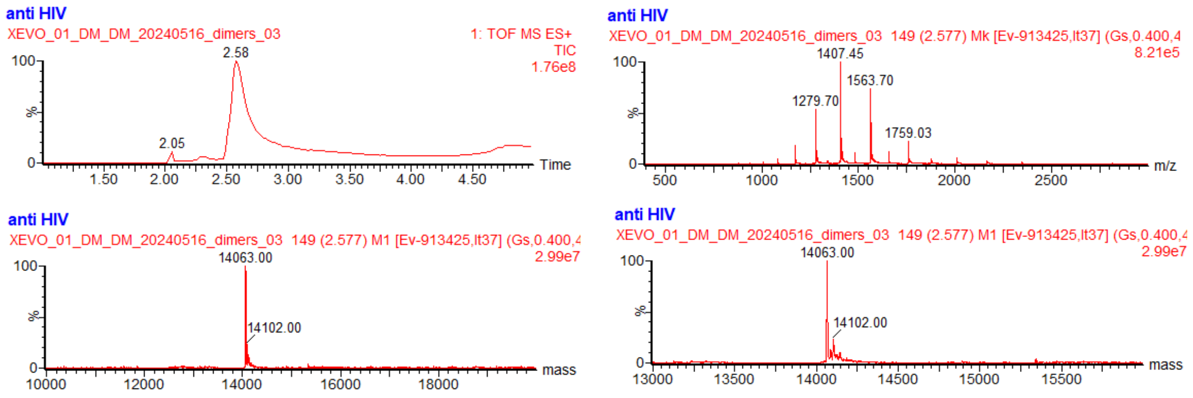


Figure 12. Calculated mass of Gluebody GbHIV: 14066, Observed mass: 14063

### General Protocol for the Chemical Dimerization of homo Di-Gluebody

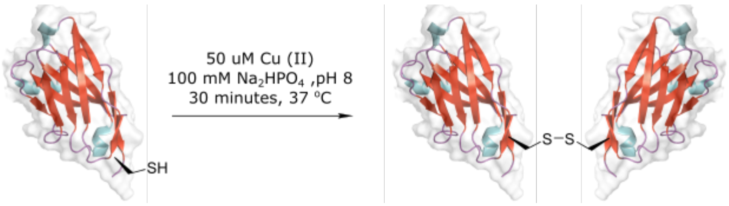


To generate homo Di-Gluebodies, purified Gluebody at 2 mg/ml in 50 mM $Na_2HPO_4$, pH 8, was treated with 50 uM of Cu (II) acetate, and the resulting solution was incubated at 37 °C temperature for 30 minutes. Conversion was monitored by intact mass spectrometry.

#### GbS2A4 homoDiGb

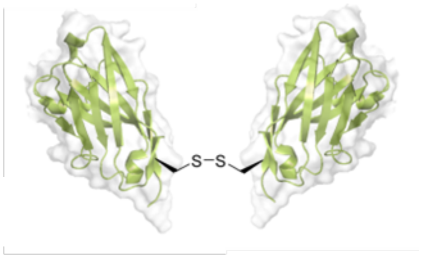


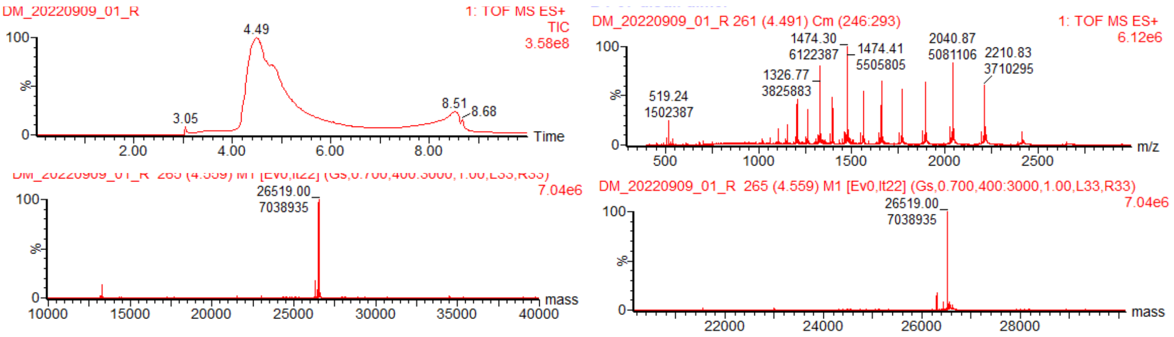


Figure 13. Calculated mass of GbS2A4 homoDiGb: 26518, Observed mass: 26519

#### Gb5-006 homoDiGb

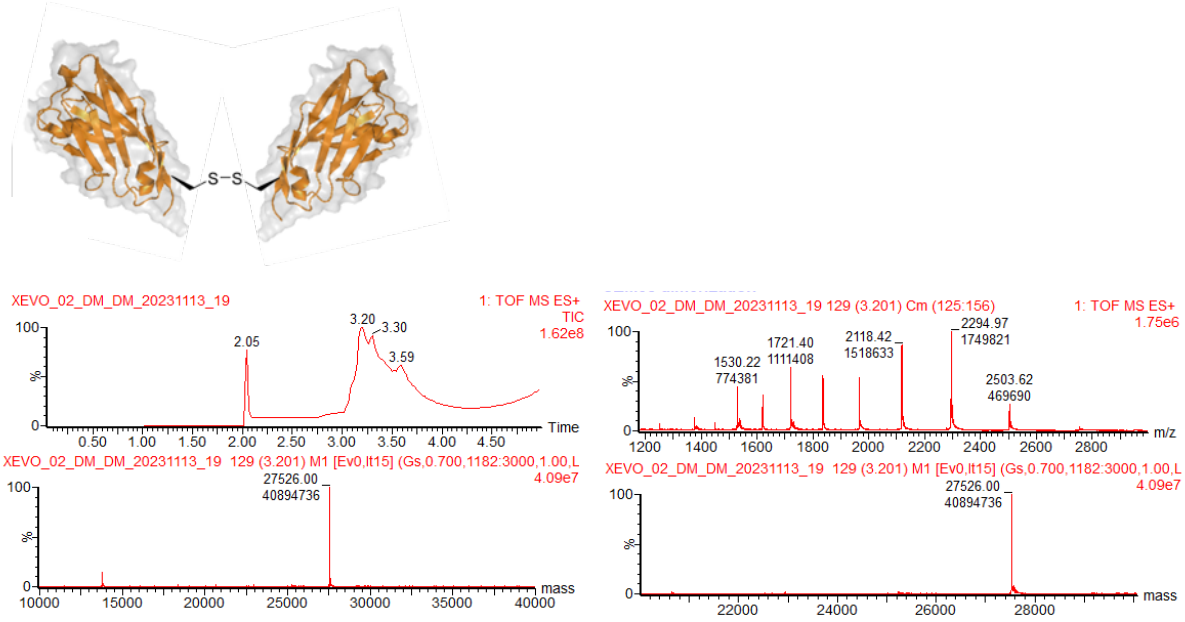


Figure 14. Calculated mass of Gb5-006 homoDiGb: 27524, Observed mass: 27526

#### GbH12 homoDiGb

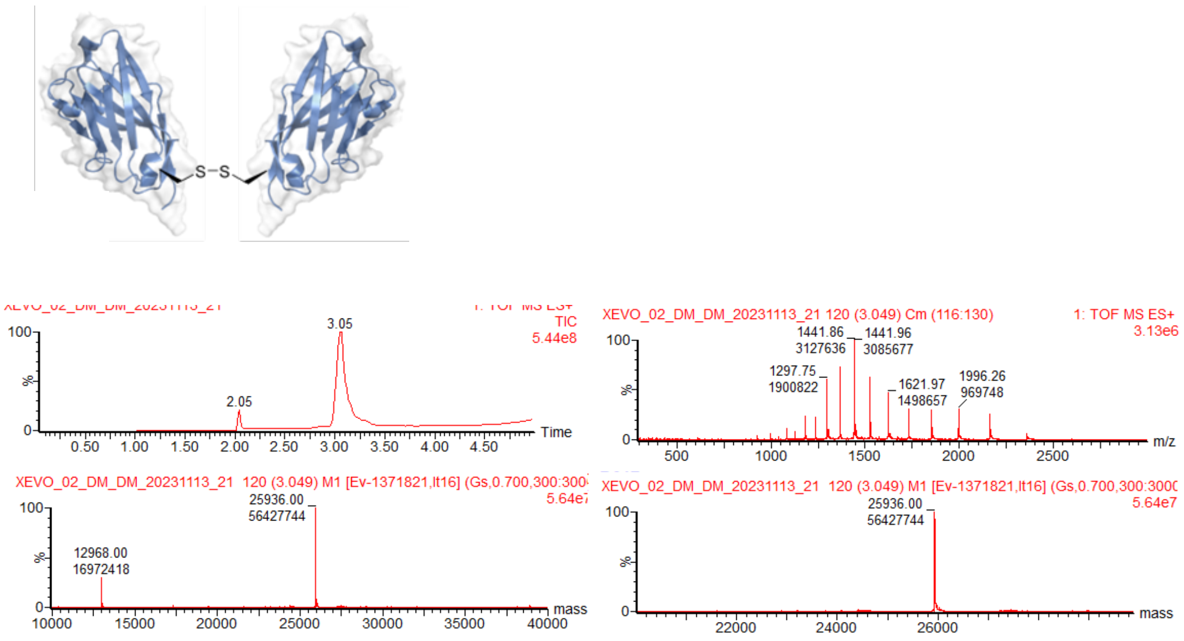


Figure 15. . Calculated mass of GbH12 homoDiGb: 25936, Observed mass: 25936

#### GbEnhancer homoDiGb

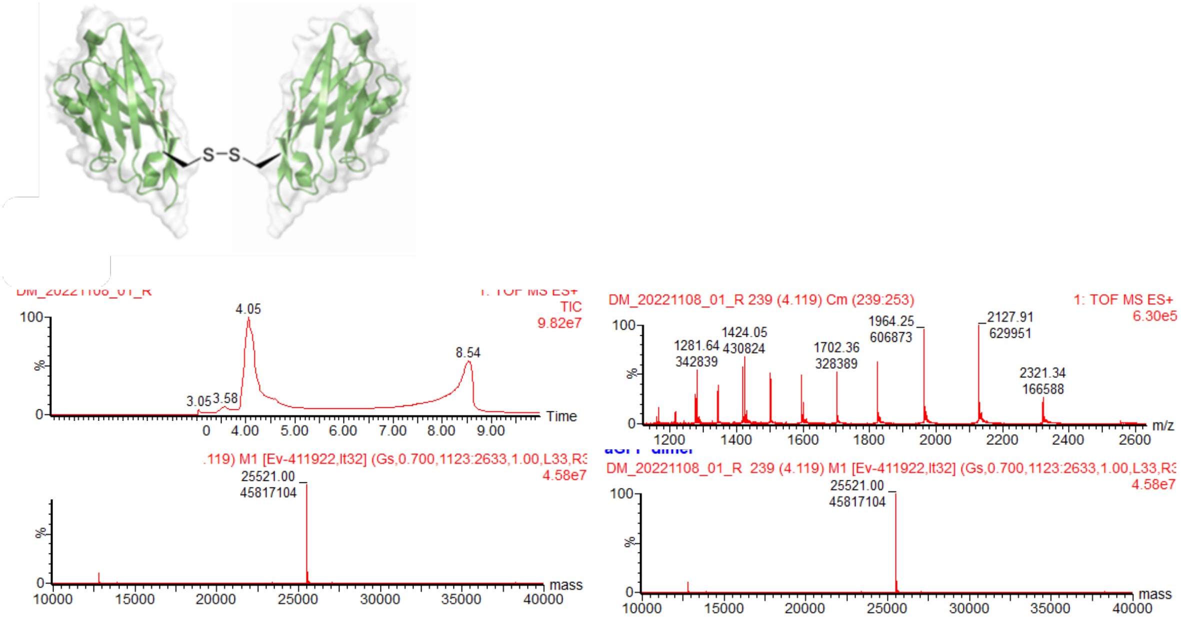


Figure 16. Calculated mass of GbEnhancer homoDiGb: 25520, Observed mass: 25521

#### GbD12 homoDiGb

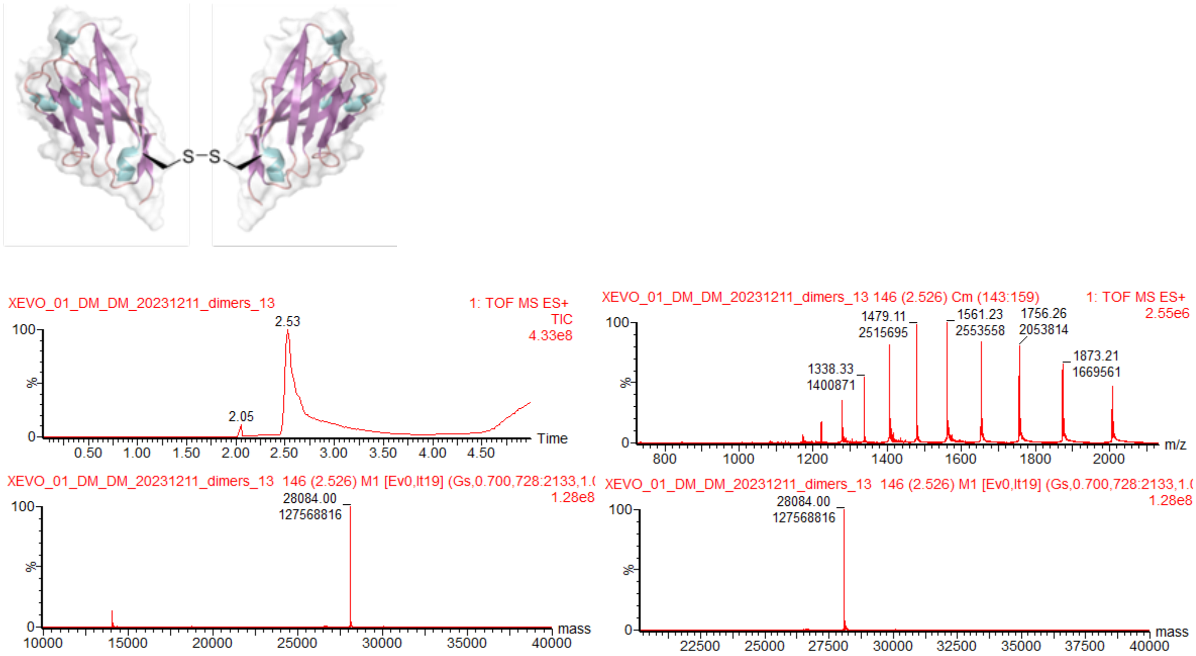


Figure 17. Calculated mass of GbD12 homoDiGb: 28084, Observed mass: 28084

#### GbMBP homoDiGb

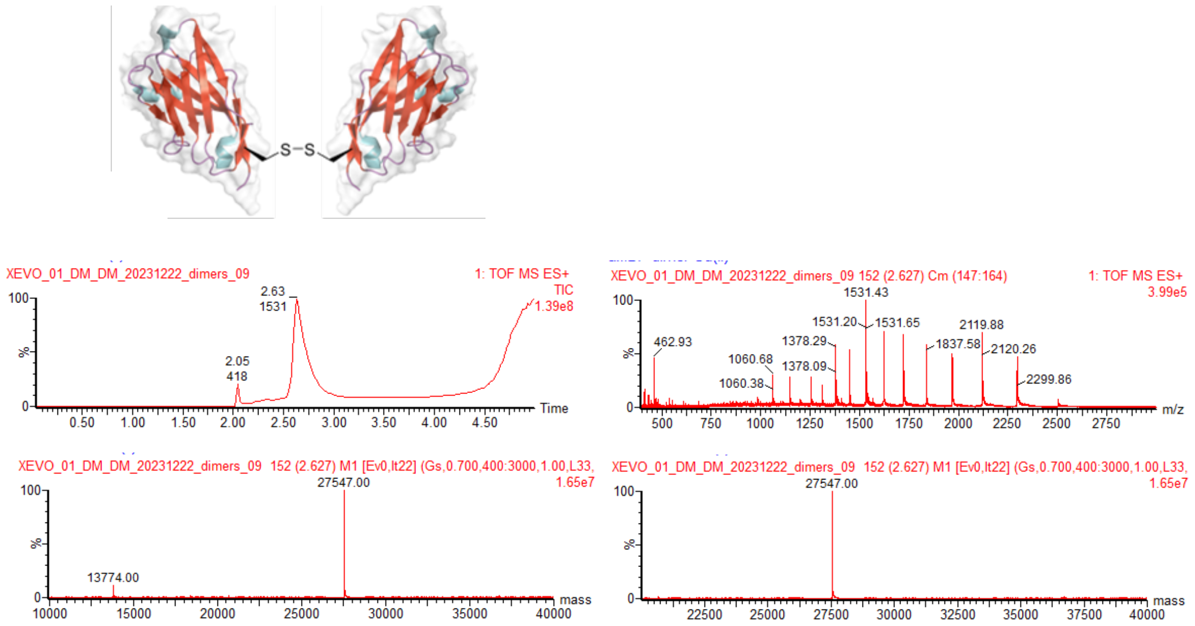


Figure 18. Calculated mass of GbMBP homoDiGb: 27546, Observed mass: 27547

#### GbRBD-1 homoDiGb

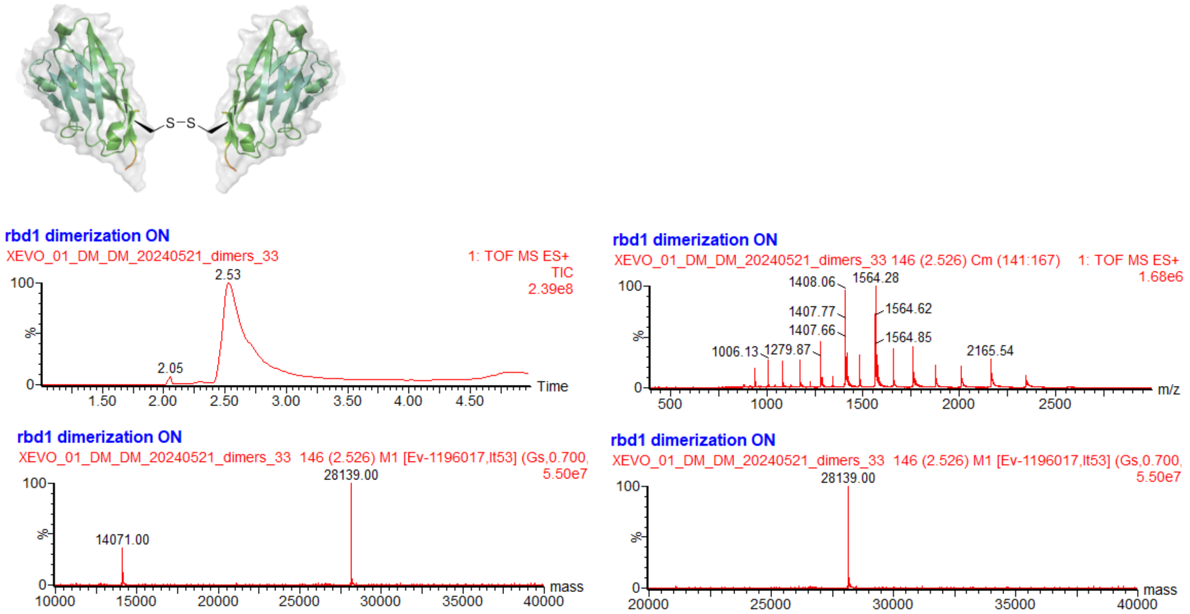


Figure 19. Calculated mass of GbRBD-1  homoDiGb: 28140, Observed mass: 28139

#### GbRBD3 homoDiGb

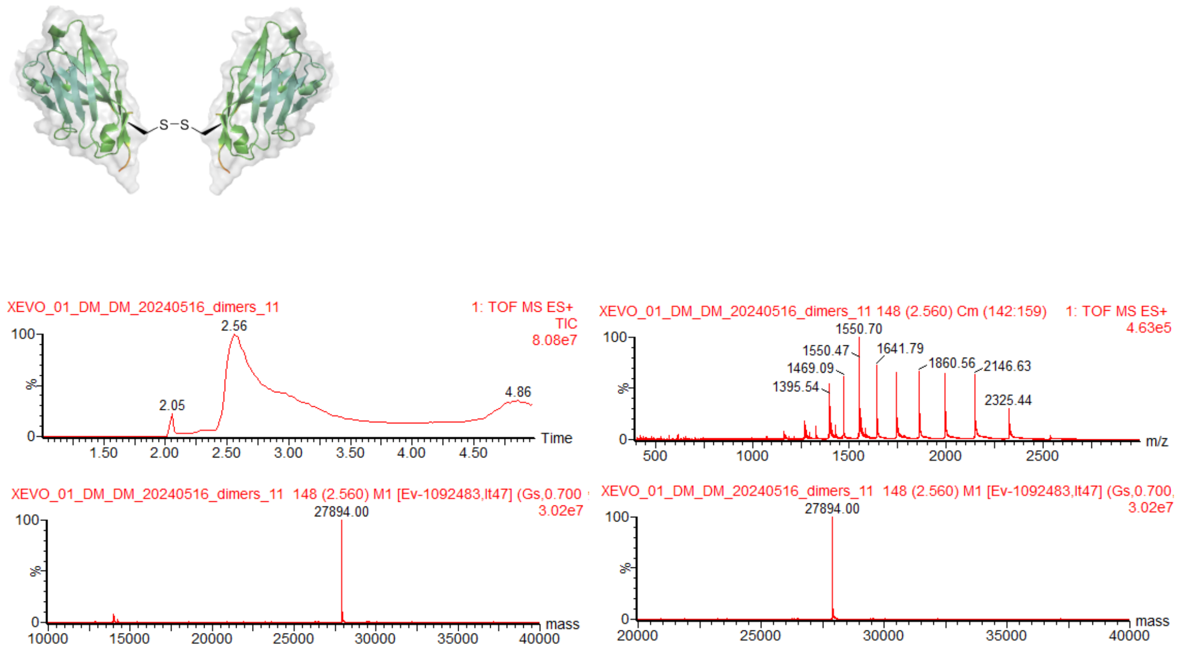


Figure 20. Calculated mass of GbRBD-3 homoDiGb: 27894, Observed mass: 27894

#### GbRBD-6 homoDiGb

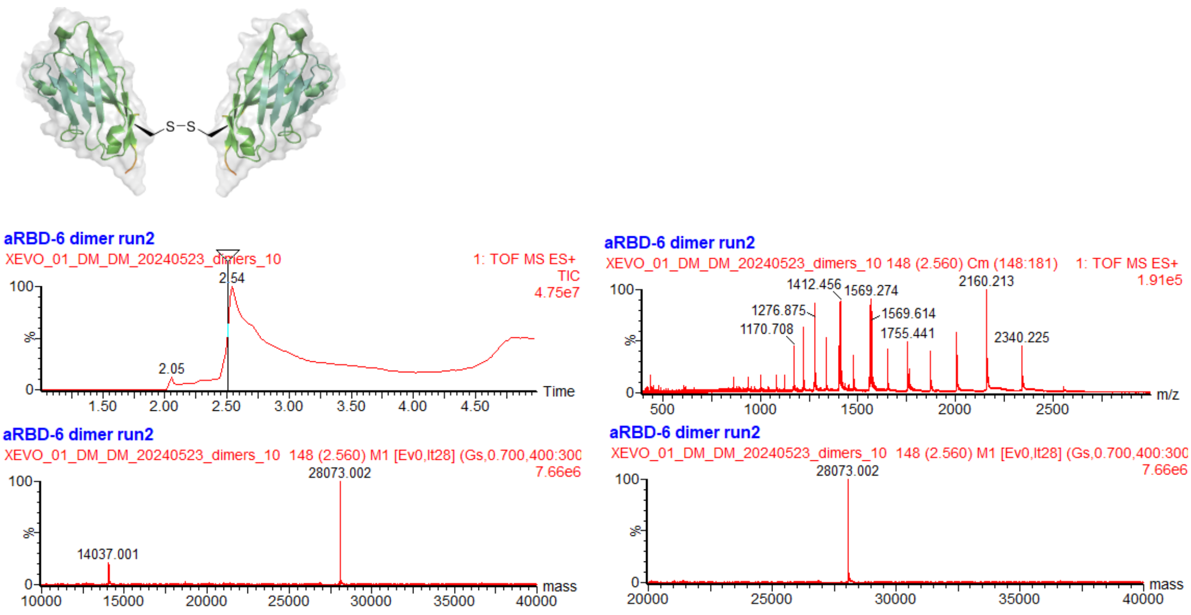


Figure 21. Calculated mass of GbRBD-6 homoDiGb: 28072, Observed mass: 28073

#### GbLys homoDiGb

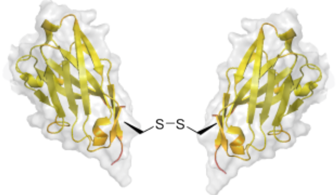


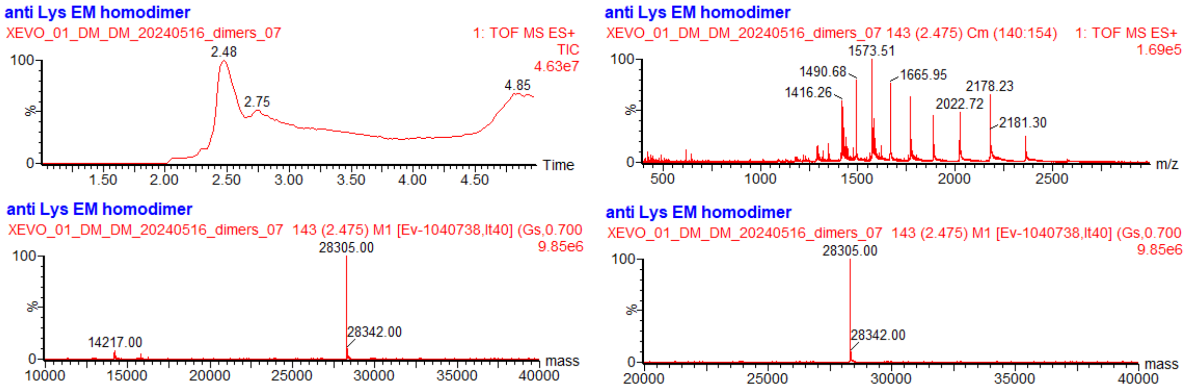


Figure 22. Calculated mass of GbLys homoDiGb: 28306, Observed mass: 28305

#### GbHIV homoDiGb

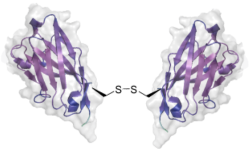


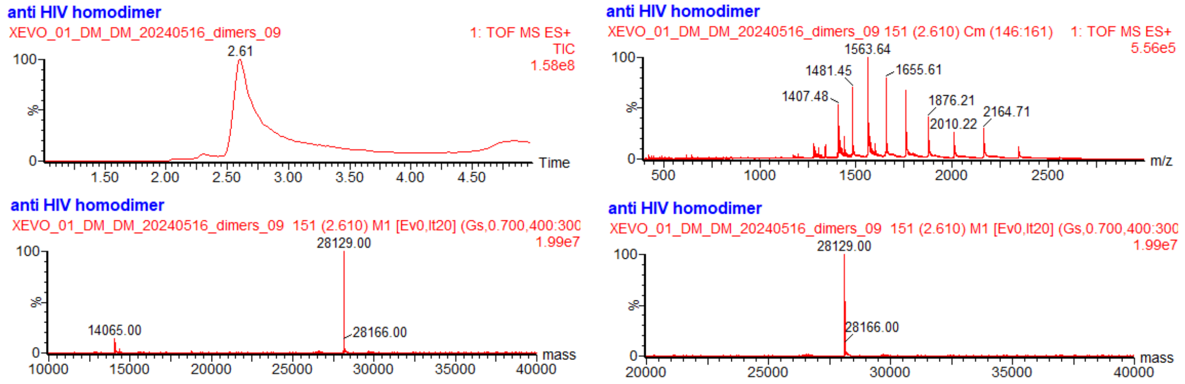


Figure 23. Calculated mass of GbHIV homoDiGb: 28130, Observed mass: 28129

#### General protocol for oxidative relay functionalization to generate heteroDiGbs

Gluebody (GluebodyA) at 1.0 mg/mL in 50 mM $Na_2HPO_4$ (pH 8.0) was treated with 10 eq of DTT at room temperature. After 15 min, DTT was removed from GluebodyA using a PD MiniTrap G25 column equilibrated in $Na_2HPO_4$ 50 mM, pH 8.0. GluebodyA was treated with 10 eq. of 5,5'dithiobis2-nitrobenzoic acid (DTNB) at rt. Conversion was monitored by intact protein mass spectrometry. After 30 minutes, DTNB was removed by buffer exchange using G-25 columns in 50 mM $Na_2HPO_4$, pH 8 and concentrated with Ultra-0.5 3 kDa centrifugal filters.

Next, GluebodyB at 1.0 mg/ml was reduced with DTT using the same protocol as for GluebodyA, desalted using minitrap G-25 columns, and then mixed with activated GluebodyACys12TNB solution from above at 2 x the molar concentration and incubated for 30 minutes at 37 °C, resulting in an indicative yellow-ish solution from displacement of TNB. Conversion was monitored by intact mass spectrometry. Small molecule byproducts were removed from the resulting mixture using a G-25 column in a buffer containing 50 mM $Na_2HPO_4$ (pH 8.0), and the purified heterDiGb flash frozen in liquid nitrogen and kept at –80 °C until further use.

##### Gb5-006-TNB

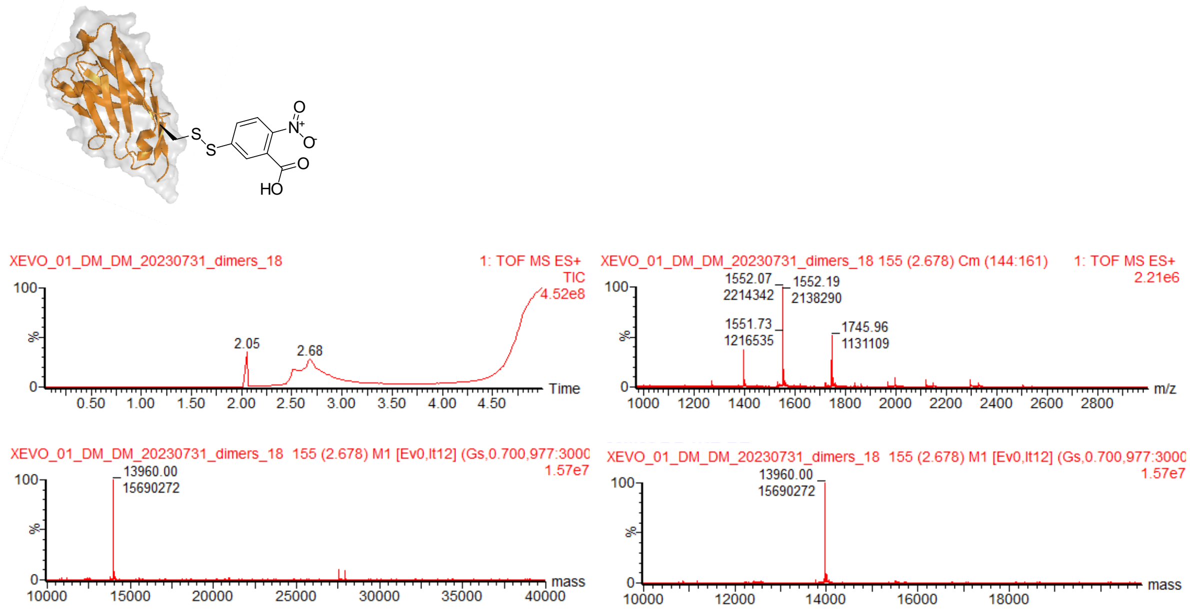


Figure 24. Calculated mass of Gb5-006-TNB: 13958, Observed mass: 13960

**GbEnhancer-TNB**

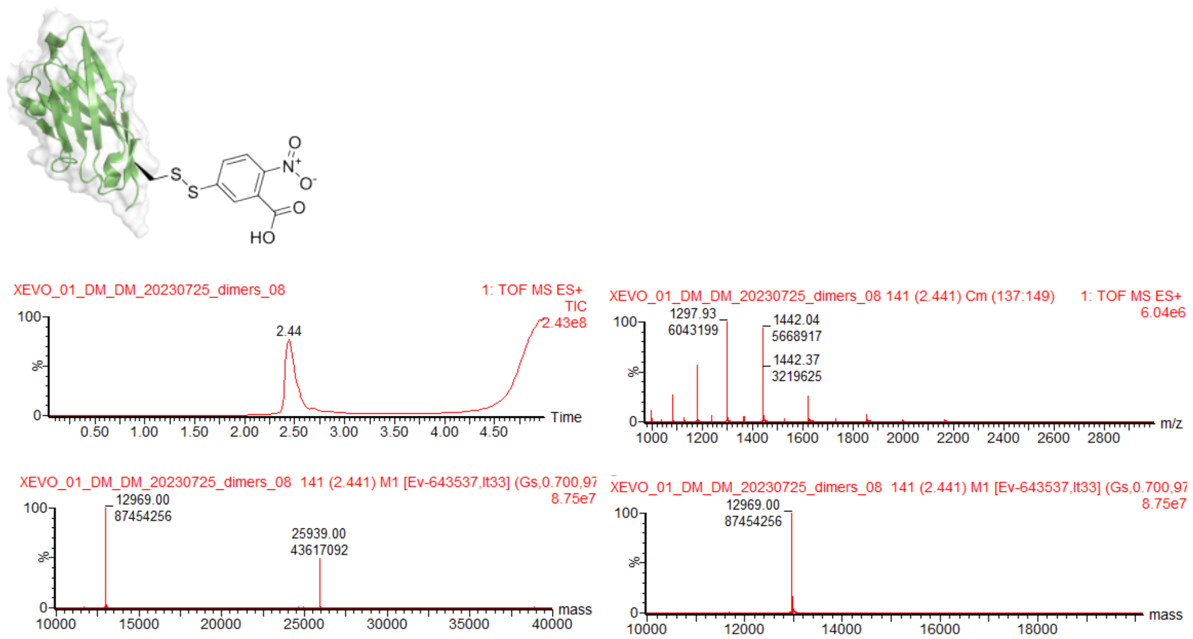


Figure 25. Calculated mass of GbEnhancer-TNB: 12958, Observed mass: 12969

**GbH12-TNB**

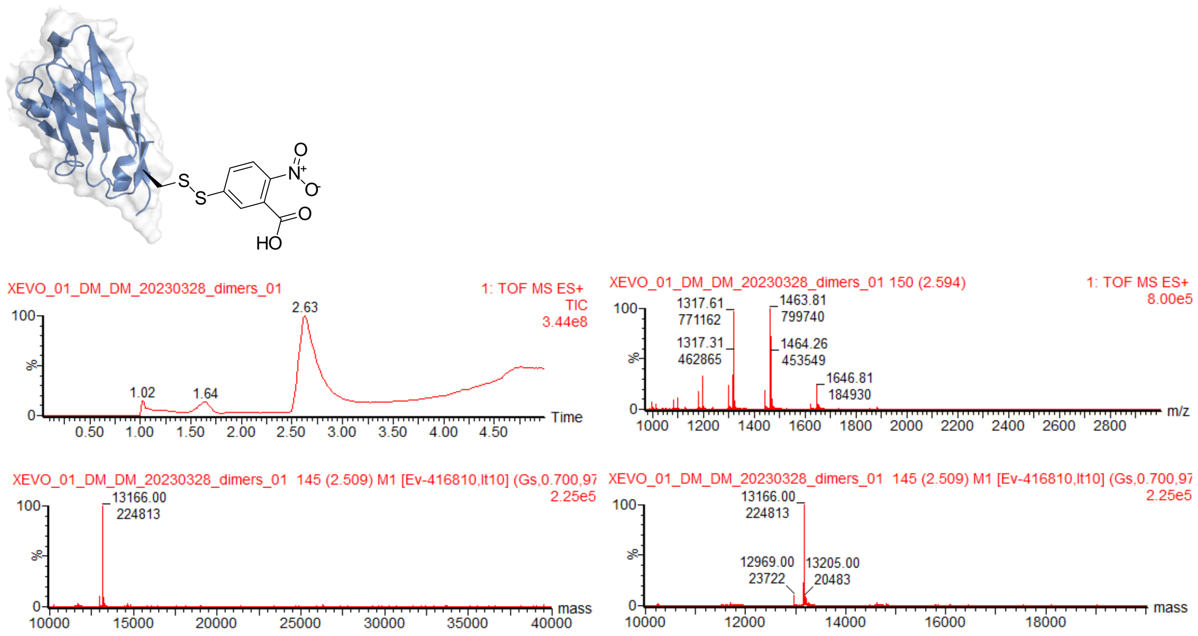


Figure 26. Calculated mass of GbH12-TNB: 13166, Observed mass: 13166

#### HeteroDiGb Gb5-006:GbEnhancer

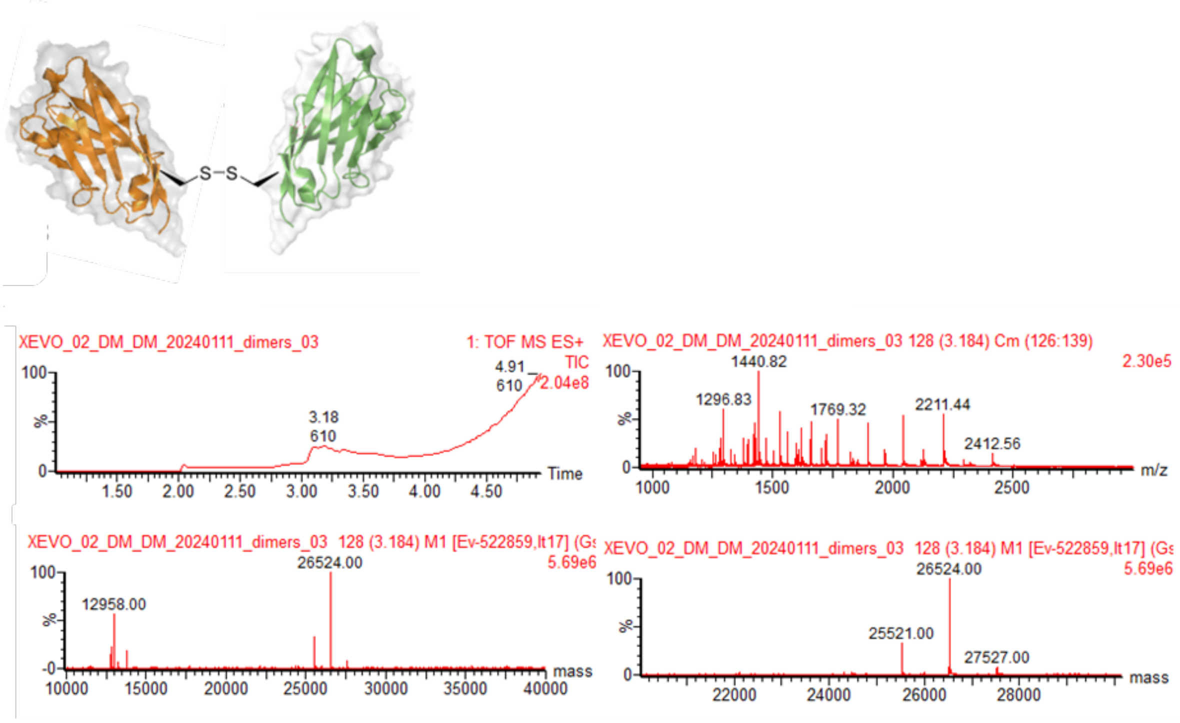


Figure 27. Calculated mass of HeteroDiGb Gb5-006:GbEnhancer: 26522, Observed mass: 26524. Peak with mass 25521 corresponds to GbEnhancer homoDiGb

#### heteroDiGb (anti-SPNS2) GbC4:GbEnhancer

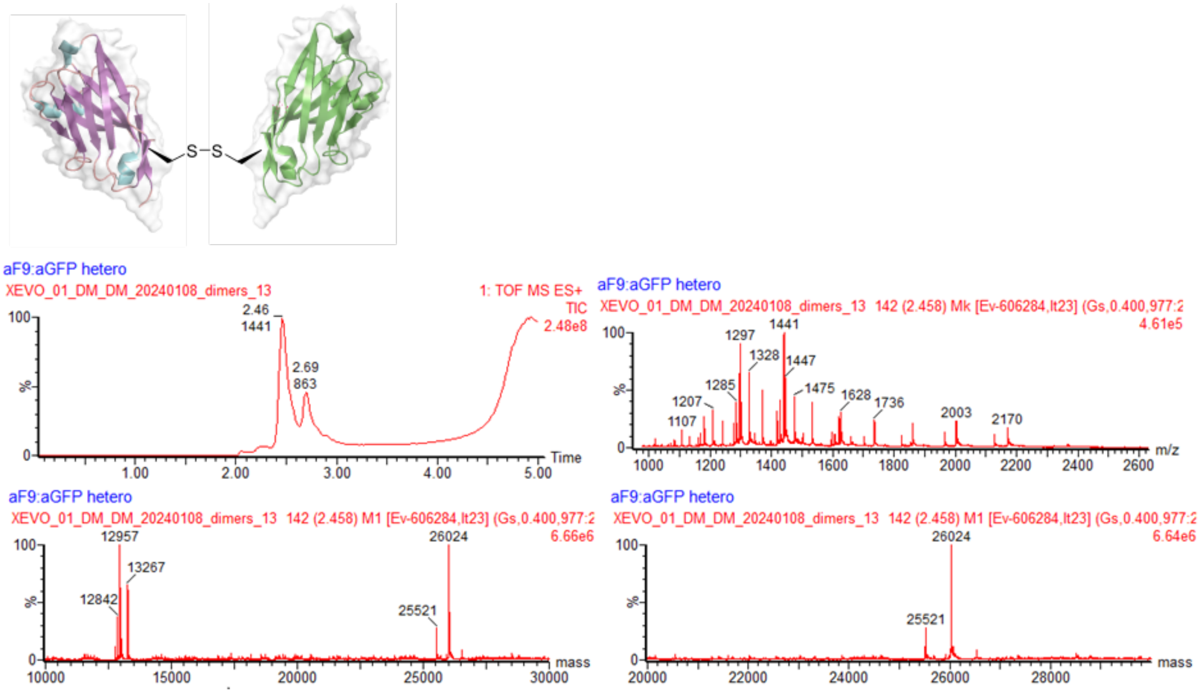


Figure 28.Calculated mass of HeteroDiGb GbEnhancer(anti-SPNS2) GbC4: 26026, Observed mass: 26024. Peak with mass 25521 corresponds to GbEnhancer homoDiGb

**HeteroDiGb GbEnhancer:GbH12**

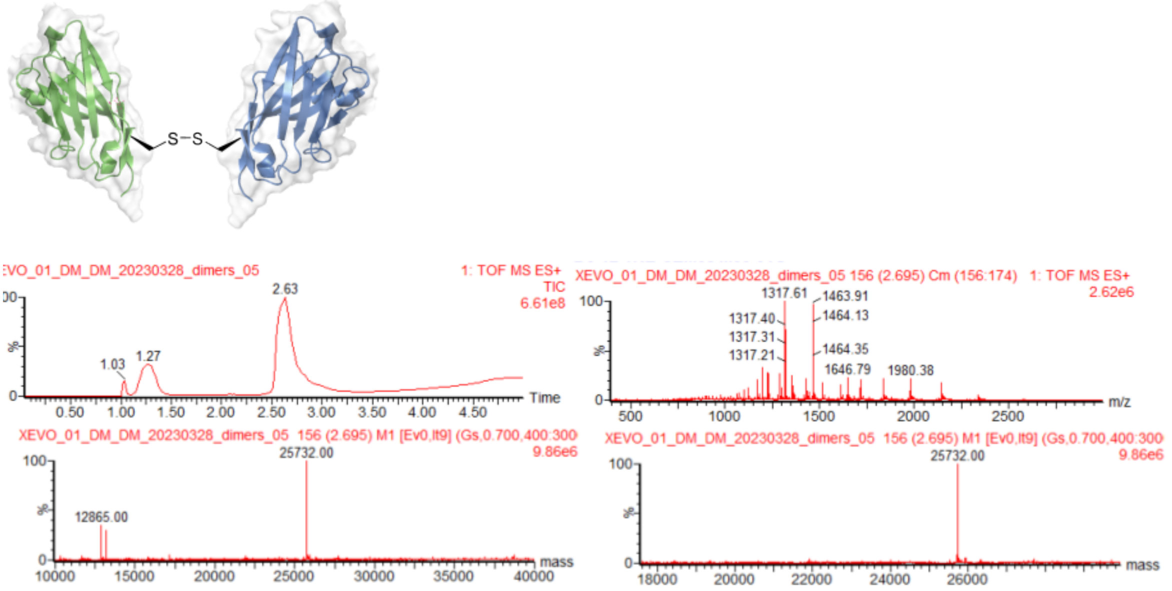


Figure 29. Calculated mass of HeteroDiGb GbEnhancer:GbH12: 25728, Observed mass: 25732

**HeteroDiGb GbEnhancer:GbRBD6**

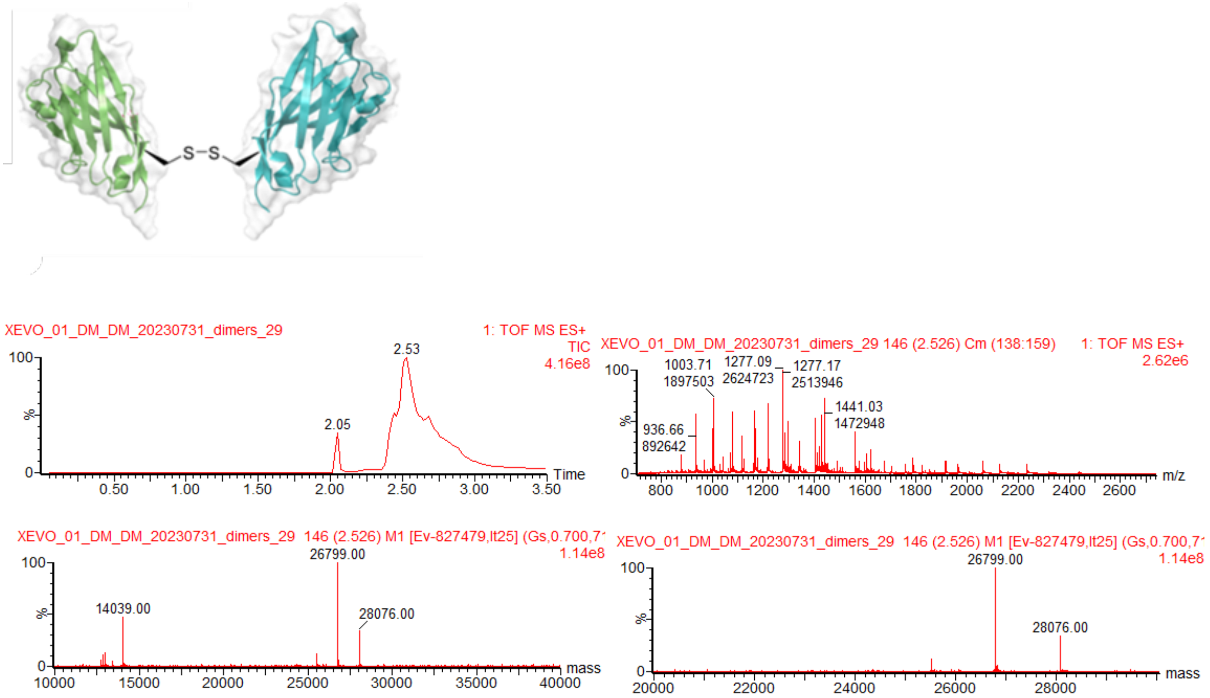


Figure 30. Calculated mass of HeteroDiGb GbEnhancer:GbRBD6: 26796, Observed mass: 26799. Peak with mass 28076 corresponds to anti-RBD6 homodimer

**HeteroDiGb GbEnhancer:GbRBD1**

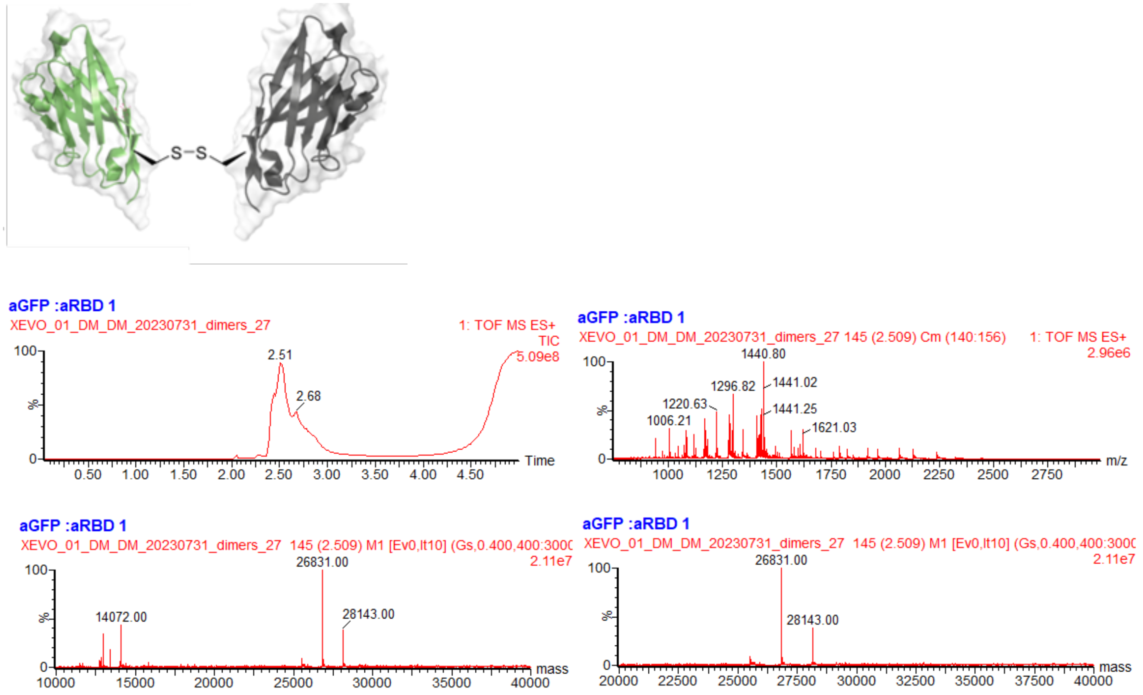


Figure 31. Calculated mass of HeteroDiGb GbEnhancer:GbRBD1: 26830, Observed mass: 26831. Peak with mass 25521 corresponds to GbEnhancer homoDiGb

### Supplementary Discussion

Balanced rigidity with flexibility at the Di-Gluebody interface proved key to ensuring both covalent dimerization and sufficient rigidity of complexation required for cryo-EM and subsequent analyses. A four-atom-bridge between peptide backbones proved optimal: longer linkers caused too much flexibility, whilst those shorter than four atoms proved to be inefficient in their formation and led to less tractable structural analyses. Thus, while the lanthionine three-atom -$CH_2$–S–$CH_2$- linkage was considered, the rigid interface did not allow the ready formation of this 'contracted' bond homologue[1].

Notably, by using four-atom-bridge -$CH_2$–S–S–$CH_2$- linkage, the global intra Di-Gluebody angle was partially varied between 71.3° and 100.7° for the different structures (Figures 2h, Extended Data Table 2) with distinct sets of interacting residues (Extended Data Table 2). At the same time, 3D variability analyses also revealed that within a single complex the intra Di-Gluebody angle varied by at most 8.1° (Extended Data Figure 12 and Extended Data Table 2). Together these results suggest that Di-Gluebody interfaces of homomeric and heteromeric complexes settle into discrete local energy minima, thereby yielding a structurally homogeneous population. Together these data show sufficient flexibility for reaction to form the covalent linkage yet sufficient rigidity to allow structural solution. This 'ideal balance' seemingly explains the ready application that we show here.
